# Life history and infection susceptibility parameters of *Drosophila* species reared on a common diet

**DOI:** 10.1101/2025.11.25.690557

**Authors:** Hongbo Sun, Millie Cole, Eduardo Gomes-Ragnoni, Jodi Johnson, Alexander Jarvis, Léna Hédelin, Hannah E. Westlake, Ben Longdon, Mark A. Hanson

## Abstract

The fruit fly *Drosophila* is a powerful model organism for many biological questions. While *Drosophila melanogaster* has been the most widely used species owing to its versatile genetic tools, comparative studies also take advantage of the rich evolutionary, ecological, and behavioural diversity of Drosophilidae. This model system has only benefitted from the sequencing of hundreds of genomes assembled to chromosome level, and rich array of publicly-available datasets. However, labs that typically use just *D. melanogaster* can have difficulties adopting other species into their experiments due to unique dietary needs and care regimens. While dedicated research groups can rear dozens of fly species, this often requires preparing a variety of food media, which can be logistically taxing. In this study, we report an agar food recipe that we have used to rear >70 species of Drosophilidae (Prop recipe). This list includes ecological specialists such as *D. sechellia*, major pest species including *D. suzukii*, cactophiles, slime flux feeders, detritivores, and even mushroom-feeding species of the Quinaria and Testacea groups. This food recipe uses only standard powdered ingredients and molasses, and is already in use by one commercial transgenic injection provider. We provide life history parameters for dozens of species reared on this recipe, including comparisons in a subset of species across a variety of diets, finding fly fitness to be comparable or better on the Prop recipe among tested species. We further provide comparative data for 42 species after systemic infection by *Acetobacter* bacteria. However, unexpectedly, we found a striking interaction of diet and the microbiota that impacts susceptibility to infection in species-specific fashion. Our results suggest certain experiment types may be impacted by diet due to the microbiome that the diet itself supports. By providing rearing advice, life history parameters, and this infection dataset and experience, we hope to enable researchers to take advantage of the diversity of *Drosophila* as a model system for evolutionary research.

## Introduction

*Drosophila* fruit flies are used in a wide range of biomedical, ecological, and evolutionary research fields. The powerful tools of *Drosophila melanogaster* have led to numerous groundbreaking discoveries in genetics, developmental biology, immunology, and physiology [1–3]. With the recent advances in CRISPR/Cas gene editing, transgenic tools development is now widely available across invertebrates [4–6]. This includes other species of *Drosophila* with unique characteristics for behavioural, ecological, and evolutionary research [7–12]. While *D. melanogaster* will continue to pioneer genetic advances, understanding the origin of various biological processes will benefit from incorporating the wider diversity of *Drosophila* species into future research [13].

The rearing of *D. melanogaster* has been practiced for over a century and generated its own global industry [14]. These practices have made for the efficient adoption of *D. melanogaster* as a research tool, and a high reproducibility of *Drosophila* research [15]. However, the procedures for rearing *D. melanogaster* are not suited to many drosophilid species of ecological, evolutionary, or economic interest. Alternate species often require specialised food preparations, supplementing *D. melanogaster* food media with specialty fruit extract (e.g., *D. sechellia*), mushroom (e.g. *D. testacea*), or plant matter (e.g., *Scaptomyza* species) [16]. Even if species are viable on a given diet, dietary nutrition is known to play key roles in fly fitness [17–19], making it difficult to perform among-species experimentation without potential confounding effects of species-specific diets. As a consequence, many research groups artificially restrict the diversity of species they study by only keeping flies that can be reared on their local food media. Thus, logistical constraints artificially restrict the utility of the *Drosophila* genus for evolutionary studies.

Here, we describe a *Drosophila* agar food recipe that rears flies from the full breadth of the genus *Drosophila*. This recipe can be prepared without any specialised equipment or procedures, and uses only ingredients that do not require refrigeration. We provide life history and fitness parameters for 66 species from lineages of *Drosophila, Scaptodrosophila, Zaprionus, Dorsilopha*, and *Hirtodrosophila* reared on this food recipe. We further compare life history parameters of select species on more traditional *Drosophila* food preparations, and a comparative infection study across 42 species that inspired us to characterise microbiota differences depending on diet. By providing data on a litany of fly species reared on this food recipe, we hope to make adopting alternative *Drosophila* species in biological research more approachable.

## Results

### Conceptualisation of this study and the Prop recipe

Traditional approaches to rearing fly species in research labs has largely been left to trial and error. In our previous studies, we have used various food recipes to rear dozens of fly species. However, revisiting rearing conditions in our lab, we realised most phylogenetic clades had species that were happily reared on the Cambridge Propionic food recipe. This led to a gradual testing of this food recipe across all species. In May 2024, our entire species collection was transferred to a slightly modified version of the Cambridge Propionic recipe (hereafter “Prop recipe”), and we have been rearing all fly species on this single recipe for over one year with no abnormal challenges.

The Prop recipe we use is given in **Box 1**, including product specifics to allow better reproducibility of our food preparations. The Prop recipe food has a good viability when stored in the refrigerator for up to 2 weeks without issue, although fresher food is always preferable. Longer refrigeration times are possible, but may encourage dessication or microbial growth.

### Rearing practices and advice

While the Prop recipe can rear all species in our lab, some perform much better with simple additional supports. We rear fly species in both fly bottles and standard 25/28.5mm *Drosophila* vials. Below we outline considerations for rearing challenging flies on the Prop recipe.

1. A piece of climbing structure is universally helpful. For bottles, we typically use a ~½ piece of non-absorbant cotton ball as a go-to simple climbing structure (**Figure 1**). One advantage of this approach is the dense fibres on the cotton ball stick to the food media, preventing it from falling into the next bottle.
2. For vials, we insert a piece of dental cotton roll into the food as a climbing structure and to stabilise vial humidity (**Figure 1**). Paper strips can also be used. If local humidity causes food to pull away from the vial, adding water or 0.5% propionic acid to active vials can prevent larval/pupal dessication.
3. In our hands, the microbiota on the Prop recipe is rich. A high larval density improves rearing success, churning the food and preventing biofilm or mould growth. Flipping repeatedly to reduce microbial loads [20], alongside increased fly density, and so egg-laying density, can improve culture health.
4. Transitioning stocks over to the Prop recipe may require acclimating flies. Carrying over a small amount of food and larvae from the previous vial can help seed the new vial with a familiar microbiota and encourage oviposition.
5. For our regular flipping, we sprinkle dried instant yeast into bottles to encourage oviposition. This does not impede oviposition in specialists, and improves oviposition in various species including Melanogaster group flies.

**Figure 1:**
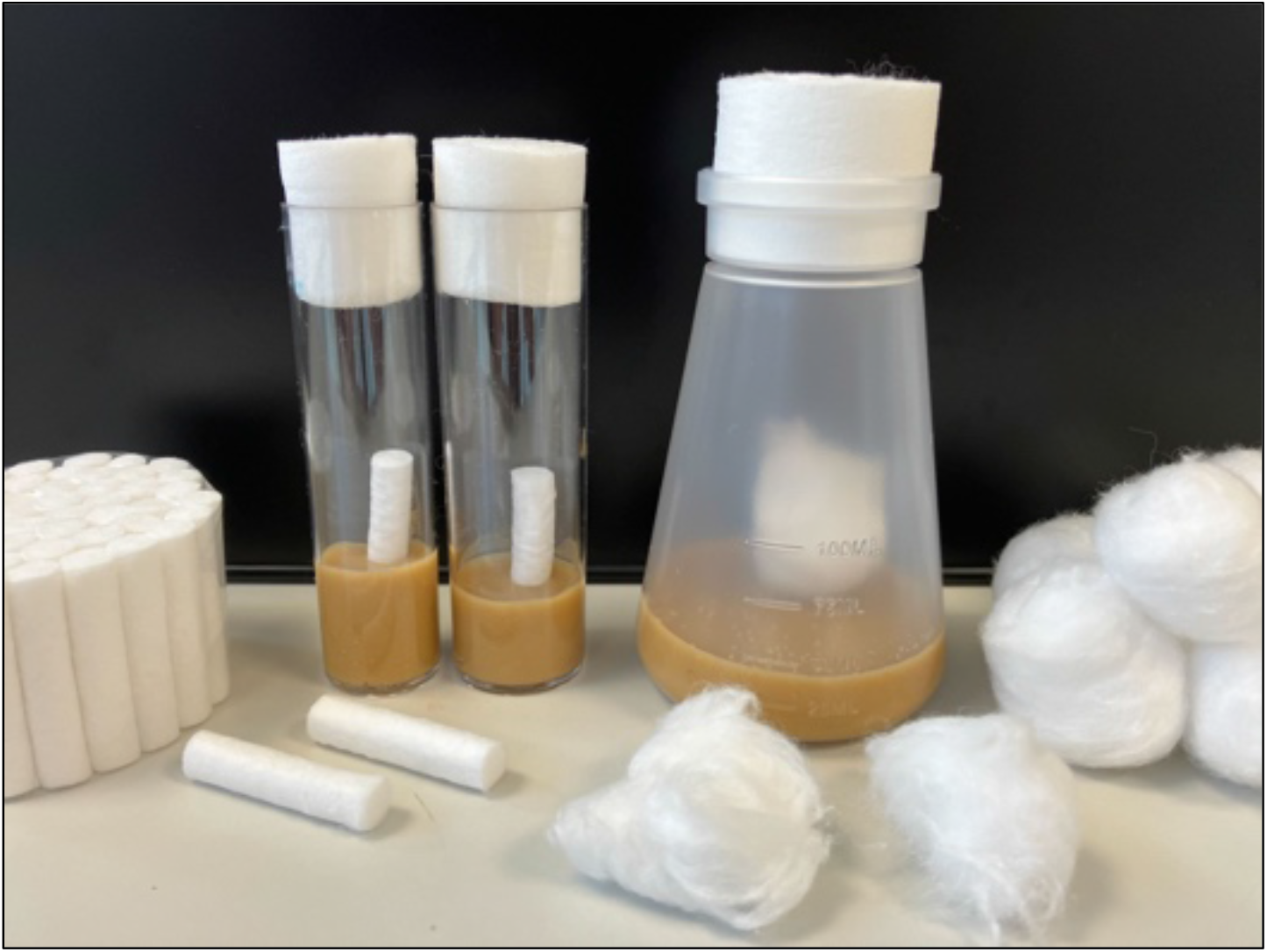
example vials and bottle containing cotton dental wick or a ½ piece of non-absorbent cotton ball. Vials and bottles can additionally contain sprinkled yeast to encourage oviposition and help regulate the microbiota.

**Box 1.** Prop recipe, per 1 Litre.

|  |  |
| --- | --- |
| • 1000mL Water | (MilliQ distilled) |
| • 10g Agar | (Agar Agar (E 406), Intra Labs) |
| • 30g Yeast | (fermipan red, dried instant yeast) |
| • 70g Cornmeal (yellow, coarse) | (Hunt's Food Group, UK) |
| • 10g Soya flour | (Hunt's Food Group, UK) |
| • 80g Spray dried malt extract | (Munton's Spray Malt Light) |
| • 22g Molasses | (NAF Value Molasses) |
| • 14mL 10% Methylparaben in EtOH | (Sigma, Cat. no: H3647) |
| • 6.2mL Propionic acid | (Fisher, Cat. no: 10634622) |

*Brief instructions: combine all ingredients except methylparaben and propionic acid, stirring constantly. Bring to a rolling boil for ~2 minutes, ensuring the agar is completely melted. Allow to cool until $\leq 70^{\circ}\text{C}$ , then add methylparaben and propionic acid; methylparaben and propionic acid act as mould inhibitor and can be deactivated if added at higher temperatures. Dispense into bottles/vials.*

## Fly species that can be reared on the Prop recipe

In total, we have reared >70 species of *Drosophila* on the Prop recipe including species from the Melanogaster, Obscura, Willistoni, Saltans, Virilis, Repleta, Melanica, Dorsilopha, Quinaria, Testacea, Bizonata, Immigrans, *Zaprionus, Scaptomyza, Scaptodrosophila*, and *Hirtodrosophila* species groups and subgenera. Some of these lineages are more fecund than others, but all of them are capable of being maintained stably for over one year. Below we provide descriptive statistics for host fitness on the Prop recipe.

### Egg to adult development time

We collected egg to adult development times for 66 unique species reared on the Prop recipe at 22°C at a relative humidity of ~50%. Egg to adult development time was assessed as the time since the vial was made to the date of first offspring emergence, summarised in **Figure 2A** and **Table 1**.

**Table 1:**
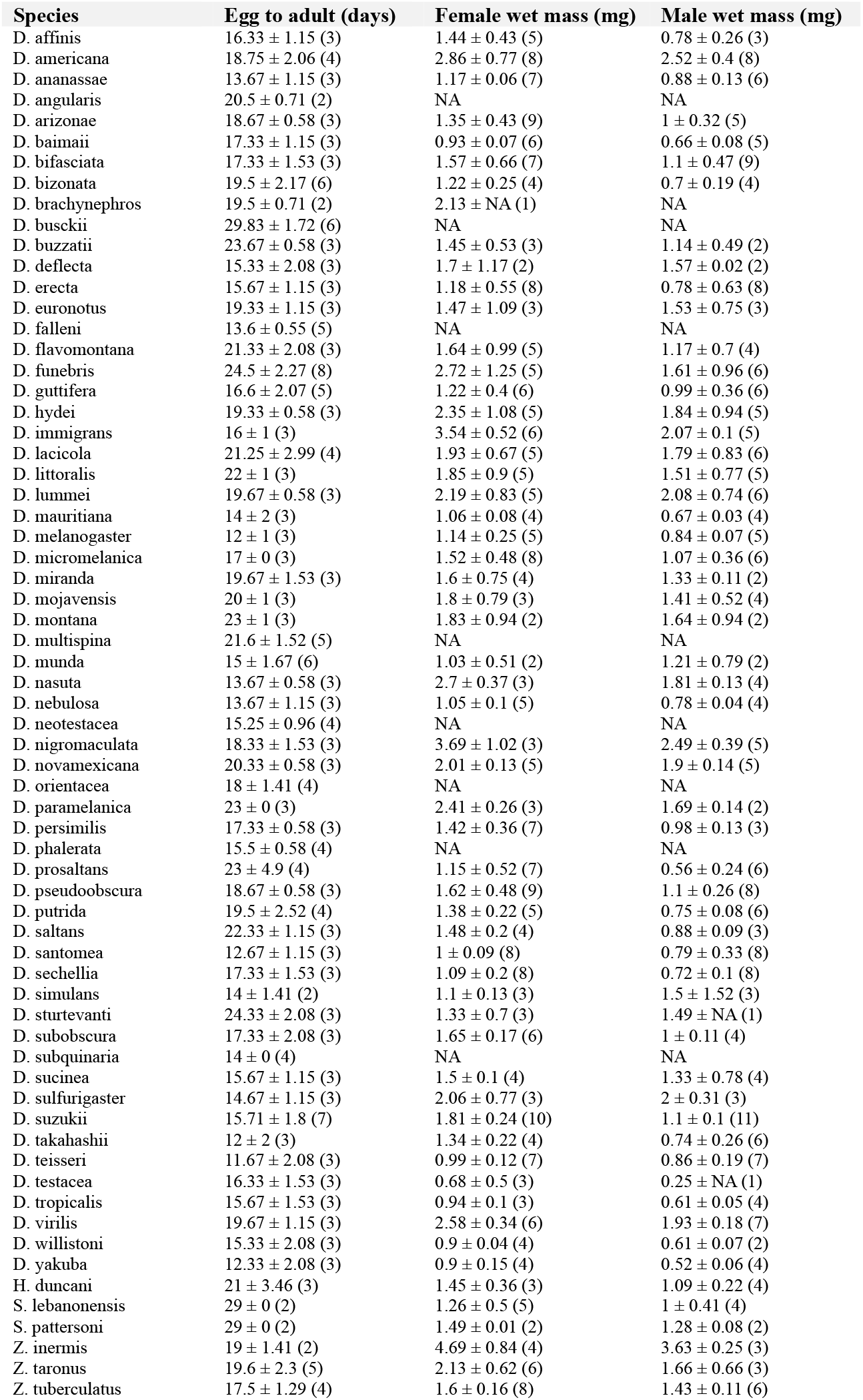
data underlying Figure 1. Data are reported as mean ± standard deviation. Replicate counts given in parentheses. NA = no data.

**Figure 2:**
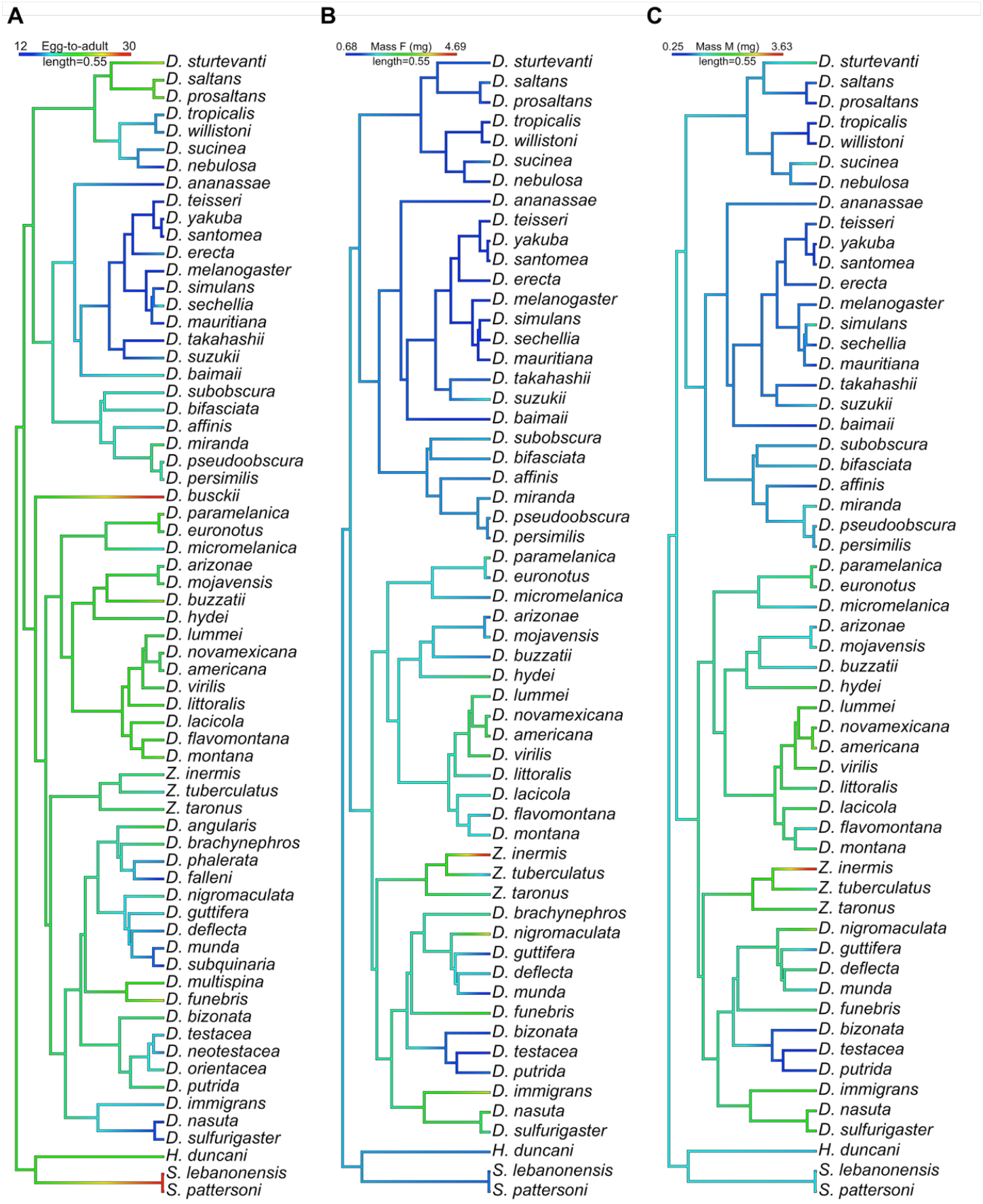
individual parameters of fly species reared on the Prop recipe. **A)** Egg-to-adult times have a very high phylogenetic heritability signal 0.98 (HPD interval: 0.93, 1.00). **B-C)** Mean wet mass of females **(B**, mean = 0.97, HDP = 0.82,1.00) and males **(C**, mean = 0.97, HDP = 0.82, 1.00). Data were collected from up to 66 species. Precise values are reported in Table 1. Correlation of male-female wet mass ratios is given in **Figure 2supp1**.

The maximum egg-to-adult development time observed was *D. busckii* at 29.8 ± 1.7 days (mean ± standard deviation) followed closely by the *Scaptodrosophila* species *S. lebanonensis* and *S. pattersoni* (29.0 ± 0.0), and the minimum was *D. melanogaster* at 12.0 ± 1.0 days, similar to the 11-12 at 22°C reported by others [21].

We performed a phylogenetic mixed model approach (MCMCglmm) including species as a random factor to estimate the phylogenetic heritability (*σ*^*2*^_*p*_*/(σ*^*2*^_*p*_*+ σ*^*2*^_*s*_) of egg-to-adult development time across *Drosophila* species (see [9]). The proportion of variance explained by the host phylogeny was 0.98 (HPD interval: 0.93 – 1.00). Thus, agreeing with heritability of developmental traits in other organisms [22,23], there is a very high phylogenetic heritability for egg-to-adult development times within *Drosophila* species separated by ~53 million years of evolution [24].

### Body mass of flies reared on the Prop recipe

Lesperance and Broderick [25] highlight that “standard” *Drosophila* food media differ markedly in their protein-to-carbohydrate ratio (P:C ratio) and nutritional value. Sannino and Dobson [26] also note that local suppliers can vary in their preparation of ingredients, even including over successive batches of the same product, with dramatic effects on lifespan. In *D*. melanogaster, male and female nutritional requirements also impose different nutritional optima for each sex, suggesting sex-by-diet interactions [17].

We therefore collected data on wet mass of males and females reared on the Prop recipe (P:C ratio ≈ 0.15) alongside egg-to-adult experiments. The proportion of variance explained by the host phylogeny for males and females was an identical 0.97 (HPD interval: 0.82 – 1.00), distinguished only if rounding to the third decimal place (Fig. 2B,C).

### Comparison of the Prop recipe to other standard Drosophila diets

We estimate our adaptation of the Prop recipe has a P:C ratio of approximately ~0.15 using the Drosophila Dietary Composition Calculator [25]. Food recipes in the literature can have P:C ratios anywhere from 0.07 to 0.33, specifically the Cambridge [9] and EPFL [27] cornmeal recipes, respectively. In initial experiments, we observed that the Prop recipe reared most species with shorter egg-to-adult times than two cornmeal recipes with P:C ratios of 0.07 (*P* = 0.03, Cambridge cornmeal) and 0.15 (**Figure 3supp1**), though the latter comparison was not significant (*P =* 0.26, a recipe we call “Penryn cornmeal”).

**Figure 3:**
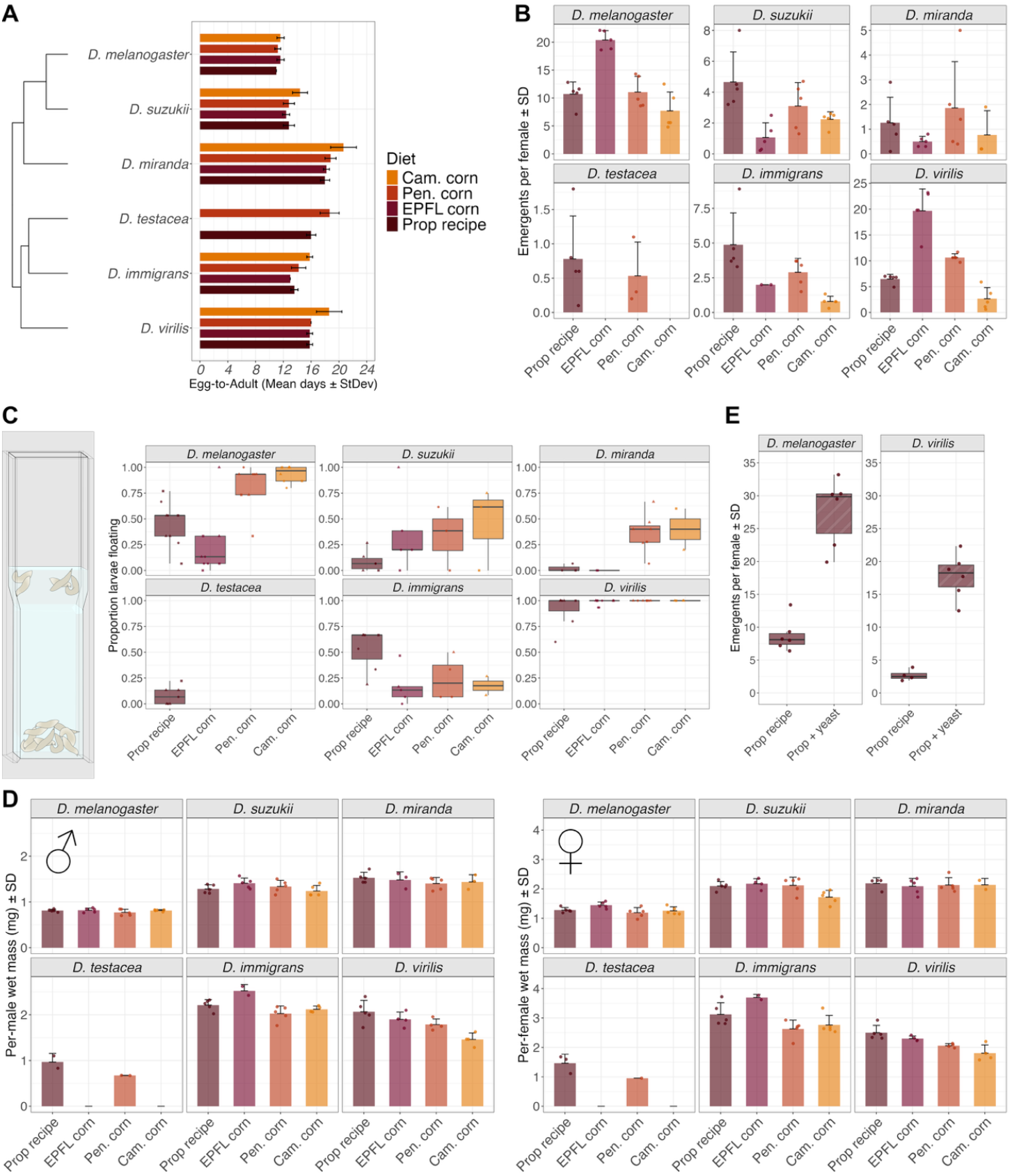
comparisons of six species on four food types. **A)** Egg-to-adult times from vials from a 24h egg-laying period. **B)** Total emergents per female by species and food type from a 24h egg-laying period. **C)** Larval buoyancy rates (example illustration provided) reared on each diet. **D)** Wet mass of up to 25 pooled 7dpe flies by species and diet for males and females. **E)** Total emergents from *D. melanogaster* and *D. virilis* is dramatically improved by the addition of 200mg of dry yeast to vials.

To better situate how rearing on the Prop recipe or other diets affects generic fitness of individuals, we additionally reared six species on four different food recipes spanning the breadth of P:C ratios. These included generalist species amenable to most diets (*D. melanogaster* and *D. virilis*), and also species known to have a range of specialised needs (*D. suzukii, D. miranda, D. immigrans, D. testacea*) – also see materials and methods. We collected egg-to-adult development times (**Figure 3A**), total emergents (**Figure 3B**), larval lipid content using a buoyancy assay (**Figure 3C**) [28], and wet mass of pooled adult emergents from each food type (**Figure 3D**).

Egg-to-adult times were broadly similar for all species when reared on the Prop recipe, EPFL cornmeal, or Penryn cornmeal, but not Cambridge cornmeal (Figure 3A). However, we found a diet-by-species interaction. For *D. melanogaster* and *D. virilis*, the EPFL cornmeal recipe recipe produced the most offspring (linear mixed model: emergents ~ diet*species + (1| block), *P* < .001, Figure 3B). On the other hand, rearing success of *D. suzukii, D. testacea*, and *D. immigrans* was generally worse on Cambridge cornmeal (*P* < .05 for *D. testacea, D. immigrans*) or the EPFL recipe (*P* < .05 for *D. suzukii, D. testacea*), compared to the Prop recipe.

Interestingly, the body mass of *D. melanogaster* and *D. virilis* flies reared on the Prop and EPFL recipes were broadly similar (**Figure 3D**, *P* > .10 for both males and females), suggesting total emergent differences may stem from parent egg-laying behaviour and not nutrient deficiency. Indeed, the emergents-per-female of *D. melanogaster* and *D. virilis* could be improved dramatically by supplementing Prop recipe vials with ~200mg yeast added atop the food (**Figure 3E**). On the other hand, the EPFL cornmeal promoted the emergence of larger flies from *D. suzukii* and *D. immigrans*, significantly so for females compared to Cambridge cornmeal (**Figure 3D**, *P* = .043). Larval buoyancy also showed species-by-diet interactions, where no diet sorted with a consistent rank order across all species (**Figure 3C**). Thus, there are various fitness parameters with species-specific effects on a given diet, and positive trends in one measurement do not necessarily correlate with positive trends in another.

Overall, these results suggest the Prop recipe is uniquely well-placed for difficult-to-rear species, emphasized by *D. suzukii, D. miranda, D. immigrans*, and *D. testacea*, alongside the versatility shown in **Figure 2**. While species such as *D. melanogaster* and *D. virilis* produce more offspring on the high P:C ratio EPFL diet, similar total emergent rates can be accomplished by supplementing the Prop recipe with yeast atop the food. This increased reproductive output could be due to increased oviposition behaviour due to yeast volatiles (as in [29]), though we cannot rule out a protein-dependent nutritional effect given our experiment design.

### The Prop recipe supports a rich microbiota

We found that microbial growth differs visibly on the Prop recipe compared to Cambridge cornmeal. We tested this effect by checking microbial loads of vials that contained flies flipped every 3 days for 9-14 days, which confirmed Prop recipe microbiota counts from 3-day old vials were orders of magnitude higher (**Figure 4A-D**). We observed higher CFU counts on MRS and YPG agar, with colony morphology and growth times consistent with *Acetobacter* and an additional unidentified microbe that was visually ~20x less abundant. We also saw increased CFU counts on LB and BHI agar, media which were not consistent with *Acetobacter* colonies, suggesting the promotion of other microbes as well. In addition, there were some species-by-diet interactions in total CFUs on a given bacterial growth media: for example, *D. immigrans* flipped on the Prop recipe shows higher CFU counts on LB and BHI compared to Cambridge cornmeal (non-*Acetobacter* colonies), as well as higher CFU totals on these bacterial growth media compared to *D. melanogaster* flipped on either fly diet (**Figure 4C,D**). This increased microbiota load observation may be generalisable, or specific to the microbiota present in our lab (Penryn, UK), where a rich diversity of bacterial genera such as *Acetobacter, Commensalibacter, Lactobacillus, Enterococcus, Serratia, Pseudomonas, Morganella*, and *Providencia* species has been found previously [30].

**Figure 4:**
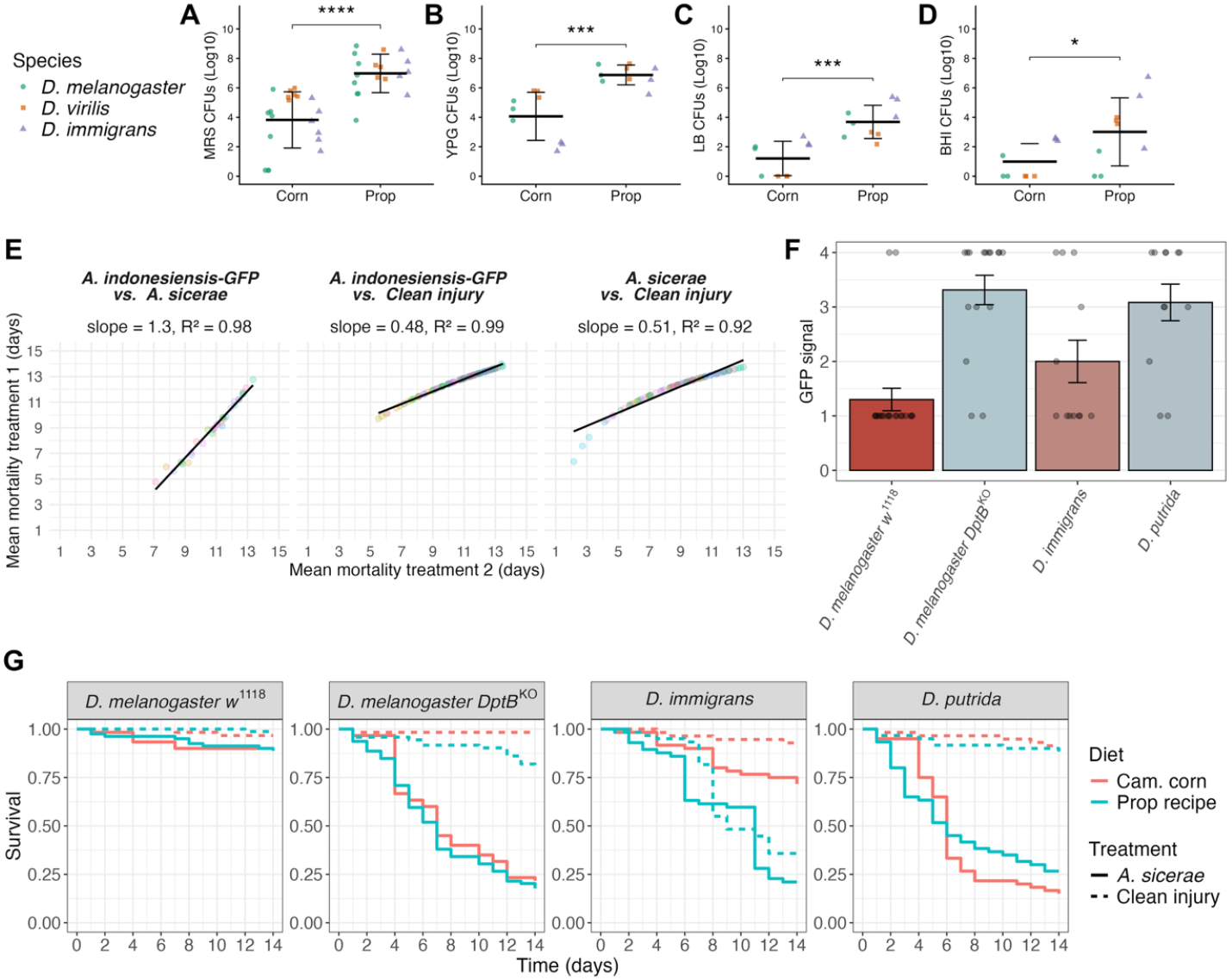
the Prop recipe exacerbates mortality compared to Cambridge cornmeal. **A-D)** Microbiota CFUs are orders of magnitude higher on the Prop recipe compared to Cambridge cornmeal on MRS agar (A), YPG agar (B), LB agar (C), and BHI agar (D). This pattern is consistent patterns across species. Overall significance: * = *P* < .05, ** = *P* < .01, *** = *P* < .001, **** = *P* < .0001. **E)** Survival correlations between infection or clean injury treatments. Data points coloured by species. Single data points for *A. sicerae* and *A. indonesiensis-GFP* are shown as these experiments were conducted at separate times. Each set of experiments was performed with its own matched clean injury controls. **F)** *A. indonesiensis-*GFP signal in flies 5 days post-infection across various species. GFP signal 1 = no signal, 2 = local fluorescence, 3 = majority of thorax fluorescent, 4 = systemic fluorescence. Each data point represents one fly. **G)** Infection of adults flipped on Cambridge cornmeal rescues survival after both clean injury and infection by *A. sicerae* in *D. immigrans*, but not *DptB*^*KO*^ or *D. putrida* flies, agreeing with expectations from our previous studies. Data in **(F)** and **(G)** reflect three replicate experiments with 20 males per vial. Barplots show Mean+StDev. Additional across-species data are presented in **Figure 4supp1,2**. *Acetobacter* infections used a dose of OD600 = 100 with flies kept at 22°C.

### The Prop recipe alters infection outcomes in species-specific ways

Infection experiments form the basis of our research program. In previous studies [9,10,31], all adult flies entering experiments were flipped on Cambridge cornmeal, EPFL cornmeal, or instant media, regardless of their larval diet. After transitioning to the Prop recipe for all species, we conducted an infection experiment across 42 fly species using *Acetobacter sicerae* and *Acetobacter indonesiensis-GFP*, bacteria that are virulent against *Diptericin B* (*DptB)* deficient *D. melanogaster* and mushroom-feeding *Drosophila* of the Quinaria and Testacea groups [10]. In these experiments, we flipped all infected adults in vials containing the Prop recipe and monitored survival for 14 days. While there was a high phylogenetic signal for mortality after *Acetobacter* septic injury (**Figure 4supp1**, phylogenetic heritability of mortality upon both infection treatments ≥ 0.70), we also observed a striking parallel mortality in the clean injury treatment, with Pearson correlations between infection and injury treatments having coefficients >0.90 (*A. indonesiensis-GFP* R^2^ = 0.99; *A. sicerae* R^2^ = 0.92, **Figure 4E**). Moreover, species such as *D. immigrans* were succumbing both to clean injury and *Acetobacter* infection, despite displaying high overall survival in a previous study [10]. By tracking *A. indonesiensis-GFP* signal, we confirmed that *D. immigrans* experiences far less *Acetobacter* proliferation compared to *D. melanogaster* flies lacking *DptB (DptB*^*KO*^), or *D. putrida* flies that lack *DptB* naturally (**Figure 4F**), suggesting the *D. immigrans* mortality we were observing was not due to the introduced *Acetobacter* infection itself (also see other species in **Figure 4supp2**).

Given increased microbiota loads on the Prop recipe (**Figure 4A-D**), we suspected microbiota coinfections were exacerbating the effect of injury and *Acetobacter* bacterial infection independent of the introduced *Acetobacter* itself. We next reared larvae on the Prop recipe, but flipped the emerging adults either on Cambridge cornmeal or the Prop recipe thrice weekly. Strikingly, we could rescue clean injury and *Acetobacter* mortality trends in *D. immigrans* (and other species) by flipping adults on Cambridge cornmeal (**Figure 4G, Figure 4supp2**). However, Cambridge cornmeal could not rescue the susceptibility of *Acetobacter*-sensitive *DptB*^*KO*^ or *D. putrida* flies. Thus, the Prop recipe contributes to increased mortality after injury and *Acetobacter* infection independent of the introduced pathogen itself.

### The microbiota of the Prop recipe includes opportunistic pathogens

We have recovered diverse putatively pathogenic taxa in our lab in Penryn, UK (e.g., *Acetobacter, Providencia, Serratia, Pseudomonas*). Curious if the correlation of clean injury and *Acetobacter* infections relied on pathogenicity contributed by our local microbiota strain of *Acetobacter*, we conducted a pilot experiment infecting a *Diptericin* mutant panel and select species with this *Acetobacter sp*. (*Acetobacter sp*. PEN). However, while this strain could kill immune-deficient *Rel*^*E20*^ flies within days, and caused some bloating in a subset of *DptB* mutants and *D. putrida*, it was not especially virulent within 21 days even against *Diptericin* mutants that are highly susceptible to other *Acetobacter sp*. (**Figure 5supp1**). Thus, we suspect that the sum effect seen in clean injury and *Acetobacter* infections we observed (**Figure 4supp1,2**) likely reflects the cumulative effect of injury, *Acetobacter*, and other coinfecting microbes with species-specific infection dynamics; as a result these mortality data produce a composite phylogenetic heritability signal unrelated to *Acetobacter* infection alone.

A previous study found a high phylogenetic signal of susceptibility to *Providencia rettgeri str Dmel* infection [31]. We were therefore curious if our lab *Providencia sp*. could be an opportunistic co-infecting pathogen contributing a confounding phylogenetic signal in our infection experiments where adults were flipped on the Prop recipe. Our local *Providencia* strain (JH10) appears to be a strain of *Providencia rettgeri* (**Figure 5A**). *Providencia rettgeri str Dmel* is a natural pathogen of *Drosophila* uniquely defended against by the *Diptericin A (DptA)* gene [10,32,33]. Interestingly, *P. rettgeri str JH10* is more virulent than *P. rettgeri str Dmel*, killing ~30% more wild-type *DrosDel w*^*1118*^ flies under identical conditions (**Figure 5B-D**). While the mortality of wild-type is high, and so the window of resolution is low, *P. rettgeri str JH10* produces a slight *DptA*-dependent increased mortality at 3 days post-infection compared to wild-type (**Figure 5D**, cox proportional hazards: *P* = 0.01 to 0.10 for all genotypes affected in *DptA*).

**Figure 5:**
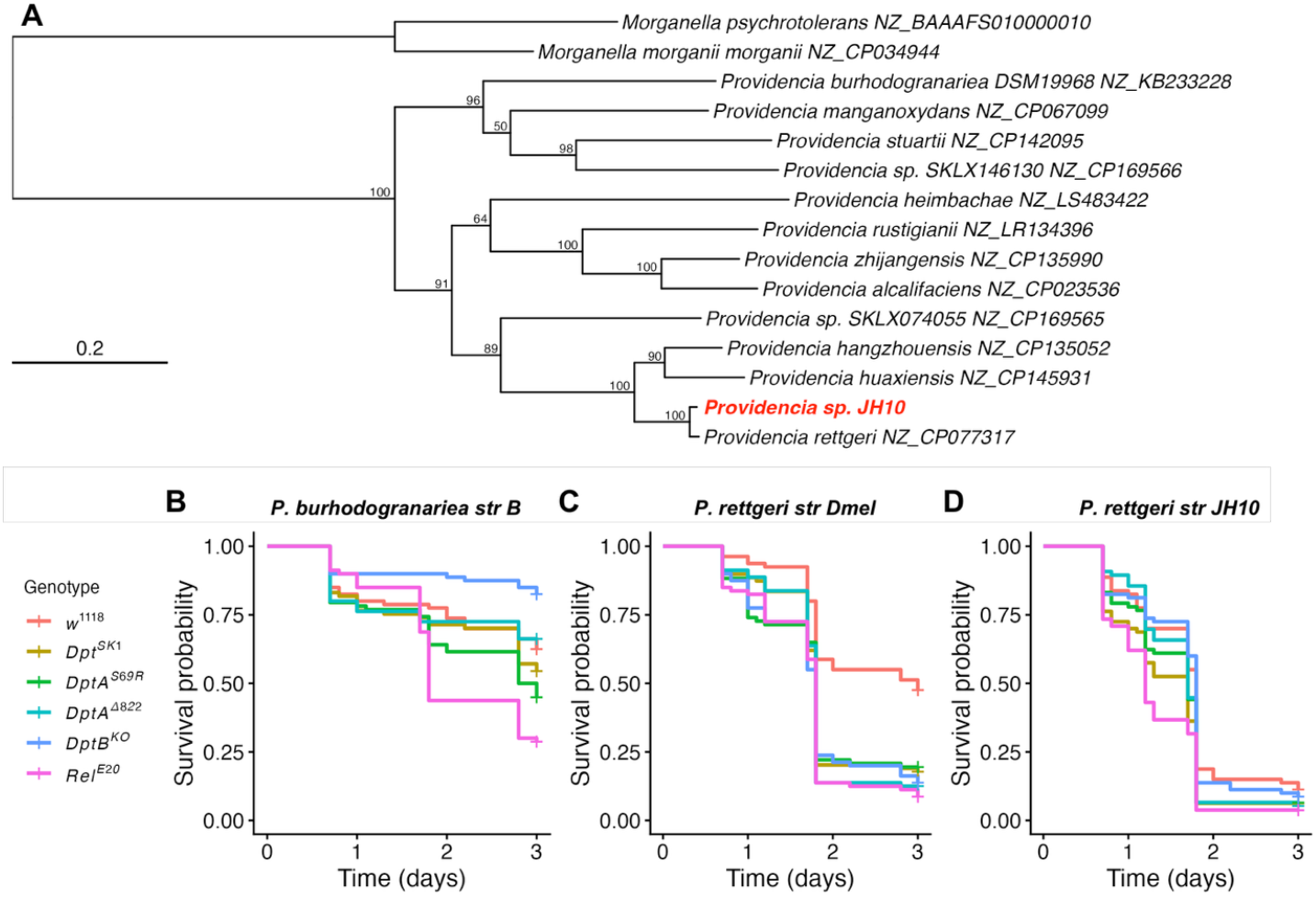
the microbiota of the Prop recipe may underlie exacerbated mortality of flies flipped on this diet. **A)** Maximum likelihood phylogenetic tree (100 bootstraps) suggests the *Providencia* species present in our local microbiota is a strain of *P. rettgeri*. **B-D)** Mortality of flies infected by **B)** *P. burhodogranariea str B*, **C)** *P. rettgeri str Dmel*, or **D)** *P. rettgeri str JH10*. Infections performed with OD600 = 0.2 bacterial pellets at 22°C, with 20 adult males flipped on the Prop recipe. Data reflect four replicate experiments.

Collectively, these results highlight important caveats to the use of the Prop recipe. While this recipe rears all fly species, its rich microbiota may impact the outcome of certain experiments in a species-dependent manner. The microbiota encouraged by different food media can include virulent opportunistic pathogens, such as *P. rettgeri str JH10* or other microbes (e.g., *Serratia, Pseudomonas*), that could contribute to coinfection dynamics and knock-on effects to the phenotype of interest. However, while we provide this cautionary note, not all phenotypes are likely to be impacted as dramatically as what we have seen in our, rather traumatic, injury and infection experiments. Moreover, we could greatly rescue confounding effects by simply flipping adults on a food media that does not support microbiota growth to the same extent. Finally, while we report these caveats that affected various species in our hands, diet did not impact the clean injury survival of *D. melanogaster* wild-type flies (**Figure 4G**), but did show trends for *DptB* mutants, suggesting an importance of considering how diet and microbiota effects affect mutant studies.

## Discussion

Here we describe the versatility of the Prop recipe in rearing fly species from across the genus *Drosophila*. By providing various fitness parameters in our hands, we hope our data give confidence in this recipe, and our rearing protocols enable other groups to adopt the use of diverse *Drosophila* species for evolutionary research. Indeed, the variety of sequenced species and their ease-of-use make *Drosophila* an excellent model system, wherein a major challenge to their use was simply rearing diverse species with unique dietary needs.

We provide data for 66 species reared on the Prop recipe in our hands. We have further successfully reared other species, such as *Scaptomyza pallida*, for many months. In other cases, bringing certain strains of species into the lab resulted in poor viability, ultimately failing to establish a stable culture. However, on a second attempt, we found success for such species, which have been stably maintained since; specific examples include *Drosophila tenebrosa, Drosophila subquinaria*, and *Drosophila neotestacea*. Thus, some initial challenges to importing species into the lab may be due to stresses outside dietary control, such as the stress of being shipped, or cryptic viruses or microbial commensals that are benign in healthy individuals, but become pathobionts in stressed flies, complicating their establishment after shipping. Alternately, some strains within a species could perform better or worse on the Prop recipe due to genetic polymorphisms in virus restriction factors [34], or antimicrobial peptide genes [35,36], that could impact the virome and microbiota and so help to suppress opportunistic infections. Attempting to rear different strains for some species could show they are not intrinsically unable to survive in lab conditions, but could artificially select for strains with immune alleles competent against lab-specific microbiota.

We observed a greater mortality rate of *Acetobacter*-susceptible *Diptericin B* mutant flies when flipped on the Prop recipe after clean injury compared to Cambridge cornmeal (**Figure 4G**). We also observed over 1000 times greater *Acetobacter* loads in Prop recipe vials compared to Cambridge cornmeal (**Figure 4A**), and if introduced as a systemic infection, our local *Acetobacter* strain can cause abdominal bloating in *DptB* mutants also seen in our previous study using other *Acetobacter* strains [10]; bloating is a disease phenotype likely related to malpighian tubule damage, the fly analogue of the kidneys [37]. *Acetobacter* stemming from the lab microbiota could therefore have exacerbated clean injury mortalities. However, we also observed *Acetobacter-*independent mortality of species like *D. immigrans* associated with flipping *D. immigrans* on the Prop recipe. Indeed, we found far higher non-*Acetobacter* CFUs in vial microbiota assessments in *D. immigrans* compared to *D. melanogaster* (**Figure 4C,D**). Thus, we suspect non-*Acetobacter* members of the microbiota opportunistically coinfect when present in the vial microbiota, contributing to increased mortality caused by flipping on the Prop recipe. Microbes such as our local strain of *P. rettgeri str JH10*, or a number of other potential pathogens (e.g., *Serratia, Pseudomonas*) could be coinfecting these flies. As a result, the signals seen in our *Acetobacter* infections across the phylogeny are likely multi-faceted and not due to *Acetobacter* virulence alone. Our variable survival results when rearing flies on the Prop recipe and different cornmeal recipes (**Figure 3C, File S1**), alongside differences in the microbiota depending on the food, suggests future studies will benefit from monitoring diet-dependent microbiota differences (also see [19,38–42]).

In infection experiments conducted here, we only monitored survival over relatively short timeframes. Longer-term experiments (such as lifespan studies) may reveal compounding effects of a richer microbiota carried over from vial to vial [26,43,44], and so the Prop recipe may negatively impact lifespan relative to microbiota-poor media such as Cambridge cornmeal. Ultimately, we now rear flies on the Prop recipe, but flip adults for infection experiments on the Cambridge cornmeal diet to reduce the potential for confounding secondary infections. We provide detailed infection data here to emphasise possible secondary effects of changing the diet, which could affect some experiment types more than others. This diet-specific trait may even be exploitable in future studies interested in the effect of an enriched microbiota.

With the advent of CRISPR/Cas gene editing, research groups are increasingly pursuing transgenic tools in other fly species [45]. The Prop recipe originally derives from the Cambridge Fly Facility “Propionic” media, and this facility offers commercial transgenic injection services. Thus, this supplier is already capable of rearing diverse fly species for commercial injections. Naturally, other suppliers could adopt the Prop recipe for rearing alternate species.

The discovery that the Prop recipe can successfully rear such ecologically-diverse species is sheer serendipity. This exploratory and descriptive effort was inspired by a new group member realising some difficult to rear species in their prior experience were already being reared on the Prop recipe in the Penryn facility with ease. Ultimately, most all species we had been rearing on the Cambridge cornmeal recipe showed improved fitness on the Prop recipe. While the *D. melanogaster* communities studying immunity, the microbiome, behaviour, and adjacent fields know the importance of diet and its effects on the microbiota, the variation we observed in host physiology and fitness across different food types suggests there is likely a high variability in basal health even for *D. melanogaster* stocks reared in different research groups (as emphasised by [25]). We would welcome feedback and reporting of successes or failures of research groups attempting to adopt the Prop recipe and our protocols.

There have been hundreds of drosophilid genomes sequenced recently [46], offering a very high resolution for molecular evolutionary studies. As a result, collecting phenotypic data across the full diversity of *Drosophila* species can now be followed-up with genomic investigations into species-level differences. Numerous studies are further generating transcriptomic data across diverse fly species, offering high levels of precision in gene annotation and expression insights [47]; indeed we are generating transcriptomic data from dozens of species reared in our lab, which we will make fully available to the community when ready for public release. There is thus a building momentum for *Drosophila* as a model of evolutionary genetics. Our hope is this simple study allows researchers to gather phenotypic data across *Drosophila* species to pair with developing genomic and transcriptomic resources, and so produce novel insights of evolutionary genetics.

## Author contributions

### Data collection

HS, LH, JJ, MC, EGR, AJ, HW, MAH

### Data analysis

HS, MC, EGR, MAH

### Writing-initial draft

MAH

### Writing-editing

HS, MC, BL, MAH

### Supervision

BL, MAH

## Acknowledgements

We thank Noah Whiteman and Susan Bernstein for providing *Scaptomyza pallida* and Helen White-Cooper for *D. miranda*. This work was funded by Wellcome Trust fellowship 227559/Z/23/Z and Swiss National Science Foundation fellowship P500PB_211082 awarded to MAH, and Henry Dale fellowship 109356/Z/15/Z jointly funded by the Wellcome Trust and UK Royal Society awarded to BL. We also thank the China Scholarship Council for PhD studentship funding to HS supervised by BL.

## Materials and Methods

Data and scripts are included in **File S2**.

### Fly strains used in this study

Species used in this study are provided in Table S1. For Figure 5 infections, isogenic DrosDel strains were used per Hanson et al. [10], including *DptB*^*KO*^ flies that also express *DptA* at lower levels than their *w*^*1118*^ control line. *Relish* mutants *(Rel*^*E20*^) used in this study were not isogenic, and were a gift from Jean-Luc Imler. We additionally used FlyBase to determine strain genetic allele effects [14].

### Experimental parameters

Flies used in this study were reared with a 12h:12h light-dark cycle at ~50% humidity at 22°C. Approximately 1-2 week old adult flies were reared with loosely controlled adult densities of ~30 flies per 28.5mm wide vial, flipped twice per week to ensure egg laying was controlled to a certain density and to prevent mixing of generations. Some less fecund species were instead flipped weekly, allowing greater densities to suppress biofilms or mould and improve vial success. Egg-to-adult times were defined as the time between flies being added to the vial and the day of first emergence. Anaesthetised flies were allowed to recover at least two days before recording vial start dates for egg-to-adult experiments.

Wet masses were collected by measuring the mass of empty screw cap O-ring-sealed microtubes prior to adding pools of 5-7 day old flies flipped on the same food recipe as their larval diet. A high-precision analytical balance accurate to 10µg (Sartorius 120g-0.01mg) kept on a shock-absorbing balance table was used for tube measurements. Microtubes containing flies were kept at −20°C until they could be measured, and still sealed, briefly dried in a drying oven at 50°C prior to measurement to prevent any condensation adhering to the tubes. Differences in emergents-per-female and body mass were analysed by glm with diet-by-species interactions and emmeans *P*-values reported for within-species diet-dependent differences in R 4.4.1.

We used a buoyancy-based screening strategy to identify larvae that have higher levels of stored fat [28]. We measured the proportion of larvae floating in 10% sucrose solution. Each replicate vial was the product of 10 adult male and 10 adult female flies aged for 2 days in individual fly vials then mixed together and allocated to vials containing an abundance of food from each diet. Parent flies were allowed to lay eggs for one week before adults were removed. Third instar larvae were collected from each diet over the following two weeks; emergence rates and development times vary between species and diet, requiring a dynamic data collection window. 15 Third instar larvae were put into cuvettes containing 1.5 mL of 10% sucrose dissolved in PBS (illustration in **Figure 3C**), and the number of floating larvae were recorded over three parallel blocks. Successful species and diet replicate blocks ranged from zero (no larvae, e.g., *D. testacea* reared on Cambridge cornmeal) to three (larvae present and screened in all three blocks, e.g., *D. melanogaster* reared on the Prop recipe).

In diet comparison experiments collecting data on egg-to-adult times, body mass, and total emergence, we reared flies in 25mm vials with 10 males and 10 females allowed to recover from anaesthesia and habitutate to their new diet for three days prior to egg laying. These flies were allowed to lay eggs over 24 hour periods before being flipped to the next vial. As such, differences in flies emerging from each diet could depend on adult oviposition and associated larval density, or larval viability interacting both with diet, and with larval density effects. However, our aim in these experiments was to assess the utility of each diet in rearing a diversity of species, and not to focus on disentangling behavioural versus density effects.

### Host phylogeny

The *Drosophila* phylogeny used in this study was produced using nucleotide and codon alignments of *16S* and *28S* ribosomal RNA (rRNA), mitochondrial *COI* and *COII*, and nuclear *adh, Gpdh1, RpL32, Sod1*, and *DptA*; in our phylogeny, we included *DptA* to act as a molecular synapomorphy to aid with phylogenetic sorting of the two main subgenera [48,49]. Gene alignments (**File S3**) were input into BEAST v1.10.4 with parameters matching those used previously [9]. In brief: we used distinct HKY nuclear substitition models for rRNA [1, 2, 3], mitochondrial [1+2, 3], and nuclear [1+2, 3] genes. We used a random starting tree with a relaxed uncorrelated lognormal clock and a speciation-extinction (birth-death) process. The BEAST analysis was run for 500 million MCMC generations sampled every 5000 steps.

For phylogenetic heritability analyses, we used an MCMCglmm approach with species-level variation included as a random effect giving the formula for heritability as phylogenetic *h*^*2*^ *= Variance*^*phylogeny*^ */ Variance*^*phylogeny*^ *+ Variance*^*species*^, as used previously [9,31].

Phylogenetic trees shown in **Figure 2** used the phytools package contMap() function with default settings: ancestral states were computed via maximum likelihood under a Brownian motion model of continuous trait evolution.

### Bacterial phylogeny

We sequenced the genome of *P. rettgeri str JH10* (GenBank BioProject PRJNA1364308) with MicrobesNG^©^ using a combination of Illumina 250bp paired end reads and Oxford Nanopore long read sequencing. The genome of *P. rettgeri str JH10* was *de novo* assembled using SPAdes with default parameters in Geneious Prime 2025.2.2. Genomes of *Providencia* species and *Morganella* species outgroups were downloaded from NCBI per identifiers reported in **Figure 4A**. Best BLAST Hits (BBHs) were extracted for the genes *ychF, rplB, rplE, lepA*, and *16s rRNA* as core phylogenetically robust genes for bacterial phylogenetics (per [50]). MAFFT alignments were performed in Geneious and concatenated prior to PhyML phylogeny construction implemented in Geneious with default parameters (100 bootstraps).

### Bacterial CFU counts

Microbiota loads were collected from 25mm vials where 5 males from each species had been flipped every three days on either Cambridge cornmeal or the Prop recipe for a total of nine days. Vial food surfaces were washed with 1000µL sterile water, and this solution was then serially diluted and plated on various bacterial growth media agar plates: De Man–Rogosa– Sharpe (MRS), Luria-Bertani (LB), Yeast-Peptone-Glucose (YPG), and Brain Heart Infusion (BHI). MRS and YPG agar are permissive to growth of *Acetobacter* and *Lactobacillus* within 24-48h at 29°C, bacteria that differ in growth rate and colony morphology. LB and BHI agar preferentially promote the growth of *Providencia, Morganella, Serratia, Enterobacter*, and more. Plates were grown overnight at 29°C before manual colony counting. We additionally left plates for another 72h checking daily for additional slower-growing colonies, which did not appear in numbers sufficient to at all affect the conclusions of the 24h time point.

Vial microbiota load differences were tested with one-way ANOVA with Tukey’s HSD (CFUs ~ Diet+Species). MRS and YPG agar displayed an overall difference in identifiable *Acetobacter* colonies (the majority of colonies observed) between the Prop recipe and Cambridge cornmeal (P < .001), and total bacterial CFUs were also significantly higher for LB and BHI on Prop recipe compared to Cambridge cornmeal (P < .001 and P < .05 respectively). MRS agar also displayed various smaller colonies at a density ~20x lower than *Acetobacter* in *D. immigrans*, and *D. virilis* that were consistent with *Commensalibacter sp*. detected in our microbiota previously [30]. In addition, *D. immigrans* displayed colonies on BHI uniquely when flipped on the prop recipe, and *D. melanogaster* displayed almost no colonies on BHI in general, suggesting species specific and species-by-diet-specific microbial taxa.

### Infections

Infection experiments were performed using a 0.1mm insect pin dipped into the concentrated bacterial pellets prior to piercing the thoracic pleura. All experiments were conducted at 22°C with up to 20 flies per 25mm vial. OD600 doses were *Acetobacter* OD600 = 100 ± 10% and *Providencia sp*. OD600 = 0.2 ± 10%. Also see **File S1** for an additional across-species egg-to-adult and infection experimental dataset comparing the outcome of infection with *P. rettgeri str Dmel, S. aureus*, and *Drosophila C virus* for flies reared on either the Cambridge cornmeal or Penryn cornmeal diets, prior to flipping on the Cambridge cornmeal as adults. These experiments suggest a species-specific effect of larval diet on infection outcome, which we note here to provide initial data on diet-dependent and species-specific susceptibilities to these pathogens.

### Generative AI usage statement

We used ChatGPT to assist with coding for data preparations and figure generation, with supervision and highly specific prompts. For instance: *“I’d like to plot df_barplot as a barplot with facet wrap by SpeciesName, fill coloured by Diet, and data point type by ExperimentBlock*.*”* Generative AI was not used at any point to assist with writing.

## Supplementary figures

**Figure 2supp1:**
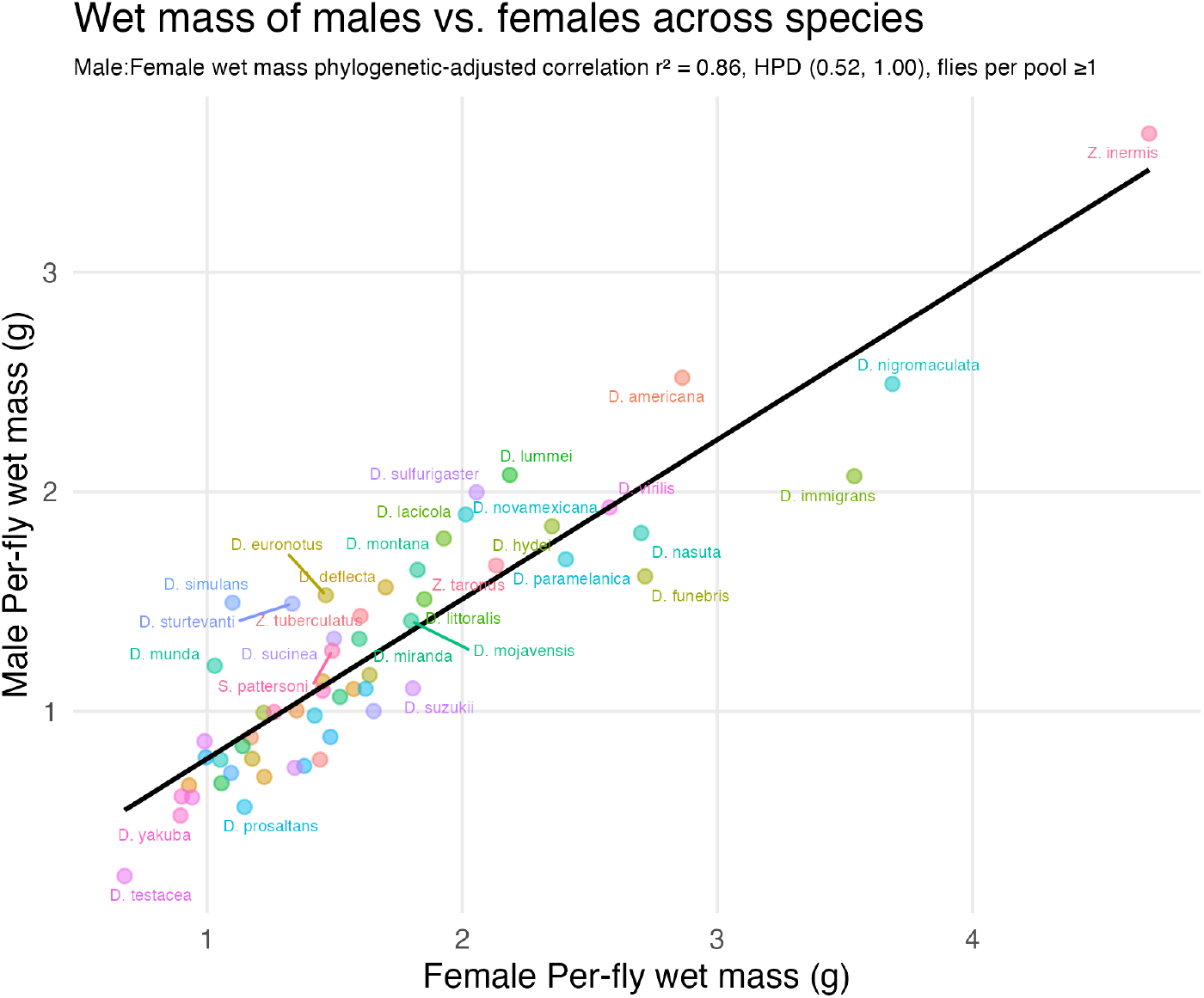
Correlation of male-female wet mass differences across species. Test statistic calculated from a bivariate phylogenetic MCMCglmm model wherein the correlation coefficient was calculated from the variance of the phylogenetic signal divided by the square root of the product of female phylogenetic and species variance multiplied by male phylogenetic and species variance; i.e. for an MCMCglmmm output named df with species given as “spp”, *r*^*2*^ = (df$VCV[,”sexF:sexM..animal”]/((df$ $VCV[, “sexF:sexF.animal”]+ df$VCV[, “sexF:sexF.spp”]* (df$VCV[, “sexM:sexM.animal”]+ df$ VCV[, “sexM:sexM.spp”])))^2^.

**Figure 3supp1:**
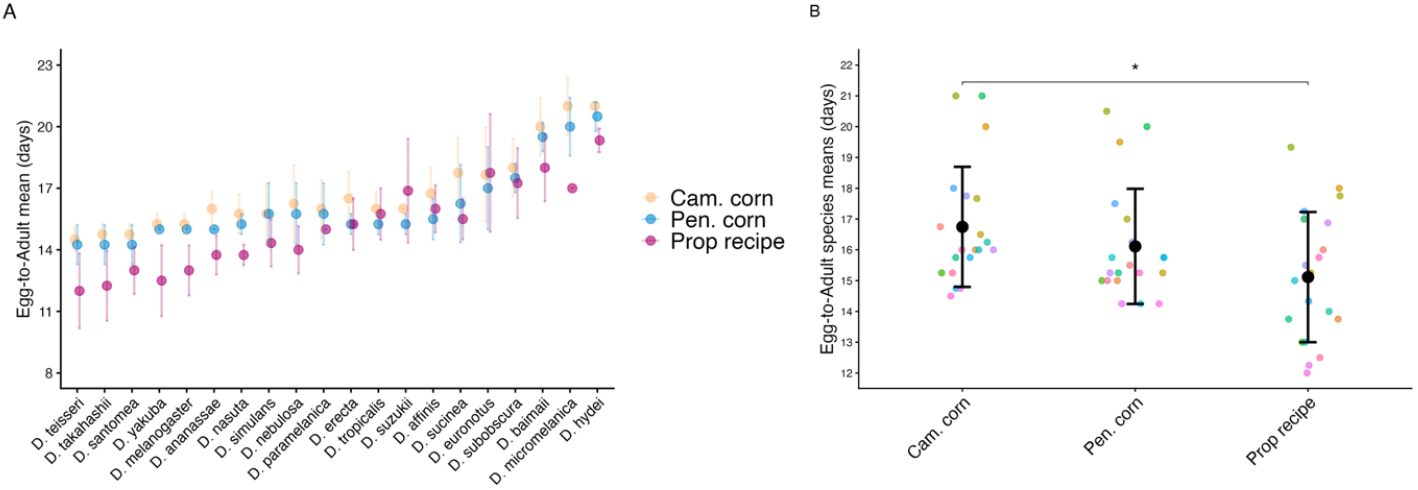
egg-to-adult times for select species on three food types. **A** and **B** show the same data with different presentations. The Cambridge cornmeal recipe tended to produce the longest egg-to-adult times, while the Prop recipe produced the shortest (Cambridge cornmeal vs. Prop, Tukey’s HSD: *P* = 0.03). Penryn cornmeal was intermediate, with minimal difference between Penryn cornmeal and Prop (Tukey’s HSD: *P* = 0.26). In **(B)**, data points are coloured by species. Error bars represent mean ± standard deviation.

**Figure 4supp1:**
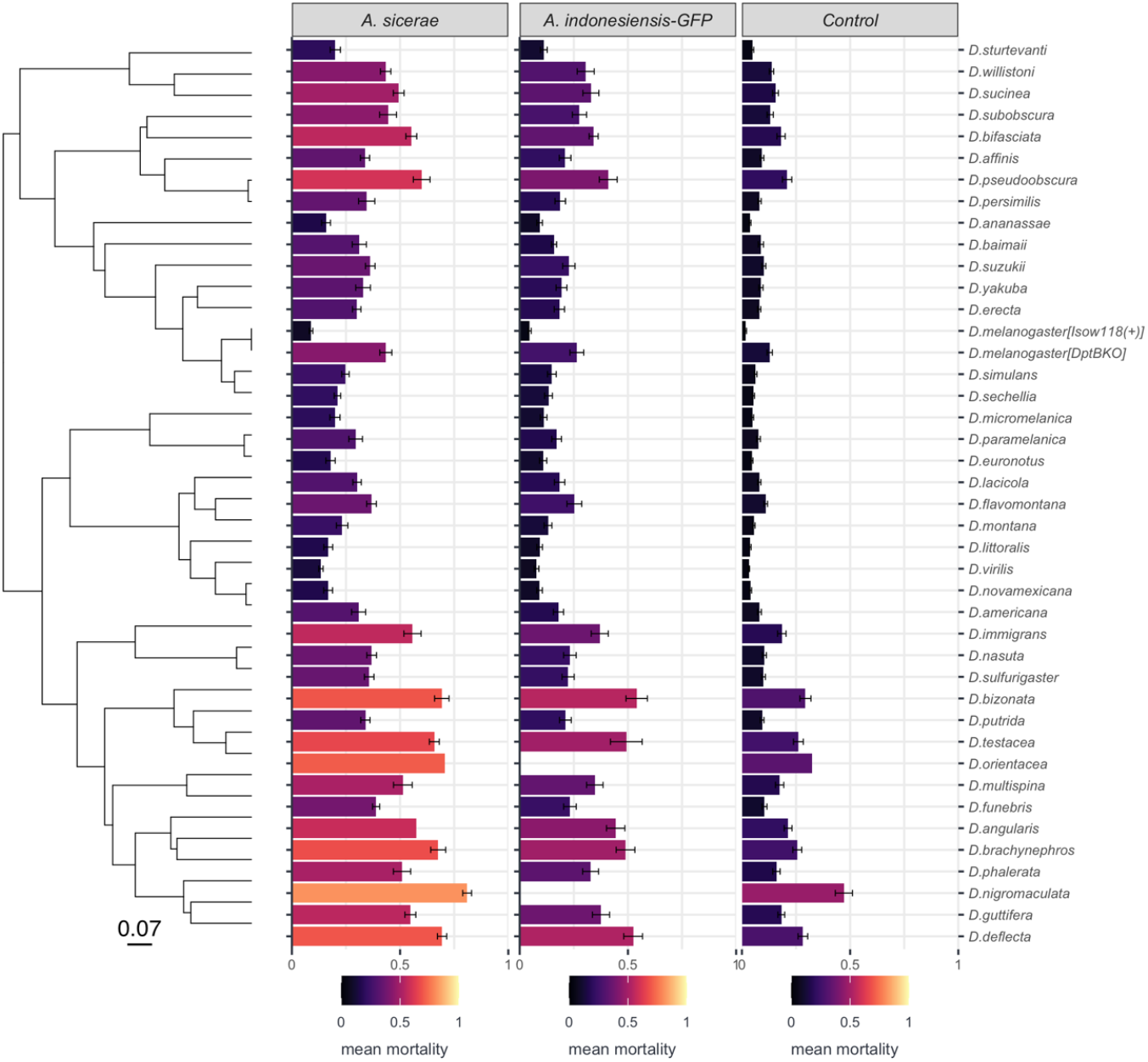
mortality after *Acetobacter* infection or clean injury of flies kept on the Prop recipe. Because the bacterial infection series were performed at different times, data shown here reflect up to three replicate experiments with 20 males per vial for *Acetobacter* infections, while clean injury reports the sum of all six replicate experiments. Barplots show Mean+StDev. *Acetobacter* infections used a dose of OD600 = 100 with flies kept at 22°C. The phylogenetic heritability *(σ*^*2*^_*p*_*/(σ*^*2*^_*p*_*+ σ*^*2*^_*s*_) of each treatment is as follows (mean, HPD upper-lower): *A. sicerae* (0.71 ± 0.24, 0.00-0.95), *A. indonesiensis GFP* (0.70, 0.00-0.95), Clean injury control (0.68, 0.00-0.95). Species-by-treatment interactions with no data (unavailable during that experiment) have no bar entered (e.g. *A. indonesiensis GFP* * *D. nigromaculata*).

**Figure 4supp2:**
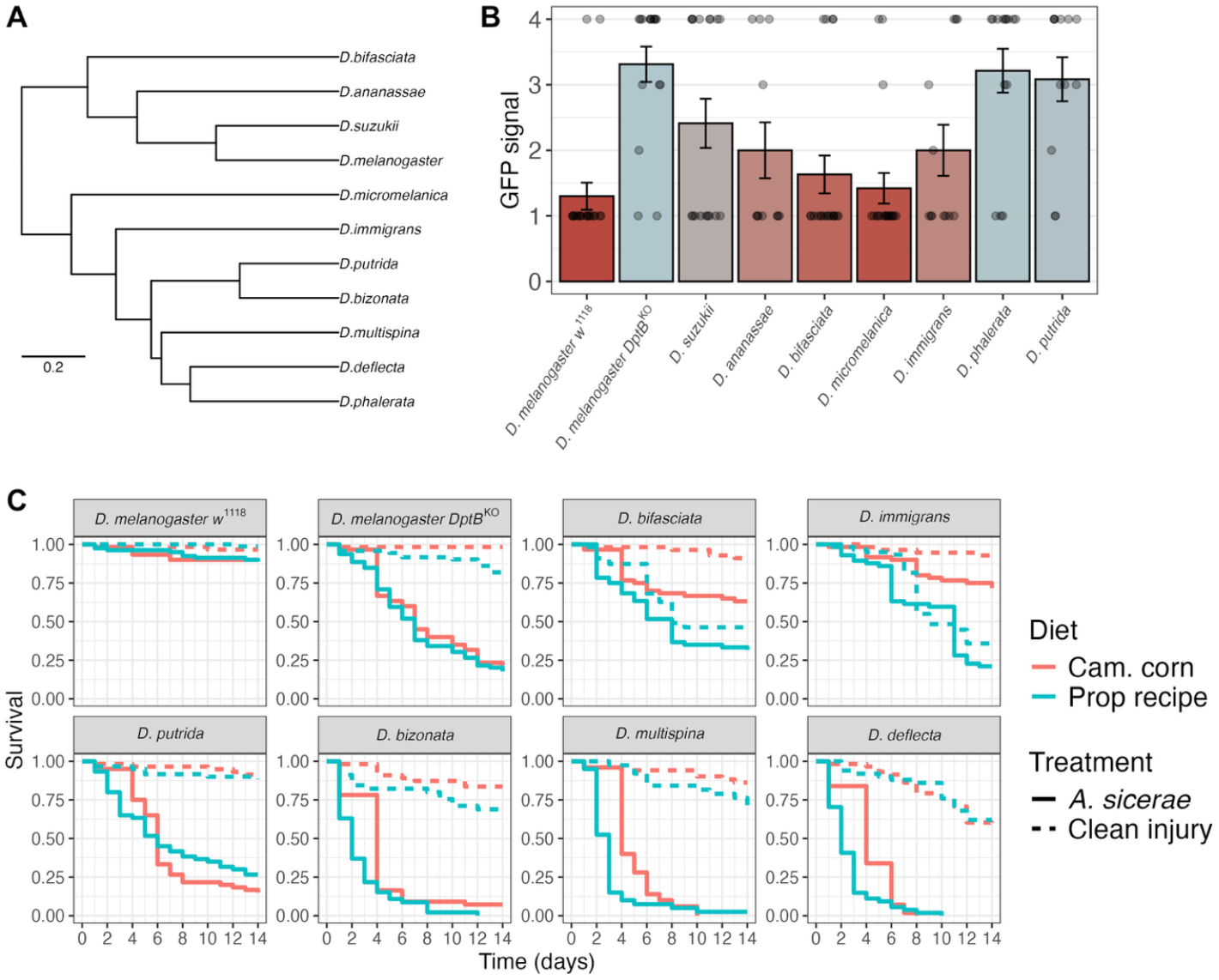
complete species data for experiments shown in Figure 4F and 4G. **A)** Phylogeny of species used. **B)** GFP signal observed 5dpi as in Figure 4. **C)** Survival data of species reared on the Prop recipe but flipped on either Cambridge cornmeal or the Prop recipe as adults, given clean injury or *A. sicerae* infection. Like *D. immigrans, D. bifasciata* mortality is comparable between *A. sicerae* infected-flies and clean injury when flipped on the Prop recipe but not the Cambridge cornmeal diet. Moreover, many species trend towards higher clean injury mortality when flipped on the prop recipe (*D. melanogaster DptB*^*KO*^, *D. bifasciata, D. immigrans, D. bizonata, D. multispina*), while the opposite trend is never observed. Likewise, for *A*. sicerae-infected flies, mortality tended to be higher if adults were flipped on the Prop recipe (*D. bifasciata, D. immigrans, D. bizonata, D. multispina, D. deflecta*), and the opposite trend was not observed in any species; *D. putrida* curves cross over, and so overall mean mortality between the *A. sicerae* and clean injury control treatments is comparable.

**Figure 5supp1:**
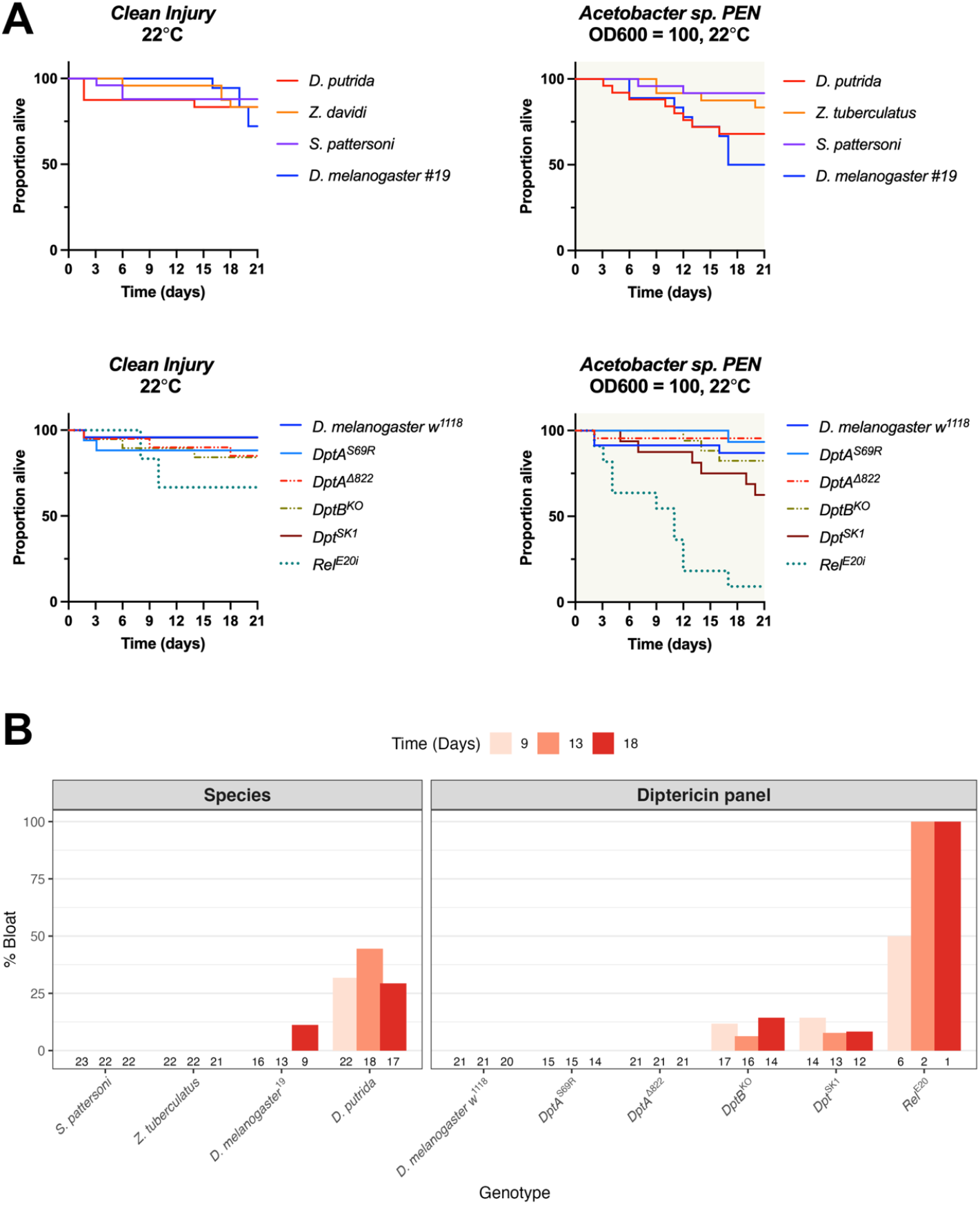
the *Acetobacter sp*. in the Penryn *(Acetobacter sp. PEN)* local microbiota is not an especially virulent pathogen. **A)** While this microbe does kill *Relish* mutants lacking the Imd pathway (*Rel*^*E20*^), it is not capable of rapidly killing even *DptA, DptB* double mutants (*Dpt*^*SK1*^) or *Acetobacter-* sensitive *D. putrida*. **B)** *Acetobacter sp. PEN* does cause bloating over time in *D. putrida* and *D. melanogaster* mutants lacking *DptB*. Numbers along the bottom of the plot represent surviving individuals at each time point. Bloating was scored per Hanson et al. [10]. These data come from a single experiment flipped in Prop recipe 28.5mm vials (n = up to 25 males per vial). *D. melanogaster*^*19*^ and *D. melanogaster w*^*1118*^ are independent wild-type strains used previously [10,31]

