## Supplementary material for "Life history and infection susceptibility parameters of *Drosophila* species reared on a common diet": File S1

**Supplementary File 1**

**Crude emergence data using Cambridge cornmeal and PENSTRAS recipes**

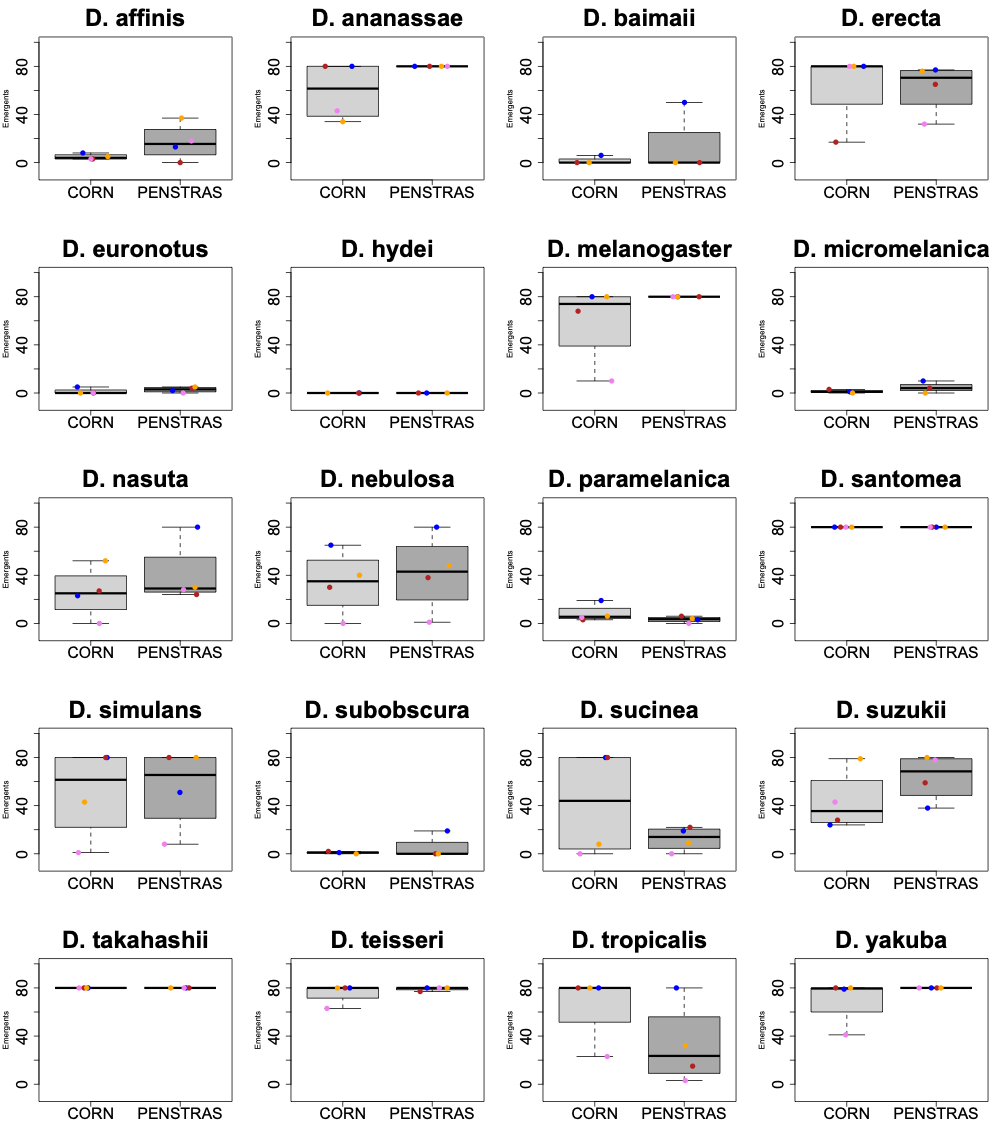
 **Supplementary File 1 Figure 1:** total male emergents emerging within 3 weeks from rearing species in bottles of either Cambridge cornmeal (CORN) or PENSTRAS food. A maximum of 80 males were recorded. Data point colours represent unique experiment blocks. In the majority of cases, rearing on PENSTRAS improved total emergents, though in a couple species (*D. sucinea, D. tropicalis)* the opposite is true. Total adults in each bottle were controlled to ~60 total flies flipped weekly. Note: low emergent totals may reflect poor rearing success, egg-to-adult times beyond our observation period, or both.

**Infection experiments using Cambridge cornmeal and PENSTRAS recipes**

Prior to our use of the Prop recipe to rear all species, we experimented with how rearing different fly species on either the Cambridge cornmeal recipe or the PENSTRAS cornmeal recipe affected survival to infection. In these experiments, we reared flies in bottles containing the larval diet food type, then flipped emerging adults on Cambridge cornmeal food. We provide these data here as a demonstration of the impact that larval diet can have on host fitness. Across experiments, we found that PENSTRAS and the Prop recipe provided similar egg-to-adult development times for species amenable to PENSTRAS rearing, however at the time we conducted these experiments we were not yet rearing all species on the Prop recipe, and so we cannot provide survival data for these species*pathogen interactions when reared on the Prop recipe in this manuscript.

**Methods**

Species were reared on either the Cambridge cornmeal food recipe or the PENSTRAS food recipe. Males were sorted and up to 20 males per vial were included in subsequent survival experiments. Collected adult males were flipped on Cambridge cornmeal prior to and following infection. Because some species-by-food interactions resulted in poor emergence rates, some species-by-pathogen survival data contain only a few individuals.

Infections were performed with three different pathogens: *Drosophila C Virus (DCV)* (TCID_50_ ≈ 6.3 x 10^9^), *Providencia rettgeri* (OD600 = 1), and *Staphylococcus aureus* (OD600 = 1). Three experimenters were involved in the septic injury experiments, with experimenter-by-pathogen changed to ensure unique combinations between each experiment block, allowing a universal comparison of experimenter-mediated variation across pathogen types.

To test for the overall effect of larval diet on survival to infection, cox mixed effects models were performed with larval diet as a fixed factor and experimenter, experiment block, and fly species as random factors. Our sampling was insufficiently powered to statistically model food-by-species interactions, however we provide total flies as “n =” per larval diet within figure panels to inform the reader of the approximate confidence that each species-by-food-by-pathogen interaction provides.

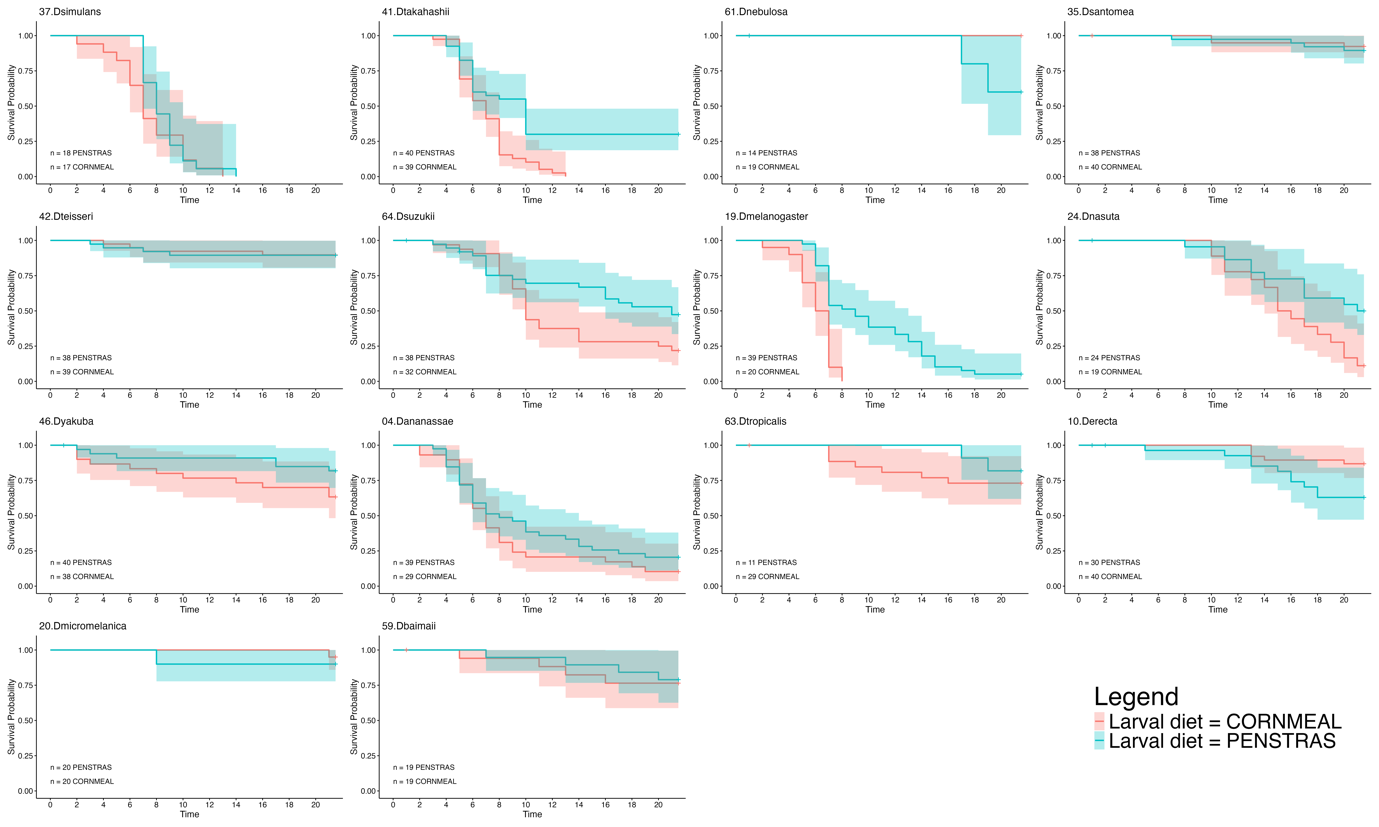

**DCV survival**

**Overall effects:**

Mixed effects coxme model

Formula: Surv(Time2, Censored) ~ LarvalDiet + (1 | Person) + (1 | Experiment) + (1 | Genotype)

Data: FlyData

events, n = 356, 808

Random effects:

group variable	sd	variance

1	Person Intercept 0.0200000 0.00040000

2 Experiment Intercept 0.2285474 0.05223392

3	Genotype Intercept 1.3748888 1.89031934

Chisq	df p	AIC	BIC

Integrated loglik 426.4 4.00 0 418.4 402.9

Penalized loglik 486.0 14.23 0 457.5 402.4

Fixed effects:

coef exp(coef) se(coef)	z	p

LarvalDietPENSTRAS -0.5512	0.5763	0.1127 -4.89 1.02e-06

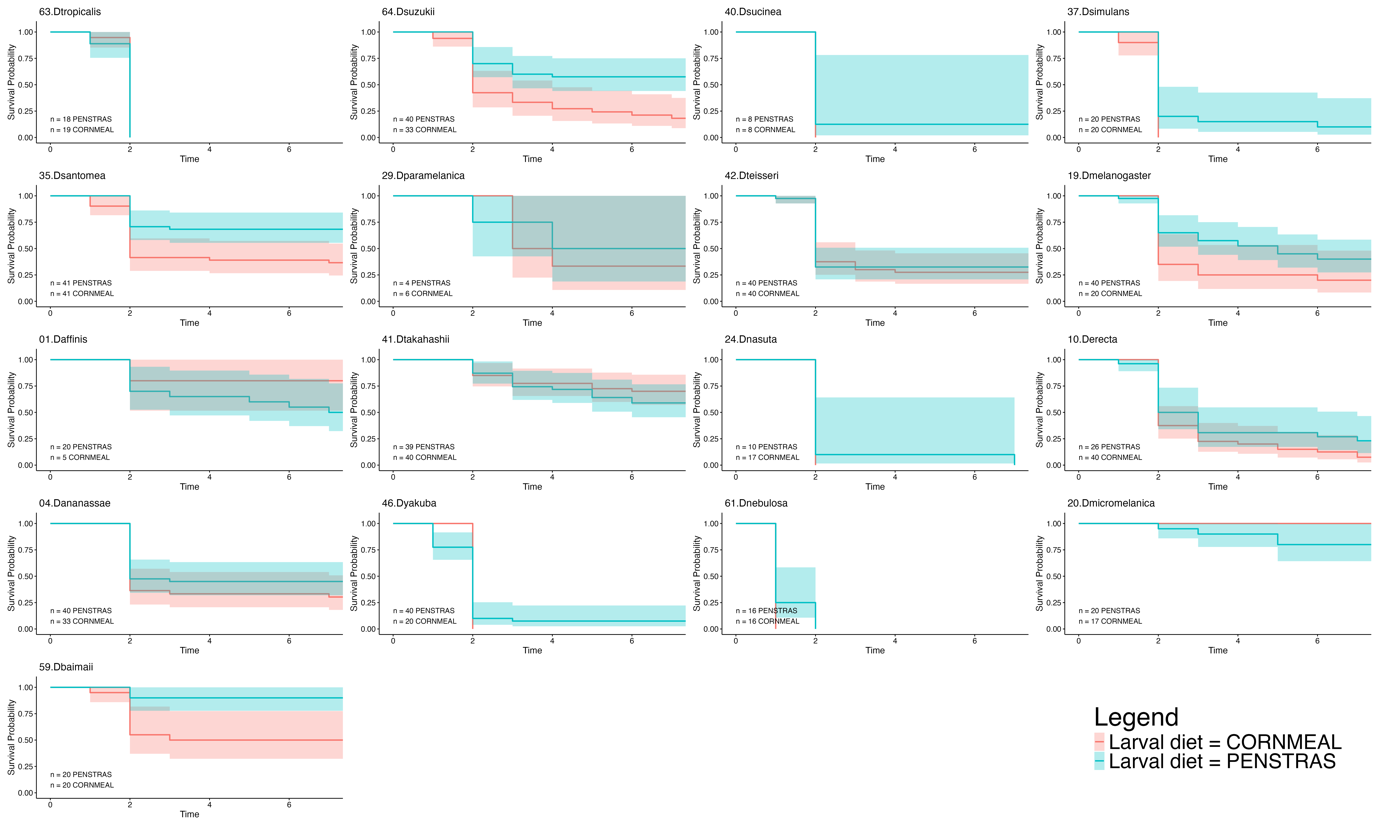

**P. rettgeri survival**

Mixed effects coxme model

Formula: Surv(Time2, Censored) ~ LarvalDiet + (1 | Person) + (1 | Experiment) + (1 | Genotype)

Data: FlyData

events, n = 548, 837

Random effects:

group variable	sd variance

1	Person Intercept 0.8332925 0.6943765

2 Experiment Intercept 0.4040307 0.1632408

3	Genotype Intercept 0.9287837 0.8626391

Chisq	df p	AIC	BIC

Integrated loglik 452.8 4.00 0 444.8 427.6

Penalized loglik 524.4 17.11 0 490.2 416.6

Fixed effects:

coef exp(coef) se(coef)	z	p

LarvalDietPENSTRAS -0.33374	0.71624 0.08805 -3.79 0.00015

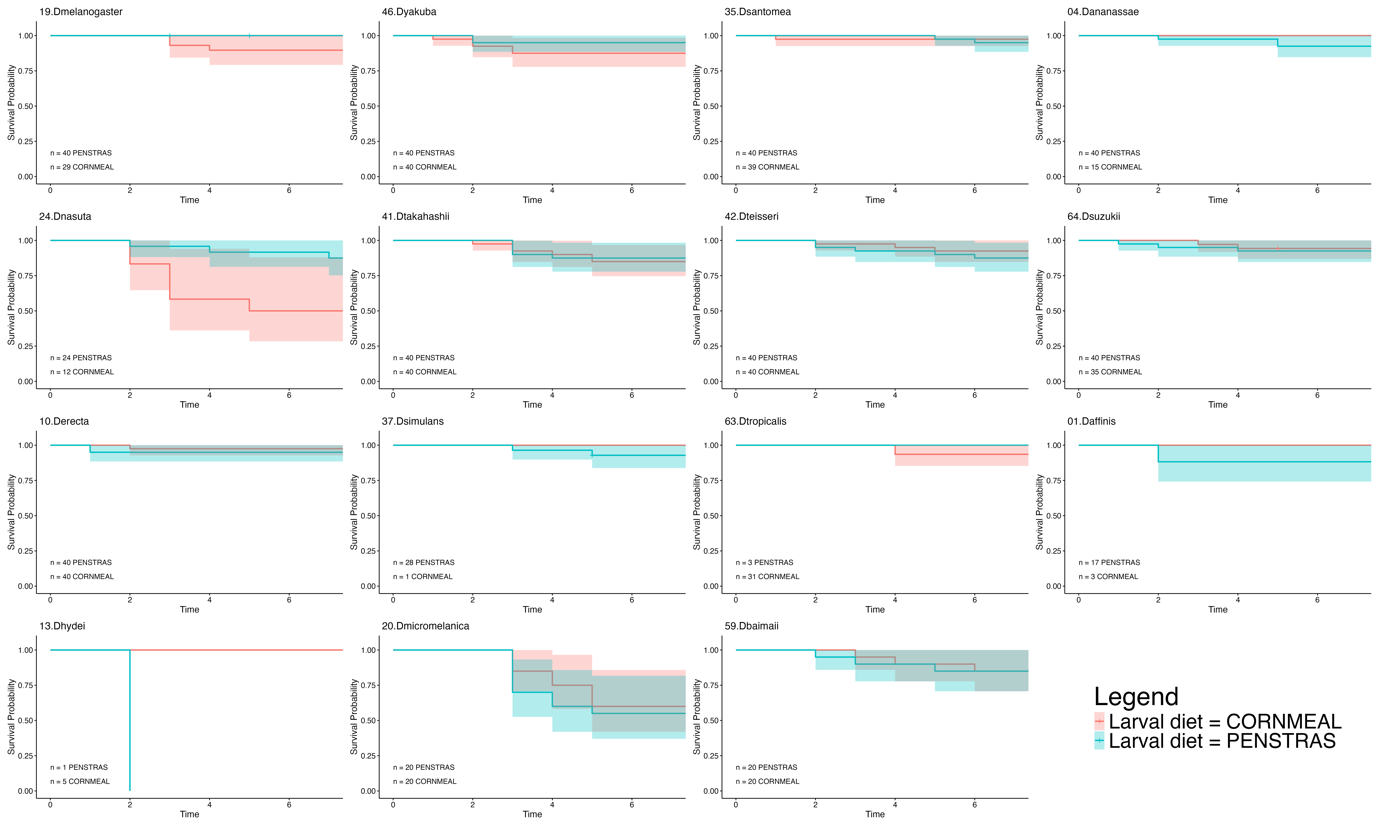

**S. aureus survival**

Mixed effects coxme model

Formula: Surv(Time2, Censored) ~ LarvalDiet + (1 | Person) + (1 | Experiment) + (1 | Genotype)

Data: FlyData

events, n = 82, 803

Random effects:

group variable	sd	variance

1	Person Intercept 0.01997819 0.0003991281

2 Experiment Intercept 0.27406448 0.0751113402

3	Genotype Intercept 0.74659723 0.5574074167

Chisq	df	p	AIC BIC

Integrated loglik 24.14 4.00 7.474e-05 16.14 6.52

Penalized loglik 53.06 11.28 2.211e-07 30.51 3.37

Fixed effects:

coef exp(coef) se(coef)	z	p

LarvalDietPENSTRAS -0.1015	0.9034	0.2272 -0.45 0.655
