## Supplementary material for "Life history and infection susceptibility parameters of *Drosophila* species reared on a common diet": File S2: 00_FoodAnalysis.R

```
#####################################################################
# TITLE: An agar food recipe that rears most Drosophila species
# AUTHORS: Hongbo Sun†, Millie Cole†, Eduardo Ragnoni, Jodi Johnson, Alexander Jarvis, Léna Hédelin, Hannah Westlake, Ben Longdon, Mark A. Hanson*
### † These authors contributed equally to this study
### * Corresponding author:
#####################################################################
### DESCRIPTION: 
### Run this script, in order, to generate all analyses, tables, and figures in the paper
#####################################################################

######################
#### 0. Libraries ####
######################

# This script loads all packages, and installs them if necessary
source("Scripts/setup_PackageInstall_n_Load.R")

#####################
#### 0. Settings ####
#####################

# # This script defines things like colour palettes used throughout
# ### I could possibly add more colour palettes or set the default ggplot theme for all plots
# source("Scripts/setup_initial_settings.R")

# This script loads the initial phylogenetic tree that will be used in many figures
source("Scripts/setup_load_tree.R")

######################################
#### PART 1.1. Egg-to-adult times ####
######################################

source("Scripts/FigA_merged_df_prep.R")

# Here is the structure of merged_df
str(merged_df)
head(merged_df)
nrow(merged_df)
unique(merged_df$SpeciesName) # 66 species entering e2a

# #############################
# ## OPTIONAL: filter for only species with at least 2 observations before moving on ##
# merged_df <- merged_df %>%
#   group_by(SpeciesName) %>%
#   filter(sum(!is.na(EggToAdult)) >= 2) %>%
#   ungroup()
# #############################

source("Scripts/FigA_Egg2Adult.R")
# How many species are in the pruned_tree used here?
cat("Total species in tree:", length(pruned_tree$tip.label), "\n") # cat = concatenate strings
# Pagel's lambda on Egg2Adult
E2A_pagel_lambda

# OPTIONAL: re-run the phylo MCMCglmm to update the Rdata file ##
# run to generate new MCMC, takes a second
# source("Scripts/FigA_Run_E2A_phyloMCMCglmm.R")

# load the phylo MCMCglmm, save the summaries to a .txt file in the Figures folder
source("Scripts/FigA_Load_E2A_phyloMCMCglmm.R")

################################
#### PART 1.2. Wet/dry mass ####
################################
# ULTIMATELY LIKELY TO PUT WET MASS TREES IN SUPP, DRY MASS IN MAIN TEXT
# BOTH WET AND DRY MASS CAN BE PUT IN TABLE

####################################
#### Phylogenetic signal of mass
####################################

##############
### Females ##
source("Scripts/FigB_WetDryMass_F.R")
### For some reason the pagel's lambda seems really off...
# FWM_pagel_lambda # female wet mass

# ## OPTIONAL: re-run the phylo MCMCglmm to update the Rdata file ##
# source("Scripts/FigB_Run_FWM_phyloMCMCglmm.R")

## load the phylo MCMCglmm
source("Scripts/FigB_Load_FWM_phyloMCMCglmm.R")

##############
### Males ####

source("Scripts/FigB_WetDryMass_M.R")
### For some reason the pagel's lambda seems really off...
# MWM_pagel_lambda # male wet mass

# ## OPTIONAL: re-run the phylo MCMCglmm to update the Rdata file ##
# source("Scripts/FigB_Run_MWM_phyloMCMCglmm.R")

## load the phylo MCMCglmm
source("Scripts/FigB_Load_MWM_phyloMCMCglmm.R")

########
# assemble figure with magick
source("Scripts/FigA_phylogenies.R")

source("Scripts/FigA_Table1_combiner.R")

########
# Assess male-female correlation in body mass

source("Scripts/setup_load_tree.R") # reload the tree
### OPTIONAL: re-run the bivariate MCMCglmm for the correlation of species
# source("Scripts/FigBsupp_bivariate_species_corr.R") # just wet mass available for now
source("Scripts/FigBsupp_plot_WetMass_correlation.R") # just wet mass available for now

############################################
#### PART 2. Figure C, food comparisons ####
############################################

# ### pick a colour scheme! These are defined in the Script: "setup_initial_settings.R"
# FigC_colours <- c("PROPIONIC" = "#4F000B", "UPLEM" = "#720026", "PENSTRAS" = "#B93513", "CORN" = "#e27602", "PROP_YEAST" = "#4F000B")

### pick a colour scheme! These are defined in the Script: "setup_initial_settings.R"
FigC_colours <- c("Prop recipe" = "#4F000B", "EPFL corn" = "#720026", "Pen. corn" = "#B93513", "Cam. corn" = "#e27602", "Prop + yeast" = "#4F000B", "Prop + wick" = "#4F000B")

##
# Get comparisons of Corn, PENSTRAS, Prop across Dmel group species and some relatives
source("Scripts/FigCsupp1_Foods_E2A_analysis.R")

# Get comparisons of Corn, PENSTRAS, Prop across Dmel group species and some relatives
source("Scripts/FigC_yeastadd_barplot.R")

##
# main food comparison script
source("Scripts/FigC_Food_Comparisons.R")

# lipid floating stuff
source("Scripts/FigC_Food_LipidFloating.R")

# # the script below is an add-on that plots a stacked barplot by sex for n_emergents and saves it in z_not_used
# # I've made it optional because plotting this takes a bit of computing power, and it's not necessarily going to get used anyways since it's kinda ugly...
# source("Scripts/FigC_optional_stacked_plot.R")

# source("Scripts/FigC_WickAdd_barplot.R") # D. miranda wick addition to vials trends in positive direction for rearing success, but further replication needed (perhaps with longer egg-laying periods)

# source("Scripts/FigC_Magick_together.R") # lipid stuff on bottom
source("Scripts/FigC_Magick_together2.R") # lipid stuff in middle

######################################################
#### PART 3. Microbiota and Acetobacter infection ####
######################################################
## Need to add Millie's data as figures, which Eduardo has processed for the poster

source("Scripts/FigD_Mark_CFU_data.R")
source("Scripts/FigD_Pearson_Corr_MillieSurvs.R")
source("Scripts/FigD_PropCorn.R") 
source("Scripts/FigDsupp_PropCorn_fullspecies.R")
source("Scripts/FigD_Magick_together.R")

#######################################
#### PART 4. Providencia JH10 data ####
#######################################

source("Scripts/FigE_JH10_surv_data.R")
source("Scripts/FigE_suppAcetoPEN.R")
```
