## Supplementary material for "Life history and infection susceptibility parameters of *Drosophila* species reared on a common diet": File S2: Checking_Data.R

```
# seems like Hongbo's data is up to date

df1<-read_excel("Data_Raw/old/Nov8_VialWeights_JJ_BK1Pg80.xlsx") #, sheet = 1)
df2<-read_excel("Data_Raw/old/Nov14_VialWeights_JJ_BK1Pg80.xlsx") #, sheet = 1)
df1$PerFlyWetMass <- as.numeric(df1$PerFlyWetMass)
df1$PerFlyWetMass<-df1$PerFlyWetMass*1000 # convert g to mg
df2$PerFlyWetMass <- as.numeric(df2$PerFlyWetMass)
df2$PerFlyWetMass<-df2$PerFlyWetMass*1000 # convert g to mg

str(df1)
str(df2)
identical(df1, df2)
all.equal(df1, df2)

cols <- names(df1)

diff_index <- lapply(cols, function(col) {
  which(df1[[col]] != df2[[col]] |
          xor(is.na(df1[[col]]), is.na(df2[[col]])))
})

names(diff_index) <- cols
diff_index

merged_df <- df2
for (col in names(df2)) {
  merged_df[[col]] <- coalesce(df2[[col]], df1[[col]])
}

str(merged_df)

merged_df <- merged_df %>%
  mutate(across(where(is.character), ~ ifelse(is.na(.x), "", .x)))

str(merged_df)

write.csv(merged_df,"Data_Raw/old/MergedData_Bk1Pg80.csv", row.names = F)
library(writexl)
write_xlsx(merged_df, "Data_Raw/old/MergedData_Bk1Pg80.xlsx")

colSums(is.na(merged_df) & !is.na(df1))


# remove any rows where per fly wet mass isn't measured yet
df1_F <- df1 %>%
  filter(PerFlyWetMass > 0) %>%
  filter(NoPerTube > 4) %>% # 5 or more individuals per tube
  filter(Sex == "F")

# remove any rows where per fly wet mass isn't measured yet
df2_F <- df2 %>%
  filter(PerFlyWetMass > 0) %>%
  filter(NoPerTube > 4) %>% # 5 or more individuals per tube
  filter(Sex == "F")

# remove any rows where per fly wet mass isn't measured yet
df1_M <- df1 %>%
  filter(PerFlyWetMass > 0) %>%
  filter(NoPerTube > 4) %>% # 5 or more individuals per tube
  filter(Sex == "M")

# remove any rows where per fly wet mass isn't measured yet
df2_M <- df2 %>%
  filter(PerFlyWetMass > 0) %>%
  filter(NoPerTube > 4) %>% # 5 or more individuals per tube
  filter(Sex == "M")

str(df1_F)
str(df1_M)
str(df2_F)
str(df2_M)

length(unique(df1_F$SpeciesName))
length(unique(df2_F$SpeciesName))
setdiff(unique(df2_F$SpeciesName),unique(df2_F$SpeciesName))
length(unique(df1_M$SpeciesName))
length(unique(df2_M$SpeciesName))
setdiff(unique(df2_M$SpeciesName),unique(df2_M$SpeciesName))

# seems like Hongbo's data is up to date

###
# FOR LATER: can sub in fly weights instead of vial weights
###
summary_table_WM_F <- df1 %>%
  group_by(SpeciesGroup, SpeciesName) %>%
  summarise(
    MeanPerFlyWetMass = mean(PerFlyWetMass, na.rm = TRUE),
    SDPerFlyWetMass = sd(PerFlyWetMass, na.rm = TRUE),
    n_vials = sum(!is.na(PerFlyWetMass)),  # count of valid observations
    .groups = "drop"
  ) %>%
  filter(n_vials > 0) %>%  # drop rows with no observations
  # mutate(
  #   Mean_SD = sprintf("%.2f ± %.2f", MeanPerFlyWetMass, SDPerFlyWetMass)
  # ) %>%
  select(SpeciesName, SpeciesGroup, n_vials, MeanPerFlyWetMass, SDPerFlyWetMass) #SpeciesNumber useful if wanting to split D. suzu into two strains
summary_table_WM_F
```
