## Supplementary material for "Life history and infection susceptibility parameters of *Drosophila* species reared on a common diet": File S2: kePrint.js

```
$(document).ready(function(){
    if (typeof $('[data-toggle="tooltip"]').tooltip === 'function') {
        $('[data-toggle="tooltip"]').tooltip();
    }
    if ($('[data-toggle="popover"]').popover === 'function') {
        $('[data-toggle="popover"]').popover();
    }
});
```
