## Supplementary material for "Life history and infection susceptibility parameters of *Drosophila* species reared on a common diet": File S2: README.rtf

For analysis of Sun, Cole et al. (2025; bioRxiv)

The Rproject file included here uses 00_FoodAnalysis.R as its main file. Opening the R project, and then opening 00_FoodAnalysis.R should allow you to run all scripts and generate all figures in this study. The Figures folder contains two existing .png files to facilitate this, as one image is an illustration used in Figure 3, and another is a pilot survival series whose figure was generated in an independent program. The underlying data are nonetheless included in Data_Raw.
