## Supplementary material for "Life history and infection susceptibility parameters of *Drosophila* species reared on a common diet": File S2: FigA_Egg2Adult.R

```
#### Table A: Summary Table ####

# Which species were screened
unique(merged_df$SpeciesName) # 66 with 1+ observation, 61 with 2+ observations

# Make a summary table
summary_table_E2A <- merged_df %>%
  group_by(SpeciesName) %>%
  summarise(
    SpeciesGroup  = unique(SpeciesGroup),
    MeanEggToAdult = mean(EggToAdult, na.rm = TRUE),
    SDEggToAdult   = sd(EggToAdult, na.rm = TRUE),
    n_vials        = sum(!is.na(EggToAdult)),
    .groups = "drop"
  ) %>%
  filter(n_vials > 0) %>%  # drop rows with no observations
  select(SpeciesName, SpeciesGroup, n_vials, MeanEggToAdult, SDEggToAdult) #SpeciesNumber useful if wanting to split D. suzu into two strains

# print(summary_table_E2A, n = 100)
sort(summary_table_E2A$SpeciesName)

write.csv(summary_table_E2A, "Figures/BuildingBlocks/TableA_Egg2Adult_summary_table_E2A.csv", row.names = FALSE, fileEncoding = "Latin1") # row.names = FALSE avoids the extraneous 1st column


###
# Figure A: Egg-to-adult development times across the phylogeny (contMap colour mapped tree diagram)
###
# str(merged_df)

# 1. Calculate mean ignoring NAs
species_means <- merged_df %>%
  group_by(SpeciesName) %>%
  summarise(
    MeanEggToAdult = mean(EggToAdult, na.rm = TRUE),
    n_obs = sum(!is.na(EggToAdult)),
    SDEggToAdult = sd(EggToAdult, na.rm=TRUE))
head(species_means)

# 2. Remove rows where mean is NaN (happens if all values were NA)
species_means <- species_means %>%
  filter(!is.nan(MeanEggToAdult))
head(species_means)

# 3. consistent naming
tip_labels_cleaned <- tree$tip.label
head(tip_labels_cleaned)

# 4. Create named vector with cleaned species means
mean_vec <- species_means$MeanEggToAdult
names(mean_vec) <- species_means$SpeciesName
head(mean_vec)


# plot the tree
plot(tree, cex=0.6)
# tree$tip.label <- gsub("^(\\w+)\\.", "\\1. ", tree$tip.label)
missing <- setdiff(tree$tip.label, names(mean_vec))
# message("Missing tips: ", paste(missing, collapse = ", "))
pruned_tree <- drop.tip(tree, missing)
mean_vec_pruned <- mean_vec[pruned_tree$tip.label]
p_tree <- contMap(pruned_tree, mean_vec_pruned, legend = TRUE)
p_tree$legend <- "Egg to Adult (days)"  # <-- Custom label

# Ben's code
#V<-contMap(tree2,a1, res=200,fsize=c(0.5,0.9), plot=FALSE, legend = TRUE)

### inverse colours ###
p_tree$cols[]<-p_tree$cols[length(p_tree$cols):1]
plot(p_tree, fsize=c(.5,0.9))
plot(p_tree, legend = TRUE)

# Export tree plot to PNG
png("Figures/BuildingBlocks/FigA_Egg2Adult_tree_plot.png", width = 2400, height = 9000, res = 300)

# tree_height <- max(nodeHeights(pruned_tree))
# ylim <- c(0, tree_height * 1.15)  # 15% extra space above tips
plot(p_tree, legend = FALSE, fsize = c(2.5,0.9))

# This next bit places the color bar at top left manually, and allows me to round the digits... it's nice for customization, but you can use legend = TRUE above to just get the original legend
# In the future, maybe we can switch to a barplot

# Get the x/y limits of the plotting region
xl <- par("usr")[1:2]  # get plot window x-axis limits, needed to place x/y colour bar coordinates
yl <- par("usr")[3:4]  # get plot window x-axis limits, needed to place x/y colour bar coordinates

par(cex=1.8)
# add colour bar manually
add.color.bar(
  leg = 0.7425 * max(nodeHeights(pruned_tree)),  # length of the bar, this value makes it "0.55" which is inoffensive
  cols = p_tree$cols,
  lims = range(mean_vec_pruned),
  digits = 0,
  title = "Egg-to-adult",
  prompt = FALSE,
  lwd = 6,
  ltext = NA,
  x = xl[1] + 0.05 * diff(xl),  # slightly right of left edge
  y = yl[2] - 0.03 * diff(yl),  # slightly down from top edge
  horizontal = TRUE             # make bar horizontal
)
# text(x = 0.2, y = -1, labels = "Egg to Adult (days)", xpd = NA, cex = 1, font = 2)
dev.off()


###
# TABLE TIME
###

# Convert SpeciesName to character just in case
summary_table_E2A <- species_means %>% mutate(SpeciesName = as.character(SpeciesName))
# str(summary_table_E2A)

# reorder columns, rename for table
summary_table_E2A_ordered <- summary_table_E2A %>%
  rename(n_vials = n_obs) %>%     # replace column name
  select(SpeciesName, n_vials, MeanEggToAdult, SDEggToAdult)

# centre
summary_kable <- summary_table_E2A_ordered %>%
  select(SpeciesName, n_vials, MeanEggToAdult, SDEggToAdult) %>%
  kable(align = "cc") %>%  # 'c' = center alignment for each column
  kable_styling(position = "center")

# Save the table as an HTML
save_kable(summary_kable, "Figures/BuildingBlocks/TableA_summary_table_E2A.html")

###
#############
# Phylo signal
#############
###
# Quick Pagel's Lambda
###

# Remove rows with NA in EggToAdult
E2A_subset_df <- merged_df[!is.na(merged_df$EggToAdult), ]
# Match only species in both tree and data
common_species <- intersect(pruned_tree$tip.label, E2A_subset_df$SpeciesName)
# Prune tree and trait vector
tree_pruned <- drop.tip(pruned_tree, setdiff(pruned_tree$tip.label, common_species))
# setNames() step required because phylosig needs a named numeric vector
E2A_vector <- setNames(E2A_subset_df$EggToAdult, E2A_subset_df$SpeciesName)[common_species]
plot(tree_pruned, cex = 0.6)

# Pagel's lambda stored as E2A_pagel_lambda
E2A_pagel_lambda<-phylosig(tree_pruned, E2A_vector, method = "lambda", test = TRUE)
```
