## Supplementary material for "Life history and infection susceptibility parameters of *Drosophila* species reared on a common diet": File S2: FigA_Load_E2A_phyloMCMCglmm.R

```
##########
# E2A phylogenetic heritability
##########

load("Data_Clean/Rdata/E2A_PhyloGlmm_WithSpecies.Rdata")
E2A_mcmc <- E2A_PhyloGlmm_WithSpecies   # shorthand

# --- Extract posterior samples ---
animal  <-E2A_mcmc$VCV[, "animal"]
species <-E2A_mcmc$VCV[, "SpeciesName"]
resid   <-E2A_mcmc$VCV[, "units"]

# define phylogenetic partition equations
h2_phy     <- animal / (animal+species)
h2_total     <- animal / (animal+species+resid)

# wrap in a helper function to summarise multiple definitions
summarise_post <- function(x){
  mcmc_x <- as.mcmc(x)
  list(
    autocorr  = autocorr.diag(mcmc_x),
    ESS       = effectiveSize(mcmc_x),
    mean      = mean(x),
    sd        = sd(x),
    quantiles = quantile(x, probs = c(0.025,0.05,0.25,0.5,0.75,0.95,0.975)),
    HPD       = HPDinterval(mcmc_x)
  )
}

s_animal  <- summarise_post(animal)
s_species <- summarise_post(species)
s_resid   <- summarise_post(resid)
s_h2phy   <- summarise_post(h2_phy)
s_h2total   <- summarise_post(h2_total)

# --- Output to file ---
sink("Figures/Fig2_E2A_phylogenetic heritability.txt")

cat("\n====================\n1. Pagel's lambda\n====================\n")
print(E2A_pagel_lambda)
cat("\n====================\n2. Inspect the variance of phy, spp, and residuals\n====================\n\n")
cat("\n--- animal (phylogenetic variance) ---\n")
print(s_animal)
cat("\n--- SpeciesName (species variance) ---\n")
print(s_species)
cat("\n--- units (residual variance) ---\n")
print(s_resid)
cat("\n====================\n3. Phylogenetic heritability\n====================\n\n")
cat("\n=== vp/vp+vs ===\n\n")
print(s_h2phy)
cat("\n=== vp/vp+vs+ve ===\n\n")
print(s_h2total)
sink()
```
