## Supplementary material for "Life history and infection susceptibility parameters of *Drosophila* species reared on a common diet": File S2: FigA_merged_df_prep.R

```
# The way the data were entered is not aesthetic for R to manage, so we've dedicated a script to preparing the dataframe merged_df

## Note: df1 was JJ first formal attempt to collect E2A and many outliers in this dataframe relative to all others (and Mark's confirmed observations)
## For safety's sake, censoring df1
df1<-read_excel("Data_Raw/Egg2Adult_JJ_Bk1Pg80.xlsx", sheet = 1) # JJ block 1 observations, likely censor
df2<-read_excel("Data_Raw/Egg2Adult_JJ_Bk1Pg80.xlsx", sheet = 2) # JJ block 2 observations
df3<-read_excel("Data_Raw/Egg2Adult_JJ_Bk1Pg80.xlsx", sheet = 3) # JJ block 3 observations
df4<-read_excel("Data_Raw/Egg2Adult_JJ_Bk1Pg80.xlsx", sheet = 4) # JJ block 4 observations
df5<-read_excel("Data_Raw/Egg2Adult_JJ_Bk1Pg80.xlsx", sheet = 5) # Mark's observations

df1 <- df1 %>%
  mutate(EmergentsStart = as.Date(as.numeric(EmergentsStart), origin = "1899-12-30"))

df2 <- df2 %>%
  mutate(EmergentsStart = as.Date(as.numeric(EmergentsStart), origin = "1899-12-30"))

df3 <- df3 %>%
  mutate(EmergentsStart = as.Date(as.numeric(EmergentsStart), origin = "1899-12-30"))

df4 <- df4 %>%
  mutate(EmergentsStart = as.Date(as.numeric(EmergentsStart), origin = "1899-12-30"))

df5 <- df5 %>%
  mutate(EmergentsStart = as.Date(as.numeric(EmergentsStart), origin = "1899-12-30"),
         Egg2Adult = as.numeric(Egg2Adult))

# Standardize and select relevant columns for each dataframe
df1_clean <- df1 %>%
  select(SpeciesNumber, SpeciesName, SpeciesNameShort, SpeciesGroup, Block, Egg2Adult) %>%
  rename(EggToAdult = Egg2Adult)
df1_clean$Person = "JJ"

df2_clean <- df2 %>%
  select(SpeciesNumber, SpeciesName, SpeciesNameShort, SpeciesGroup, Block, Egg2Adult) %>%
  rename(EggToAdult = Egg2Adult)
df2_clean$Person = "JJ"

df3_clean <- df3 %>%
  select(SpeciesNumber, SpeciesName, SpeciesNameShort, SpeciesGroup, Block, Egg2Adult) %>%
  rename(EggToAdult = Egg2Adult)
df3_clean$Person = "JJ"

df4_clean <- df4 %>%
  select(SpeciesNumber, SpeciesName, SpeciesNameShort, SpeciesGroup, Block, Egg2Adult) %>%
  rename(EggToAdult = Egg2Adult)
df4_clean$Person = "JJ"

df5_clean <- df5 %>%
  filter(!is.na(Egg2Adult)) %>%
  select(SpeciesNumber, SpeciesName, SpeciesNameShort, SpeciesGroup, Block, Egg2Adult) %>%
  rename(EggToAdult = Egg2Adult)
df5_clean$Person = "MAH"

# Combine all into one dataframe
# merged_df <- bind_rows(df1_clean, df2_clean, df3_clean, df4_clean, df5_clean)
merged_df <- bind_rows(df2_clean, df3_clean, df4_clean, df5_clean) # Suspect these 1st observations by JJ were not accurate (many purported to be days longer than minimum) 
# merged_df <- bind_rows(df2_clean, df3_clean, df5_clean) # df1 and df4 are the biggest outliers, see phylogenetic signal with df2, 3, 5

write.csv(merged_df, "Data_Clean/Egg2Adult_merged_df.csv", row.names = FALSE) # row.names = FALSE avoids the extraneous 1st column
```
