## Supplementary material for "Life history and infection susceptibility parameters of *Drosophila* species reared on a common diet": File S2: FigA_Phylogenies.R

```
##############
# stick em together
##############
library(magick)

# Read images
FigA <- image_read("Figures/BuildingBlocks/FigA_Egg2Adult_tree_plot.png")
FigB <- image_read("Figures/BuildingBlocks/FigB_FWM_tree_plot.png")
FigC <- image_read("Figures/BuildingBlocks/FigB_MWM_tree_plot.png")

# Combine top + spacer + top
Fig2_bottom <- c(FigA, FigB, FigC) %>%
  image_append(stack = FALSE)   # TRUE -> vertical

# Create a transparent spacer (adjust height as needed)
spacer_height <- 100  # pixels
spacer <- image_blank(width = image_info(Fig2_bottom)$width, 
                      height = spacer_height, 
                      color = "none")  # transparent

# Combine top + spacer + top
Fig2_plot <- c(spacer, Fig2_bottom) %>%
  image_append(stack = TRUE)   # TRUE -> vertical

# Save plot collection image
image_write(Fig2_plot, path = "Figures/BuildingBlocks/Fig2_Combined.png", format = "png")

library(png)

# Load the image
img <- readPNG("Figures/BuildingBlocks/Fig2_Combined.png")
img_dim <- dim(img)

# Open PNG device (same size as image)
png("Figures/Fig2_Combined_Labeled.png", width = img_dim[2], height = img_dim[1], bg = "transparent") # bg = "transparent" prevents the hairline border being drawn alongside the par expression below
par(mar = c(0, 0, 0, 0), xaxs = "i", yaxs = "i") # Remove margins and set exact axis limits

# Plot blank canvas
plot(1, type = "n", xlim = c(0, img_dim[2]), ylim = c(0, img_dim[1]),
     xaxt = 'n', yaxt = 'n', xlab = '', ylab = '', asp = 1)

# Draw the image
rasterImage(img, 0, 0, img_dim[2], img_dim[1])

# Add labels
text(x = 100,  y = 9000, labels = "A", cex = 14, font = 2)
text(x = 2400,  y = 9000, labels = "B", cex = 14, font = 2)
text(x = 4800, y = 9000, labels = "C", cex = 14, font = 2)

# Close device to save
dev.off()
```
