## Supplementary material for "Life history and infection susceptibility parameters of *Drosophila* species reared on a common diet": File S2: FigA_Table1_combiner.R

```
# Simplify the Egg-to-Adult table
E2A_simple <- summary_table_E2A_ordered %>%
  mutate(EggToAdult = paste0(round(MeanEggToAdult, 2), " ± ", 
                             round(SDEggToAdult, 2), 
                             " (", n_vials, ")")) %>%
  select(SpeciesName, EggToAdult)

# Simplify the Wet Mass (Males) table
WM_M_simple <- summary_table_WM_M_ordered %>%
  mutate(WetMass_M = paste0(round(MeanPerFlyWetMass, 2), " ± ", 
                            ifelse(is.na(SDPerFlyWetMass), "NA", round(SDPerFlyWetMass, 2)), 
                            " (", n_vials, ")")) %>%
  select(SpeciesName, WetMass_M)

# Simplify the Wet Mass (Females) table
WM_F_simple <- summary_table_WM_F_ordered %>%
  mutate(WetMass_F = paste0(round(MeanPerFlyWetMass, 2), " ± ", 
                            ifelse(is.na(SDPerFlyWetMass), "NA", round(SDPerFlyWetMass, 2)), 
                            " (", n_vials, ")")) %>%
  select(SpeciesName, WetMass_F)

# Merge all three simplified tables by SpeciesName
Table1_combined <- E2A_simple %>%
  full_join(WM_F_simple, by = "SpeciesName") %>%
  full_join(WM_M_simple, by = "SpeciesName") %>%
  arrange(SpeciesName)

# Print combined summary table
Table1_combined
Table1_names<-c("Species","Egg to adult","Female wet mass","Male wet mass")
colnames(Table1_combined) <- Table1_names

write.csv(Table1_combined,"Figures/Table1_combined.csv",row.names = FALSE)

# centre
summary_kable <- Table1_combined %>%
  select(Species, `Egg to adult`,`Female wet mass`,`Male wet mass`) %>%
  kable(align = "cc") %>%  # 'c' = center alignment for each column
  kable_styling(position = "center")

# Save the table as an HTML
save_kable(summary_kable, "Figures/Table1_combined.html")
```
