## Supplementary material for "Life history and infection susceptibility parameters of *Drosophila* species reared on a common diet": File S2: FigB_Run_MWM_phyloMCMCglmm.R

```
###
# Mark's quick phylo.glmm / MCMCglmm
###

# Make 'animal' a factor in E2A_subset_df, and make the tibble into a dataframe
MWM_subset_df$animal <- factor(as.character(MWM_subset_df$SpeciesName))
MWM_subset_df <- as.data.frame(MWM_subset_df)

str(MWM_subset_df)

# Model 1, with species

### Priors seetings for phylogenetic mixed models
prior_with_species <- list(G = list(
  G1 = list(V = diag(1), nu = 1, alpha.mu = rep(0,1), alpha.V = diag(1) * 1000),
  G2 = list(V = diag(1), nu = 1, alpha.mu = rep(0,1), alpha.V = diag(1) * 1000)),
  R = list(V = diag(1), nu = 0.002))

### MCMC
MWM_PhyloGlmm_WithSpecies<- MCMCglmm(
  PerFlyWetMass ~ 1,
  random = ~animal + SpeciesName,
  rcov = ~units, # idh, independent heterogenous, factors inside idh have own variance
  pedigree = pruned_tree,
  prior = prior_with_species,
  data = MWM_subset_df,
  nitt = 13000000, # add a 0 later
  thin = 5000, # add a 0 later
  burnin = 3000000, # add a 0 later
  pr = TRUE
)
save(MWM_PhyloGlmm_WithSpecies, file = "Data_Clean/Rdata/MWM_PhyloGlmm_WithSpecies.Rdata")
```
