## Supplementary material for "Life history and infection susceptibility parameters of *Drosophila* species reared on a common diet": File S2: FigB_WetDryMass_F.R

```
# this script generates a tree output for the wet and dry mass for females accompanying the experiments shown in FigA

df1<-read_excel("Data_Raw/VialWeights_JJ_BK1Pg80.xlsx") #, sheet = 1)
df1$PerFlyWetMass <- as.numeric(df1$PerFlyWetMass)
# df1$PerFlyWetMass<-df1$PerFlyWetMass*1000 # convert g to mg

# remove any rows where per fly wet mass isn't measured yet
df1 <- df1 %>%
  filter(PerFlyWetMass > 0) %>%
  filter(NoPerTube > 0) %>% # 1 or more individuals per tube
  filter(Sex == "F")

###
summary_table_WM_F <- df1 %>%
  group_by(SpeciesGroup, SpeciesName) %>%
  summarise(
    MeanPerFlyWetMass = mean(PerFlyWetMass, na.rm = TRUE),
    SDPerFlyWetMass = sd(PerFlyWetMass, na.rm = TRUE),
    n_vials = sum(!is.na(PerFlyWetMass)),  # count of valid observations
    .groups = "drop"
  ) %>%
  filter(n_vials > 0) %>%  # drop rows with no observations
  # mutate(
  #   Mean_SD = sprintf("%.2f ± %.2f", MeanPerFlyWetMass, SDPerFlyWetMass)
  # ) %>%
  select(SpeciesName, SpeciesGroup, n_vials, MeanPerFlyWetMass, SDPerFlyWetMass) #SpeciesNumber useful if wanting to split D. suzu into two strains
summary_table_WM_F
sort(unique(summary_table_WM_F$SpeciesName))

write.csv(summary_table_WM_F, "Figures/BuildingBlocks/TableB_FWM_summary_table.csv", row.names = FALSE, fileEncoding = "Latin1")

###
# PLOT
###

# 1. Calculate mean ignoring NAs
species_means <- df1 %>%
  group_by(SpeciesName) %>%
  summarise(MeanPerFlyWetMass = mean(PerFlyWetMass, na.rm = TRUE),
            n_vials = sum(!is.na(PerFlyWetMass)),
            SDPerFlyWetMass = sd(PerFlyWetMass, na.rm=TRUE)) %>%
  ungroup()
head(species_means)

# 2. Remove rows where mean is NaN (happens if all values were NA)
species_means <- species_means %>%
  filter(!is.nan(MeanPerFlyWetMass))
head(species_means)

# 3. consistent naming
tip_labels_cleaned <- tree$tip.label
head(tip_labels_cleaned)

# 4. Create named vector with cleaned species means
mean_vec <- species_means$MeanPerFlyWetMass
names(mean_vec) <- species_means$SpeciesName
head(mean_vec)

# plot the tree
plot(tree, cex=0.6)
# tree$tip.label <- gsub("^(\\w+)\\.", "\\1. ", tree$tip.label)
missing <- setdiff(tree$tip.label, names(mean_vec))
# message("Missing tips: ", paste(missing, collapse = ", "))
pruned_tree <- drop.tip(tree, missing)
mean_vec_pruned <- mean_vec[pruned_tree$tip.label]
p_tree <- contMap(pruned_tree, mean_vec_pruned, legend = TRUE)
p_tree$legend$title <- "Wet mass"  # <-- Change title here
plot(p_tree, legend = TRUE)

### inverse colours ###
p_tree$cols[]<-p_tree$cols[length(p_tree$cols):1]
plot(p_tree, fsize=c(0.5,0.9))
plot(p_tree, legend = TRUE)

# Export tree plot to PNG
png("Figures/BuildingBlocks/FigB_FWM_tree_plot.png", width = 2400, height = 9000, res = 300)
plot(p_tree, legend = FALSE, fsize = c(2.5,0.9))

# This next bit places the color bar at top left manually, and allows me to round the digits... it's nice for customization, but you can use legend = TRUE above to just get the original legend

# Get the x/y limits of the plotting region
xl <- par("usr")[1:2]  # get plot window x-axis limits, needed to place x/y colour bar coordinates
yl <- par("usr")[3:4]  # get plot window x-axis limits, needed to place x/y colour bar coordinates

par(cex=1.6)
# add colour bar manually
add.color.bar(
  leg = 0.7425 * max(nodeHeights(pruned_tree)),  # length of the bar, this value makes it "0.55" which is inoffensive
  cols = p_tree$cols,
  lims = range(mean_vec_pruned),
  digits = 2,
  title = "Mass F (mg)",
  prompt = FALSE,
  lwd = 4,
  ltext = NA,
  x = xl[1] + 0.05 * diff(xl),  # slightly right of left edge
  y = yl[2] - 0.03 * diff(yl),  # slightly down from top edge
  horizontal = TRUE             # make bar horizontal
)
# text(x = 0.2, y = -1, labels = "Egg to Adult (days)", xpd = NA, cex = 1, font = 2)
dev.off()


###
# TABLE TIME
###

# # redo depending on species included
# tip_label_order <- c("D. sturtevanti", "D. saltans", "D. prosaltans", "D. tropicalis", "D. willistoni", "D. sucinea", "D. nebulosa", "D. ananassae", "D. teisseri", "D. yakuba", "D. santomea", "D. erecta", "D. melanogaster", "D. simulans", "D. sechellia", "D. mauritiana", "D. takahashii", "D. suzukii", "D. baimaii", "D. subobscura", "D. bifasciata", "D. affinis", "D. miranda", "D. pseudoobscura", "D. persimilis", "D. paramelanica", "D. euronotus", "D. micromelanica", "D. arizonae", "D. mojavensis", "D. buzzatii", "D. hydei", "D. lummei", "D. novamexicana", "D. americana", "D. virilis", "D. littoralis", "D. lacicola", "D. flavomontana", "D. montana", "Z. inermis", "Z. tuberculatus", "Z. taronus", "D. brachynephros", "D. phalerata", "D. nigromaculata", "D. guttifera", "D. deflecta", "D. munda", "D. funebris", "D. bizonata", "D. testacea", "D. putrida", "D. immigrans", "D. nasuta", "D. sulfurigaster", "H. duncani", "S. lebanonensis") # "S. pattersoni"
# # 59 total in tip_label_order
# unique(pruned_tree$tip.label) # 58 total in pruned_tree$tip.label

# Convert SpeciesName to character just in case
summary_table_WM_F <- species_means %>% mutate(SpeciesName = as.character(SpeciesName))
str(summary_table_WM_F)

# reorder columns, rename for table
summary_table_WM_F_ordered <- summary_table_WM_F %>%
  select(SpeciesName, n_vials, MeanPerFlyWetMass, SDPerFlyWetMass)

# centre
summary_kable <- summary_table_WM_F_ordered %>%
  select(SpeciesName, n_vials, MeanPerFlyWetMass, SDPerFlyWetMass) %>%
  kable(align = "cc") %>%  # 'c' = center alignment for each column
  kable_styling(position = "center")

# Save the table as an HTML
save_kable(summary_kable, "Figures/BuildingBlocks/TableB_FWM_summary_table.html")


# ###
# #############
# # Quick phylo signal
# #############
# ###
# # Quick Pagel's Lambda
# ###
# 
# # Remove rows with NA in EggToAdult
# FWM_subset_df <- df1[!is.na(df1$PerFlyWetMass), ]
# # Match only species in both tree and data
# common_species <- intersect(pruned_tree$tip.label, FWM_subset_df$SpeciesName)
# # Prune tree and trait vector
# tree_pruned <- drop.tip(pruned_tree, setdiff(pruned_tree$tip.label, common_species))
# # setNames() step required because phylosig needs a named numeric vector
# E2A_vector <- setNames(FWM_subset_df$PerFlyWetMass, FWM_subset_df$SpeciesName)[common_species]
# plot(tree_pruned, cex = 0.6)
# 
# # Pagel's lambda stored as E2A_pagel_lambda
# FWM_vec <- setNames(FWM_subset_df$PerFlyWetMass, FWM_subset_df$SpeciesName)
# FWM_pagel_lambda<-phylosig(tree_pruned, FWM_vec, method = "lambda", test = TRUE)
# FWM_pagel_lambda
```
