## Supplementary material for "Life history and infection susceptibility parameters of *Drosophila* species reared on a common diet": File S2: FigBsupp_bivariate_species_corr.R

```
# load data
df1<-read_excel("Data_Raw/VialWeights_JJ_BK1Pg80.xlsx") #, sheet = 1)
df1$animal<-df1$SpeciesName
df1 <- df1 %>%
  mutate(Sex = toupper(trimws(Sex))) %>%
  filter(Sex %in% c("M", "F"))   # "f" → "F"
# remove NAs and autofilled cells where wet mass < 0
df1 <- df1 %>%
  filter(!is.na(PerFlyWetMass), PerFlyWetMass > 0)
str(df1)
df1.1<-df1 %>%
  select(SpeciesName,Sex,Block,PerFlyWetMass,animal)
df1.1$block<-as.factor(df1.1$Block)
df1.1$sex<-as.factor(df1.1$Sex)
df1.1$spp<-as.factor(df1.1$SpeciesName)
unique(df1$Sex)
unique(df1$Block)
# create new df where columns are species, wmm, wmf
df2 <- df1 %>%
  transmute(
    spp = SpeciesName,
    animal = animal,
    date = Block,
    Sex,
    wmass = PerFlyWetMass
  ) %>%
  # one row per date × animal × spp × sex
  group_by(date, animal, spp, Sex) %>%
  summarise(wmass = mean(wmass, na.rm = TRUE), .groups = "drop") %>%
  # wide format: wm_m, wm_f
  pivot_wider(
    names_from = Sex,
    values_from = wmass,
    names_prefix = "wm_"
  )

df2$date<-as.factor(df2$date)
df2$animal<-as.factor(df2$animal)
df2$spp<-as.factor(df2$spp)
str(df2)

# sanity checks, how does this filtering summarise a species like D. ananassae
df1_Dana <- df1[df1$SpeciesName=="D. ananassae",]
print(df1_Dana,n=30)
df2_Dana <- df2[df2$spp=="D. ananassae",]
df2_Dana

head(df2)


# tree bits
str(tree)
# 3. consistent naming
tip_labels_cleaned <- tree$tip.label
head(tip_labels_cleaned)

# 4. Create named vector with cleaned species means
df3 <- df2 %>%
  filter(!is.na(df2$wm_M))
wetmass_M <- df3$wm_M
names(wetmass_M) <- df3$spp
head(wetmass_M)

# plot the tree as a sanity check that the tips all map
plot(tree, cex=0.6)
# tree$tip.label <- gsub("^(\\w+)\\.", "\\1. ", tree$tip.label)
missing <- setdiff(tree$tip.label, names(wetmass_M))
# message("Missing tips: ", paste(missing, collapse = ", "))
pruned_treeM <- drop.tip(tree, missing)
plot(pruned_treeM) # check
wetmass_M_pruned <- wetmass_M[pruned_treeM$tip.label]
p_tree <- contMap(pruned_tree, wetmass_M_pruned, legend = TRUE)
p_tree$legend <- "Egg to Adult (days)"  # <-- Custom label


# 4. Create named vector with cleaned species means
df4 <- df2 %>%
  filter(!is.na(df2$wm_F))
wetmass_F <- df4$wm_F
names(wetmass_F) <- df4$spp
head(wetmass_F)

# plot the tree as a sanity check that the tips all map
plot(tree, cex=0.6)
# tree$tip.label <- gsub("^(\\w+)\\.", "\\1. ", tree$tip.label)
missing <- setdiff(tree$tip.label, names(wetmass_F))
# message("Missing tips: ", paste(missing, collapse = ", "))
pruned_treeF <- drop.tip(tree, missing)
plot(pruned_treeF) # check
wetmass_F_pruned <- wetmass_F[pruned_treeF$tip.label]
p_tree <- contMap(pruned_treeF, wetmass_F_pruned, legend = TRUE)
p_tree$legend <- "Egg to Adult (days)"  # <-- Custom label


# check the tree for the whole dataset
str(tree)
# tree$tip.label <- gsub("^(\\w+)\\.", "\\1. ", tree$tip.label)
missing <- setdiff(tree$tip.label, df2$animal)
# message("Missing tips: ", paste(missing, collapse = ", "))
pruned_tree_wm <- drop.tip(tree, missing)
plot(pruned_tree_wm) # check


# result so far
str(wetmass_M) # vector of wm_M column
str(wetmass_F) # vector of wm_F column
str(df1)
str(df1.1)
str(df2)
str(df3) # removed all rows with male NAs
str(df4) # removed all rows with female NAs
str(pruned_tree_wm)

# TO AVOID THE GINVERSE() ISSUE, MAKE SURE YOUR DF IS A DATAFRAME AND NOT A TIBBLE
df1.1<-as.data.frame(df1.1)

# Define priors
prior.with.species1 <- list(
  G = list(
    # G1 = animal: unstructured 2x2 (sex has 2 levels)
    G1 = list(
      V = diag(2),
      nu = 3,               # slightly > dimension (2) -> diffuse IW prior, nu = dimension + nu so 2x2 = 2 + weak weighting
      alpha.mu = rep(0, 2),     # parameter-expansion mean
      alpha.V = diag(2) * 1000   # large variance for parameter-expansion
    ),
    # G2 = spp: also unstructured 2x2
    G2 = list(
      V = diag(2),
      nu = 3,
      alpha.mu = rep(0, 2),
      alpha.V = diag(2) * 1000
    )
  ),
  # R: idh(sex):units -> diagonal residual matrix with 2 entries
  R = list(
    V = diag(2),  # residual scale (start with somewhat smaller scale)
    nu = 0.002                      # very diffuse residual prior
  )
)

#
phylo.glmm.wetmass <- MCMCglmm(
  PerFlyWetMass ~ sex, ##
  random = ~us(sex):animal + us(sex):spp, #random effect with phylogeny and specie not-connected to phylogeny
  rcov = ~idh(sex):units, #formula for residue variance defined by every spot in the data tghat is associated with a residue ~idh(trait):units
  pedigree = pruned_tree_wm, #tree arvore filogenética
  prior = prior.with.species1,
  data = df1.1, #data input Input de dados
  nitt = 13000000, #Iteration number Numero de iterações
  thin = 5000, # Sample frequency Frequência de amostragem
  burnin = 3000000, #Calibration Iterações antes de usar (Calibragem)
  pr = TRUE #Save posterior distribution of random effect Salvar a distribuição posterior dos efeitos aleatórios
)

save(phylo.glmm.wetmass, file = "Data_Clean/Rdata/phylo.glmm.wetmass.Rdata")

# check results
load("Data_Clean/Rdata/phylo.glmm.wetmass.Rdata")
wm_mcmc <- phylo.glmm.wetmass   # shorthand
summary(wm_mcmc)

# Phylogenetic (animal) covariance
animal_FF <- wm_mcmc$VCV[, "sexF:sexF.animal"]
animal_MM <- wm_mcmc$VCV[, "sexM:sexM.animal"]
animal_FM <- wm_mcmc$VCV[, "sexF:sexM.animal"]  # same as sexM:sexF.animal

# Non-phylogenetic species-level covariance
species_FF <- wm_mcmc$VCV[, "sexF:sexF.spp"]
species_MM <- wm_mcmc$VCV[, "sexM:sexM.spp"]
species_FM <- wm_mcmc$VCV[, "sexF:sexM.spp"]

# Residuals
resid_F <- wm_mcmc$VCV[, "sexF.units"]
resid_M <- wm_mcmc$VCV[, "sexM.units"]
resid_FM <- 0  # no residual covariance between sexes

summarise_vector <- function(x) {
  x <- as.numeric(x)
  mcmc_x <- as.mcmc(x)
  list(
    mean = mean(mcmc_x),
    sd   = sd(mcmc_x),
    quantiles = quantile(mcmc_x, probs = c(0.025, 0.5, 0.975)),
    HPD  = HPDinterval(mcmc_x),
    ESS  = effectiveSize(mcmc_x)
  )
}

# correlation of VCV
#colnames(model)
# select VCV columns of interest

# autocorr < .01
# 95% CI, defined in methods as HPD

### Inter-species repeatability
# phylo + species/ phylo + species + resid (units)

### correlation statistics
# vp/vp+vs
r_phy_adjusted_FM <- animal_FM / sqrt( (animal_FF + species_FF) * (animal_MM + species_MM) )
summary_r_phy_adjusted <- summarise_vector(r_phy_adjusted_FM)

# vp/vp+vs+ve
r_phy_over_total_FM <- animal_FM / sqrt( (animal_FF + species_FF + resid_F) * (animal_MM + species_MM + resid_M) )
summary_r_phy_over_total <- summarise_vector(r_phy_over_total_FM)

# (vp/vp+vs)^2
r2_phy_adjusted_FM <- r_phy_adjusted_FM^2
summary_r2_phy_adjusted <- summarise_vector(r2_phy_adjusted_FM)


sink("Figures/Fig2supp_wetmass_mf_correlation.txt")

cat("\n====================\n1. Pagel's lambda\n====================\n")
cat("\nmale\n")
print(MWM_pagel_lambda)
cat("\nfemale\n")
print(FWM_pagel_lambda)

cat("\n====================\n2. Male:Female correlations across species\n====================\n\n")

cat("\n======= Correlation of M:F wet mass across species =======\n\n")
cat("\n=== equivalent of rho for:  phy / phy + spp ===\n\n")
print(summary_r_phy_adjusted)
cat("\n=== equivalent of rho for: phy / phy + spp + resid ===\n\n")
print(summary_r_phy_over_total)
cat("\n=== equivalent of rho^2 for: phy / phy + spp ===\n\n")
print(summary_r2_phy_adjusted)
sink()


########
# THIS STUFF BELOW COULD BE USEFUL
########


# # --- Extract posterior samples ---
# colnames(wm_mcmc$VCV)
# animal <- wm_mcmc$VCV[, c(
#   "sexF:sexF.animal",
#   "sexM:sexF.animal",
#   "sexF:sexM.animal",
#   "sexM:sexM.animal"
# )]
# species <- wm_mcmc$VCV[, c(
#   "sexF:sexF.spp",
#   "sexM:sexF.spp",
#   "sexF:sexM.spp",
#   "sexM:sexM.spp"
# )]
# resid <- wm_mcmc$VCV[, c(
#   "sexF.units",
#   "sexM.units"
# )]
# 
# 
# 
# # define phylogenetic partition equations
# h2_phy     <- animal / (animal+species)
# h2_total     <- animal / (animal+species+resid)
# 
# # wrap in a helper function to summarise multiple definitions
# 
# # helper function to summarise
# summarise_post <- function(x) {
#   # If x is a vector → convert to a 1-column matrix
#   if (is.vector(x)) {
#     x <- matrix(x, ncol = 1)
#     colnames(x) <- "value"
#   }
#   # Apply the summary to each column
#   results <- lapply(seq_len(ncol(x)), function(i) {
#     col <- x[, i]
#     mcmc_col <- as.mcmc(col)
#     list(
#       autocorr  = autocorr.diag(mcmc_col),
#       ESS       = effectiveSize(mcmc_col),
#       mean      = mean(col),
#       sd        = sd(col),
#       quantiles = quantile(col, probs = c(0.025,0.05,0.25,0.5,0.75,0.95,0.975)),
#       HPD       = HPDinterval(mcmc_col)
#     )
#   })
#   names(results) <- colnames(x)
#   return(results)
# }
# 
# 
# # summarise posterior distributions
# s_animal    <- summarise_post(animal)
# s_species   <- summarise_post(species)
# s_resid     <- summarise_post(resid)
# s_h2phy     <- summarise_post(h2_phy)
# s_h2total   <- summarise_post(h2_total)
# 
# # output
# sink("Figures/Fig2_wetmass_phylogenetic heritability.txt")
# 
# cat("\n====================\n1. Pagel's lambda\n====================\n")
# cat("\nmale\n")
# print(MWM_pagel_lambda)
# cat("\nfemale\n")
# print(FWM_pagel_lambda)
# 
# cat("\n====================\n2. Inspect the variance of phy, spp, and residuals\n====================\n")
# 
# cat("\n--- animal (phylogenetic variance) ---\n")
# print(s_animal)
# 
# cat("\n--- spp (species variance) ---\n")
# print(s_species)
# 
# cat("\n--- units (residual variance) ---\n")
# print(s_resid)
# 
# cat("\n====================\n3. Phylogenetic heritability\n====================\n")
# 
# cat("\n=== vp / (vp + vs) ===\n\n")
# print(s_h2phy)
# 
# cat("\n=== vp / (vp + vs + ve) ===\n\n")
# print(s_h2total)
# 
# sink()
```
