## Supplementary material for "Life history and infection susceptibility parameters of *Drosophila* species reared on a common diet": File S2: FigBsupp_plot_WetMass_correlation.R

```
library(dplyr)
library(lme4)
library(lmerTest)  # for p-values
library(ggrepel)

# load data
df1<-read_excel("Data_Raw/VialWeights_JJ_BK1Pg80.xlsx") #, sheet = 1)
# remove any rows where per fly wet mass isn't measured yet
df1 <- df1 %>%
  filter(PerFlyWetMass > 0) %>%
  filter(NoPerTube > 0) # 1 or more individuals per tube

# Make sure factors are factors
df1 <- df1 %>%
  mutate(Sex = factor(Sex),
         SpeciesName = factor(SpeciesName),
         Block = factor(Block)
  )

str(df1)

###
# plotting time

df_plot <- df1 %>%
  group_by(SpeciesName, Sex) %>%
  summarise(meanWet = mean(PerFlyWetMass, na.rm = TRUE), .groups = "drop") %>%
  pivot_wider(names_from = Sex, values_from = meanWet, names_prefix = "WetMass_") %>%
  drop_na(WetMass_F, WetMass_M)


# Create the subtitle using paste0()
subtitle_text <- paste0("Male:Female wet mass phylogenetic-adjusted correlation r² = 0.86, HPD (0.52, 1.00), flies per pool ≥1")

# Plot
sex_corr_plot <- ggplot(df_plot, aes(x = WetMass_F, y = WetMass_M, color = SpeciesName)) +
  geom_point(size = 2, alpha = 0.5) +
  geom_smooth(method = "lm", se = FALSE, color = "black", linewidth = 0.8) +
  geom_text_repel(aes(label = SpeciesName), size = 2, max.overlaps = 10) +
  coord_cartesian() +
  theme_minimal(base_size = 12) +
  labs(
    x = "Female Per-fly wet mass (g)",
    y = "Male Per-fly wet mass (g)",
    title = "Wet mass of males vs. females across species",
    subtitle = subtitle_text,
    color = "Species"
  ) +
  theme(
    legend.position = "none",
    plot.subtitle = element_text(size = 8),
    panel.grid.minor = element_blank()
  )
sex_corr_plot

# Save it
ggsave("Figures/Fig2supp_PerFlyWetMass_SexCorr.png", sex_corr_plot,
       width = 6, height = 5, dpi = 300)
```
