## Supplementary material for "Life history and infection susceptibility parameters of *Drosophila* species reared on a common diet": File S2: FigC_Food_Comparisons.R

```
library(lme4)
library(lmerTest)   # gives p-values
library(emmeans)    # summarises by species

# analysis of different diets

df<-read_excel("Data_Raw/FoodComparisons_MAH_Bk3Pg40.xlsx") #
df <- df %>%
  mutate(EmergentsStart = as.Date(as.numeric(EmergentStart), origin = "1899-12-30"),
         EggToAdult = as.numeric(Egg2Adult))
df$PerFlyWM<-(df$TubeWeightWetMass-df$TubeWeightInitial)/df$n_in_Tube # in grams
str(df)

df$Diet <- recode(df$Diet,
                          "PENSTRAS" = "Pen. corn",
                          "CORN" = "Cam. corn",
                          "PROPIONIC" = "Prop recipe",
                          "UPLEM" = "EPFL corn")

# summarise dataset for plot later
summary_df <- df %>%
  filter(EggToAdult > 0, Diet != "INSTANT") %>%   # keep only valid measurements, remove INSTANT diet
  group_by(SpeciesName, Diet) %>%
  summarise(
        MeanEggToAdult = mean(EggToAdult, na.rm = TRUE),
    SDEggToAdult   = sd(EggToAdult, na.rm = TRUE),
    n_vials             = n_distinct(paste(SpeciesNumber, Diet, DateVialStart)),  # unique vials used to calc data
    n_m_emerg_perF      = mean(n_emergents[Sex == "M"]/Parent_F_N, na.rm = TRUE),
    n_f_emerg_perF      = mean(n_emergents[Sex == "F"]/Parent_F_N, na.rm = TRUE),
    n_emerg_perF        = ((n_emergents[Sex == "F"])+(n_emergents[Sex == "M"]))/Parent_F_N, na.rm = TRUE, # this number is outputting half of what it wants to
    PerFlyWM_mean       = mean(PerFlyWM, na.rm = TRUE),
    n_m_emerg_perF_SD      = sd(n_emergents[Sex == "M"]/Parent_F_N, na.rm = TRUE),
    n_f_emerg_perF_SD      = sd(n_emergents[Sex == "F"]/Parent_F_N, na.rm = TRUE),
    n_emerg_perF_SD        = sd((n_emergents[Sex == "F"])+(n_emergents[Sex == "M"])/Parent_F_N, na.rm = TRUE), #
    PerFlyWM_SD            = sd(PerFlyWM, na.rm = TRUE),
    Block = Block,
    DateVialStart = DateVialStart,
    Sex = Sex,
    .groups = "drop"
  ) %>%
  filter(n_vials > 0) %>%
  select(SpeciesName, n_vials, MeanEggToAdult, SDEggToAdult, Diet, n_m_emerg_perF, n_m_emerg_perF_SD, n_f_emerg_perF, n_f_emerg_perF_SD, n_emerg_perF, n_emerg_perF_SD, PerFlyWM_mean, PerFlyWM_SD,Block,DateVialStart,Sex)

summary_df$Diet <- recode(summary_df$Diet,
                  "PENSTRAS" = "Pen. corn",
                  "CORN" = "Cam. corn",
                  "PROPIONIC" = "Prop recipe",
                  "UPLEM" = "EPFL corn"
                  )

# summary_df$n_emerg_perF <- summary_df$n_m_emerg_perF + summary_df$n_f_emerg_perF
str(summary_df)
# print(summary_df, n=80)

write.csv(summary_df, "Data_clean/Rdata/FigC_line30_summary_df.csv", row.names = FALSE)

### Tree stuff

# plot tree with pruned species
all_tips <- tree$tip.label
species_with_data <- unique(summary_df$SpeciesName)
species_with_data <- gsub("\\s+", " ", species_with_data) # Replace multiple spaces with a single space

tips_to_drop <- setdiff(all_tips, species_with_data)
pruned_tree <- drop.tip(tree, tips_to_drop)

plot(pruned_tree, cex = 0.7, main = "")

### ggtree() part

# Ensure your data and tree are pruned and cleaned as you did
species_means <- summary_df %>% filter(!is.nan(MeanEggToAdult))

# Prune tree as you did
all_tips <- tree$tip.label
species_with_data <- species_means$SpeciesName
tips_to_drop <- setdiff(all_tips, species_with_data)
pruned_tree <- drop.tip(tree, tips_to_drop)

# Reorder species_means to match tree tip order
species_means <- species_means %>%
  mutate(SpeciesName = factor(SpeciesName, levels = pruned_tree$tip.label))

# Create tree plot with ggtree
is.waive <- function(x) inherits(x, "waiver") # updates ggtree to function with new ggplot2 version... is.waive() was changed to waiver()
p_tree <- ggtree(pruned_tree) + 
  geom_tiplab(size = 0, align = TRUE) + 
  theme_tree2() +
  theme(
    axis.text.x = element_blank(),
    axis.ticks.x = element_blank(),
    axis.title.x = element_blank(),
    axis.line.x = element_blank()   # all these element_blank() bits remove the genetic distance line and labels
    )
p_tree

### Egg-to-adult Bar plot

# Define order of species (top to bottom in plot)
species_order <- c("D. virilis", "D. immigrans", "D. testacea", "D. miranda", "D. suzukii", "D. melanogaster") # because of coord_flip, this has to be in reverse of the tree itself top to bottom

# Reorder factor levels accordingly
summary_df <- summary_df %>%
  mutate(SpeciesName = factor(SpeciesName, levels = species_order))
summary_df <- summary_df %>%
  filter(Diet != "INSTANT")

# Set the order manually
summary_df$Diet <- factor(summary_df$Diet, 
                          levels = c("Prop recipe", "EPFL corn", "Pen. corn", "Cam. corn")) #, "INSTANT"))

### fill in missing data as NA
# Create all combinations of SpeciesName and Diet
all_combos <- expand.grid(
  SpeciesName = levels(summary_df$SpeciesName),
  Diet = levels(summary_df$Diet)
)

# Join with your existing data
summary_df_filled <- all_combos %>%
  left_join(summary_df, by = c("SpeciesName", "Diet"))

# gg barplot
p_bar <- ggplot(summary_df_filled, aes(x = SpeciesName, # because of coord_flip(), species on the left, egg to adult on the bottom
                                       y = MeanEggToAdult, # because of coord_flip(), species on the left, egg to adult on the bottom
                                       fill = Diet), # fill based on diet
                                       group = Diet) + # group based on diet within species to produce multiple bars per species
  geom_col(position = position_dodge(width = 0.8), width = 0.6, alpha = 1, na.rm = FALSE) + # dodges the bars so they don't overlap
  geom_errorbar(aes(ymin = MeanEggToAdult - SDEggToAdult,
                    ymax = MeanEggToAdult + SDEggToAdult),
                width = 0.2,
                position = position_dodge(width = 0.8),
                na.rm = TRUE) +
  coord_flip() +
  scale_y_continuous(limits = c(0, 24), breaks = seq(0, 24, by = 4)) +
  scale_fill_manual(values = FigC_colours) +
  labs(x = "", y = "Egg-to-Adult (Mean days ± StDev)", fill = "Diet",
       title = "") + # Development time by species and diet (Mean ± StDev)
  theme_bw() +
  theme(strip.background = element_rect(fill = "grey90"),
        strip.text = element_text(face = "bold"),
        legend.position = "right",
        legend.text = element_text(size = 24),
        legend.title = element_text(size = 24),
        axis.title.x = element_text(size = 18),
        axis.text = element_text(face = "italic", size = 20)) +
  guides(fill = guide_legend(reverse = TRUE)) # reverse legend
p_bar

# Using Patchwork
p_combined <- p_tree + p_bar + plot_layout(widths = c(1, 3))
p_combined

# Save as PNG
ggsave(filename = "Figures/BuildingBlocks/FigC_egg2adult_barplot.png", plot = p_combined, width = 10, height = 8, dpi = 300)

###
# n_emergents
###
# Filter out invalid values if any

str(summary_df)

summary_df$Diet <- recode(summary_df$Diet,
                          "PENSTRAS" = "Pen. corn",
                          "CORN" = "Cam. corn",
                          "PROPIONIC" = "Prop recipe",
                          "UPLEM" = "EPFL corn"
                          )

summary_df$DateVialStart <- as.character(summary_df$DateVialStart)
df_plot <- summary_df %>%
  filter(!is.na(Diet), !is.na(SpeciesName), !is.na(n_emerg_perF),   Diet != "INSTANT") %>%
  group_by(SpeciesName, DateVialStart, Diet) %>%
  slice(which(!duplicated(n_emerg_perF))[1]) %>%  # first unique n_emerg_perF per group
  ungroup()

diet_order <- c("Prop recipe", "EPFL corn", "Pen. corn", "Cam. corn")
species_order <- c("D. melanogaster", "D. suzukii", "D. miranda", "D. testacea", "D. immigrans", "D. virilis")
df_plot <- df_plot %>%
  mutate(Diet = factor(Diet, levels = diet_order)) %>%
  mutate(SpeciesName = factor(SpeciesName, levels = species_order))

str(df_plot)
df_plot$Block<-as.character(df_plot$Block)

# --- Boxplot ---
p_box <- ggplot(df_plot, aes(x = Diet, y = n_emerg_perF, fill = Diet)) +
  geom_boxplot(alpha = 0.6, outlier.shape = NA, colour = "grey30") +   # semi-transparent boxes
  geom_jitter(aes(color = Diet, shape = Block),
              width = 0.3, height = 0, size = 1.5, alpha = 0.7, show.legend = FALSE) +  # jittered data points
  facet_wrap(~ SpeciesName, scales = "free_y") +   # one facet per species
  scale_fill_manual(values = FigC_colours) +
  scale_color_manual(values = FigC_colours) +
  labs(x = "",
       y = "Emergents per female",
       fill = "Diet",
       title = "") +
  theme_bw() +
  theme(
    strip.background = element_rect(fill = "grey90"),
    strip.text = element_text(face = "italic", size = 20), # facet individual titles
    axis.text.x = element_text(angle = 45, hjust = 1, size = 20),
    axis.text.y = element_text(size = 18),
    axis.title.y = element_text(size = 20),
    axis.title = element_text(size = 18),
    legend.position = "none"
  )
p_box

# --- Save to file ---
ggsave(filename = "Figures/z_not_used/FigC_n_emergentsperF_boxplot_facet.png",
       plot = p_box, width = 14, height = 8, dpi = 300)

#######
# Let's try that as a barplot
#######
str(df_plot)

# mixed model stats
df_emerg <- subset(df_plot)
model_emerg <- lmer(n_emerg_perF ~ Diet * SpeciesName + (1 | Block), data = df_emerg)
summary(model_emerg)
emmeans(model_emerg, pairwise ~ Diet | SpeciesName)

# Summarize per SpeciesName × Diet (mean ± SD)
df_bar1 <- df_plot %>%
  group_by(SpeciesName, Diet) %>%
  summarise(
    mean_emerg = mean(n_emerg_perF, na.rm = TRUE),
    sd_emerg   = sd(n_emerg_perF, na.rm = TRUE),
    .groups = "drop"
  )

# Barplot
p_bar1 <- ggplot(df_bar1, aes(x = Diet, y = mean_emerg, fill = Diet)) +
  geom_col(position = position_dodge(width = 0.8), width = 0.6, alpha = 0.6) +
  geom_errorbar(aes(ymin = mean_emerg, ymax = mean_emerg + sd_emerg),
                width = 0.2, position = position_dodge(width = 0.8)) +
  # overlay jittered raw points from df_plot
  geom_jitter(data = df_plot, aes(x = Diet, y = n_emerg_perF, color = Diet),
              width = 0.2, height = 0, size = 1.5, alpha = 0.7, show.legend = FALSE) +
  facet_wrap(~ SpeciesName, scales = "free_y") +
  scale_fill_manual(values = FigC_colours) +
  scale_color_manual(values = FigC_colours) +
  labs(x = "", y = "Emergents per female ± SD", fill = "Diet") +
  theme_bw() +
  theme(
    strip.background = element_rect(fill = "grey90"),
    strip.text = element_text(face = "italic", size = 20), # facet individual titles
    axis.text.x = element_text(angle = 45, hjust = 1, size = 20),
    axis.text.y = element_text(size = 18),
    axis.title.y = element_text(size = 20),
    axis.title = element_text(size = 18),
    legend.position = "none"
  )

p_bar1

# --- Save to file ---
ggsave(filename = "Figures/BuildingBlocks/FigC_n_emergentsperF_barplot_facet.png",
       plot = p_bar1, width = 10, height = 8, dpi = 300)


#######
# wet and dry mass
#######

str(df)
df$PerFlyWM<-df$PerFlyWM*1000 # convert g to mg

# will want to make a new df_plot because I removed the duplicate vials previously because it was n_emerg per Female rather than interested in male/female offspring themselves
df$Block <- as.factor(df$Block)
str(summary_df)
df_plot <- df %>%
  select(ParentVial,SpeciesName, Sex,Block,Diet,PerFlyWM)

diet_order <- c("Prop recipe", "EPFL corn", "Pen. corn", "Cam. corn")
species_order <- c("D. melanogaster", "D. suzukii", "D. miranda", "D. testacea", "D. immigrans", "D. virilis")
str(df_plot)
df_plot <- df_plot %>%
  filter(!is.na(Diet), !is.na(SpeciesName),  Diet != "INSTANT") %>%
  mutate(Diet = factor(Diet, levels = diet_order)) %>%
  mutate(SpeciesName = factor(SpeciesName, levels = species_order))
str(df_plot)

print(df_plot, n=50)

##########
# mixed effects model
##########

# Males only
df_M <- subset(df_plot, Sex == "M")
model_M <- lmer(PerFlyWM ~ Diet * SpeciesName + (1 | Block) + (1 | ParentVial), data = df_M)
summary(model_M)

# Females only
df_F <- subset(df_plot, Sex == "F")
model_F <- lmer(PerFlyWM ~ Diet * SpeciesName + (1 | Block) + (1 | ParentVial), data = df_F)
summary(model_F)

# get species-specific comparisons
emmeans(model_M, pairwise ~ Diet | SpeciesName)
emmeans(model_F, pairwise ~ Diet | SpeciesName)


# record output
sink("Figures/FigC_PerFlyWM_Diet_by_Species.txt")

cat("##### Models below use Prop recipe and D. melanogaster as the reference per their variable #####\n\n")
cat("===== MALES: Diet × SpeciesName model =====\n\n")
print(summary(model_M))
cat("\n\nANOVA (Type III):\n")
print(anova(model_M))

cat("\n\n\n===== FEMALES: Diet × SpeciesName model =====\n\n")
print(summary(model_F))
cat("\n\nANOVA (Type III):")
print(anova(model_F))

cat("\n\n\n\n##### emmeans gives within-species comparisons #####\n\n")
cat("\n\n===== emmeans =====\n\n\n")
cat("\n\n=== Males ===\n")
print(emmeans(model_M, pairwise ~ Diet | SpeciesName))
cat("\n\n=== Females ===\n")
print(emmeans(model_F, pairwise ~ Diet | SpeciesName))

sink()   # close sink


##########
##########

str(df_plot)

# Summarize per SpeciesName × Diet (mean ± SD)
df_bar1 <- df_plot %>%
  group_by(SpeciesName, Diet, Sex) %>%
  summarise(
    mean_PerFlyWM = mean(PerFlyWM, na.rm = TRUE),
    sd_PerFlyWM   = sd(PerFlyWM, na.rm = TRUE),
    .groups = "drop"
  )
df_bar1 <- df_bar1 %>%
  mutate(across(where(is.numeric),    ~ ifelse(is.infinite(.x) | is.nan(.x), 0, .x))) # handles missing values as 0

print(df_bar1,n=40)

### Males only
# Summarized data for males only
df_bar1_m <- df_bar1 %>%
  filter(Sex == "M")
# Corresponding raw data for jitter points
df_plot_m <- df_plot %>%
  filter(Sex == "M")

# Barplot
p_bar1_m <- ggplot(df_bar1_m, aes(x = Diet, y = mean_PerFlyWM, fill = Diet)) +
  geom_col(position = position_dodge(width = 0.8), width = 0.6, alpha = 0.6) +
  geom_errorbar(aes(ymin = mean_PerFlyWM, ymax = mean_PerFlyWM + sd_PerFlyWM),
                width = 0.2, position = position_dodge(width = 0.8)) +
  # overlay jittered raw points from df_plot
  geom_jitter(data = df_plot_m, aes(x = Diet, y = PerFlyWM, color = Diet),
              width = 0.2, height = 0, size = 1.5, alpha = 0.7, show.legend = FALSE) +
  facet_wrap(~ SpeciesName, scales = "fixed") +
  coord_cartesian(ylim = c(0, 0.0027*1000)) +
  scale_fill_manual(values = FigC_colours) +
  scale_color_manual(values = FigC_colours) +
  labs(x = "", y = "Per-male wet mass (mg) ± SD", fill = "Diet") +
  theme_bw() +
  theme(
    strip.background = element_rect(fill = "grey90"),
    strip.text = element_text(face = "italic", size = 20), # facet individual titles
    axis.text.x = element_text(angle = 45, hjust = 1, size = 20),
    axis.text.y = element_text(size = 18),
    axis.title.y = element_text(size = 20),
    axis.title = element_text(size = 18),
    legend.position = "none"
  )
p_bar1_m


# --- Save to file ---
ggsave(filename = "Figures/BuildingBlocks/FigC_mean_PerFlyWM_barplot_facet_M.png",
       plot = p_bar1_m, width = 10, height = 8, dpi = 300)


### Females only
# Summarized data for males only
df_bar1_f <- df_bar1 %>%
  filter(Sex == "F")
# Corresponding raw data for jitter points
df_plot_f <- df_plot %>%
  filter(Sex == "F")

# Barplot
p_bar1_f <- ggplot(df_bar1_f, aes(x = Diet, y = mean_PerFlyWM, fill = Diet)) +
  geom_col(position = position_dodge(width = 0.8), width = 0.6, alpha = 0.6) +
  geom_errorbar(aes(ymin = mean_PerFlyWM, ymax = mean_PerFlyWM + sd_PerFlyWM),
                width = 0.2, position = position_dodge(width = 0.8)) +
  # overlay jittered raw points from df_plot
  geom_jitter(data = df_plot_f, aes(x = Diet, y = PerFlyWM, color = Diet),
              width = 0.2, height = 0, size = 1.5, alpha = 0.7, show.legend = FALSE) +
  facet_wrap(~ SpeciesName, scales = "fixed") +
  coord_cartesian(ylim = c(0, 0.0042*1000)) +
  scale_fill_manual(values = FigC_colours) +
  scale_color_manual(values = FigC_colours) +
  labs(x = "", y = "Per-female wet mass (mg) ± SD", fill = "Diet") +
  theme_bw() +
  theme(
    strip.background = element_rect(fill = "grey90"),
    strip.text = element_text(face = "italic", size = 20), # facet individual titles
    axis.text.x = element_text(angle = 45, hjust = 1, size = 20),
    axis.text.y = element_text(size = 18),
    axis.title.y = element_text(size = 20),
    axis.title = element_text(size = 18),
    legend.position = "none"
  )
p_bar1_f


# --- Save to file ---
ggsave(filename = "Figures/BuildingBlocks/FigC_mean_PerFlyWM_barplot_facet_F.png",
       plot = p_bar1_f, width = 10, height = 8, dpi = 300)
```
