## Supplementary material for "Life history and infection susceptibility parameters of *Drosophila* species reared on a common diet": File S2: FigC_Food_LipidFloating.R

```
df_lipid<-read_excel("Data_Raw/FoodComparisons_LipidFloating.xlsx")
df_lipid$ratio<- df_lipid$floater/df_lipid$`total larvae`
str(df_lipid)
unique(df_lipid$species)

df_lipid <- df_lipid %>%
  mutate(
    diet = case_when(
      diet == "prop" ~ "Prop recipe",
      diet == "uplem" ~ "EPFL corn",
      diet == "penstra" ~ "Pen. corn",
      diet == "cambridge" ~ "Cam. corn",
      TRUE ~ diet),
    diet = factor(diet, levels = diet_order),
    block = factor(block)
    )

diet_order <- c("Prop recipe", "EPFL corn", "Pen. corn", "Cam. corn")
species_order <- c("D. melanogaster", "D. suzukii", "D. miranda", "D. testacea", "D. immigrans", "D. virilis")
df_plot <- df_lipid %>%
  mutate(diet = factor(diet, levels = diet_order)) %>%
  mutate(species = factor(species, levels = species_order))

# --- Boxplot ---
p_box <- ggplot(df_plot, aes(x = diet, y = ratio, fill = diet)) +
  geom_boxplot(alpha = 0.6, outlier.shape = NA, colour = "grey30") +   # semi-transparent boxes
  geom_jitter(aes(color = diet, shape = block),
              width = 0.3, height = 0, size = 1.5, alpha = 0.7, show.legend = FALSE) +  # jittered data points
  scale_y_continuous(limits = c(0, 1), breaks = seq(0, 1, by = .25)) +
  facet_wrap(~ species, scales = "free_y") +   # one facet per species
  scale_fill_manual(values = FigC_colours) +
  scale_color_manual(values = FigC_colours) +
  labs(x = "",
       y = "Proportion larvae floating",
       fill = "diet",
       title = "") +
  theme_bw() +
  theme(
    strip.background = element_rect(fill = "grey90"),
    strip.text = element_text(face = "italic", size = 20), # facet individual titles
    axis.text.x = element_text(angle = 45, hjust = 1, size = 20),
    axis.text.y = element_text(size = 18),
    axis.title.y = element_text(size = 20),
    axis.title = element_text(size = 18),
    legend.position = "none"
  )
p_box

# --- Save to file ---
ggsave(filename = "Figures/BuildingBlocks/FigC_FloatingRatio_boxplot_facet.png",
       plot = p_box, width = 14, height = 8, dpi = 300)
```
