## Supplementary material for "Life history and infection susceptibility parameters of *Drosophila* species reared on a common diet": File S2: FigC_Magick_together2.R

```
##############
# magick together
##############
library(magick)

# Read the images
e2a_img <- image_read("Figures/BuildingBlocks/FigC_egg2adult_barplot.png")
n_e_img  <- image_read("Figures/BuildingBlocks/FigC_n_emergentsperF_barplot_facet.png")
mwm_img <- image_read("Figures/BuildingBlocks/FigC_mean_PerFlyWM_barplot_facet_M.png")
fwm_img <- image_read("Figures/BuildingBlocks/FigC_mean_PerFlyWM_barplot_facet_F.png")
lipid_img <- image_read("Figures/BuildingBlocks/FigC_FloatingRatio_boxplot_facet.png")
yeast_img <- image_read("Figures/BuildingBlocks/FigC_yeast_added.png")
float_img <- image_read("Figures/BuildingBlocks/FloatingLarvae.png")


# Create a transparent spacer (adjust height as needed)
spacer2_width <- 100  # pixels
spacer2 <- image_blank(width = spacer2_width,
                       height = image_info(n_e_img)$height,
                       color = "white") # "none" = transparent

top_row <- c(e2a_img,spacer2,n_e_img) %>%
  image_append(stack = FALSE)  # FALSE = horizontal

# CHANGED BOT AND MID ROWS
bot_row <- c(mwm_img,spacer2,fwm_img) %>%
  image_append(stack = FALSE)  # FALSE = horizontal

# CHANGED BOT AND MID ROWS
mid_row <- c(float_img,lipid_img,spacer2,yeast_img) %>%
  image_append(stack = FALSE)  # FALSE = horizontal

# Scale each image to the same height
scaled_imgs <- lapply(
  list(float_img, lipid_img, spacer2, yeast_img),
  function(img) image_scale(img, "x2100"))
mid_row <- image_append(do.call(c, scaled_imgs), stack = FALSE)

# Create a transparent spacer (adjust height as needed)
spacer_height <- 100  # pixels
spacer <- image_blank(width = image_info(top_row)$width,
                      height = spacer_height,
                      color = "white") # "none" = transparent

# Combine top row with bottom row (facet)
final_plot <- image_append(c(top_row, spacer, spacer,mid_row,spacer,bot_row), stack = TRUE)

# Save final combined image
image_write(final_plot, path = "Figures/BuildingBlocks/Combined_FigureC.png", format = "png")


### add letters
# Load the image
library(png)
img <- readPNG("Figures/BuildingBlocks/Combined_FigureC.png")
img_dim <- dim(img)

# Open PNG device (same size as image)
png("Figures/Fig3_FoodComparisons2.png", width = img_dim[2], height = img_dim[1], bg = "transparent") # bg = "transparent" prevents the hairline border being drawn alongside the par expression below
par(mar = c(0, 0, 0, 0), xaxs = "i", yaxs = "i") # Remove margins and set exact axis limits

# Plot blank canvas
plot(1, type = "n", xlim = c(0, img_dim[2]), ylim = c(0, img_dim[1]),
     xaxt = 'n', yaxt = 'n', xlab = '', ylab = '', asp = 1)

# Draw the image
rasterImage(img, 0, 0, img_dim[2], img_dim[1])

# Add labels
text(x = 50, y = 7130, labels = "A", cex = 12, font = 2)
text(x = 3150, y = 7130, labels = "B", cex = 12, font = 2)
text(x = 50, y = 4700, labels = "C", cex = 12, font = 2)
text(x = 4420, y = 4700, labels = "E", cex = 12, font = 2)
text(x = 50, y = 2380, labels = "D", cex = 12, font = 2)
# text(x = 3150, y = 2380, labels = "D''", cex = 12, font = 2)
text(x = 360, y = 2060, labels = "♂", cex = 24, font = 2)
text(x = 3440, y = 2060, labels = "♀", cex = 24, font = 2)

# Close device to save
dev.off()
```
