## Supplementary material for "Life history and infection susceptibility parameters of *Drosophila* species reared on a common diet": File S2: FigC_optional_stacked_plot.R

```
###
# TESTING A STACKED BARPLOT
###

# Step 1: pivot longer for stacking, keeping Diet as a variable
summary_long <- summary_df_filled %>%
  pivot_longer(
    cols = c(n_m_emerg_perF, n_f_emerg_perF),
    names_to = "Sex",
    values_to = "n_emerg_perF_stack",
    names_prefix = "n_",
    names_transform = list(Sex = ~ifelse(.x == "m_emerg_perF", "Male", "Female"))
  )

# Step 2: Plot
p_stacked_diet <- ggplot(summary_long, 
                         aes(x = Diet,
                             y = n_emerg_perF_stack,
                             fill = Diet,
                             pattern = Sex)) +
  geom_col_pattern(
    position = "stack",
    width = 0.6,
    pattern_fill = "white",      # background of hatch
    pattern_colour = "white",    # color of hatch lines
    pattern_density = 0.03,
    pattern_spacing = 0.025,
    pattern_angle = 90           # first set of lines
  ) +
  # Error bars from total per female
  geom_errorbar(data = summary_long,
                aes(x = Diet,
                    ymin = n_emerg_perF,
                    ymax = n_emerg_perF + n_emerg_perF_SD),
                width = 0.0,
                inherit.aes = FALSE) +
  facet_wrap(~ SpeciesName, ncol = 3, scales = "free_x") +
  coord_flip() +
  scale_fill_manual(values = FigC_colours) +
  labs(x = "", 
       y = "Total emergents per female", 
       fill = "Diet",
       title = "Total emergents per female by species and diet (Mean ± StDev)") +
  theme_bw() +
  theme(strip.background = element_rect(fill = "grey90"),
        strip.text = element_text(face = "bold.italic", size = 12))
# p_stacked_diet

# Save as PNG
ggsave(filename = "Figures/z_not_used/FigC_n_emergentsperF_barplot_wrap_stacked.png", plot = p_stacked_diet, width = 9, height = 8, dpi = 300)
```
