## Supplementary material for "Life history and infection susceptibility parameters of *Drosophila* species reared on a common diet": File S2: FigC_WickAdd_barplot.R

```
library(ggpattern)
# analysis of different diets

df<-read_excel("Data_Raw/WickAdded_MAH_Bk3Pg57.xlsx") #
df <- df %>%
  mutate(EmergentsStart = as.Date(as.numeric(EmergentStart), origin = "1899-12-30"),
         EggToAdult = as.numeric(Egg2Adult))
df$PerFlyWM<-(df$TubeWeightWetMass-df$TubeWeightInitial)/df$n_in_Tube # in grams
str(df)


n_Females<-sum(df$n_emergents[df$Sex=="F"], na.rm=T)
n_Females
n_Males<-sum(df$n_emergents[df$Sex=="M"], na.rm=T)
n_Males
DmirRatioMF<-n_Females/n_Males
DmirRatioMF
n_Males/(n_Males+n_Females)
n_Females/(n_Males+n_Females)
sum(df$n_emergents,na.rm=T)

df$Diet <- recode(df$Diet,
                  "PROP_WICK" = "Prop + wick",
                  "PROPIONIC" = "Prop recipe"
)

# summarise dataset for plot later
summary_df <- df %>%
  group_by(SpeciesName, Diet, Block,) %>%
  summarise(
    # MeanEggToAdult = mean(EggToAdult, na.rm = TRUE),
    # SDEggToAdult   = sd(EggToAdult, na.rm = TRUE),
    n_m_emerg_perF      = mean(n_emergents[Sex == "M"]/Parent_F_N, na.rm = TRUE),
    n_f_emerg_perF      = mean(n_emergents[Sex == "F"]/Parent_F_N, na.rm = TRUE),
    n_emerg_perF        = mean((n_emergents[Sex == "F"])+(n_emergents[Sex == "M"]))/Parent_F_N, na.rm = TRUE, # this number is outputting half of what it wants to
    # PerFlyWM_mean       = mean(PerFlyWM, na.rm = TRUE),
    n_m_emerg_perF_SD      = sd(n_emergents[Sex == "M"]/Parent_F_N, na.rm = TRUE),
    n_f_emerg_perF_SD      = sd(n_emergents[Sex == "F"]/Parent_F_N, na.rm = TRUE),
    n_emerg_perF_SD        = sd((n_emergents[Sex == "F"])+(n_emergents[Sex == "M"])/Parent_F_N, na.rm = TRUE), #
    # PerFlyWM_SD            = sd(PerFlyWM, na.rm = TRUE),
    .groups = "drop"
  ) %>%
  select(SpeciesName, Block,Diet, n_m_emerg_perF, n_m_emerg_perF_SD, n_f_emerg_perF, n_f_emerg_perF_SD, n_emerg_perF, n_emerg_perF_SD)

# summary_df$n_emerg_perF <- summary_df$n_m_emerg_perF + summary_df$n_f_emerg_perF
str(summary_df)
print(summary_df, n = 20)

write.csv(summary_df, "Data_clean/Rdata/FigC_wick_df.csv", row.names = FALSE)


# Filter out invalid values if any

str(summary_df)

summary_df$Block <- as.character(summary_df$Block)
df_plot <- summary_df %>%
  filter(!is.na(Diet), !is.na(SpeciesName), !is.na(n_emerg_perF)) %>%
  group_by(SpeciesName, Block, Diet) %>%
  slice(which(!duplicated(n_emerg_perF))[1]) %>%  # first unique n_emerg_perF per group
  ungroup()

str(df_plot)

unique(df_plot$Diet)

diet_order <- c("Prop recipe", "Prop + wick")
species_order <- c("D. miranda")
df_plot <- df_plot %>%
  mutate(Diet = factor(Diet, levels = diet_order)) %>%
  mutate(SpeciesName = factor(SpeciesName, levels = species_order))


#######
# Let's try that as a barplot
#######


# Summarize per SpeciesName × Diet (mean ± SD)
df_bar1 <- df_plot %>%
  group_by(SpeciesName, Diet) %>%
  summarise(
    mean_emerg = mean(n_emerg_perF, na.rm = TRUE),
    sd_emerg   = sd(n_emerg_perF, na.rm = TRUE),
    .groups = "drop"
  )

# Barplot
p_bar1 <- ggplot(df_bar1, aes(x = Diet, y = mean_emerg, fill = Diet, pattern = ifelse(as.numeric(Diet) == 2, "stripe", "none"))) + # Diet = 2 because PROP_YEAST is alphabetically first
  #geom_col(position = position_dodge(width = 0.8), width = 0.6, alpha = 0.6) + # old code if I don't want to do a pattern fill
  geom_col_pattern(
    position = position_dodge(width = 0.8), width = 0.6, alpha = 0.6, pattern_angle = 45, pattern_density = 0.3, pattern_spacing = 0.1,
    pattern_alpha = .1, pattern_fill = "white") +
  geom_errorbar(aes(ymin = mean_emerg, ymax = mean_emerg + sd_emerg),
                width = 0.2, position = position_dodge(width = 0.8)) +
  # overlay jittered raw points from df_plot
  geom_jitter(data = df_plot, aes(x = Diet, y = n_emerg_perF, color = Diet),
              width = 0.2, height = 0, size = 1.5, alpha = 0.8, show.legend = FALSE) +
  facet_wrap(~ SpeciesName, scales = "free_y") +
  scale_fill_manual(values = FigC_colours) +
  scale_color_manual(values = FigC_colours) +
  scale_pattern_manual(values = c("stripe" = "stripe", "none" = "none")) +
  labs(x = "", y = "Emergents per female ± SD", fill = "Diet") +
  theme_bw() +
  theme(
    strip.background = element_rect(fill = "grey90"),
    strip.text = element_text(face = "italic", size = 11),
    axis.text.x = element_text(angle = 45, hjust = 1, size = 14),
    axis.title.y = element_text(size = 14),
    axis.title = element_text(size = 12),
    legend.position = "none"
  )
p_bar1

# --- Save to file ---
ggsave(filename = "Figures/z_not_used/FigC_wick_added.png",
       plot = p_bar1, width = 6, height = 8, dpi = 300)


# boxplot version
p_box1 <- ggplot(df_plot, aes(x = Diet, y = n_emerg_perF, fill = Diet,
                              pattern = ifelse(as.numeric(Diet) == 2, "stripe", "none"))) +
  geom_boxplot_pattern(
    alpha = 0.6,
    pattern_angle = 45,
    pattern_density = 0.3,
    pattern_spacing = 0.1,
    pattern_alpha = 0.1,
    pattern_fill = "white",
    outlier.shape = NA  # hide default outliers since you add jitter
  ) +
  scale_y_continuous(limits = c(0, 2.5), breaks = seq(0, 2.5, by = .5)) + # adjust later
  geom_jitter(aes(color = Diet),
              width = 0.2, height = 0,
              size = 2, alpha = 0.8, show.legend = FALSE) +
  facet_wrap(~ SpeciesName, scales = "free_y") +
  scale_fill_manual(values = FigC_colours) +
  scale_color_manual(values = FigC_colours) +
  scale_pattern_manual(values = c("stripe" = "stripe", "none" = "none")) +
  labs(x = "", y = "Emergents per female ± SD", fill = "Diet") +
  theme_bw() +
  theme(
    strip.background = element_rect(fill = "grey90"),
    strip.text = element_text(face = "italic", size = 11),
    axis.text.x = element_text(angle = 45, hjust = 1, size = 14),
    axis.title.y = element_text(size = 14),
    axis.title = element_text(size = 12),
    legend.position = "none"
  )
p_box1

str(df_plot)
t.test(df_plot$n_emerg_perF~df_plot$Diet)

# --- Save to file ---
ggsave(filename = "Figures/Fig3supp2_wick_added.png",
       plot = p_box1, width = 6, height = 8, dpi = 300)
```
