## Supplementary material for "Life history and infection susceptibility parameters of *Drosophila* species reared on a common diet": File S2: FigCsupp1_Foods_E2A_analysis.R

```
library(tidyr)
library(ggplot2)
library(dplyr)
library(readxl)
library(RColorBrewer)
library(forcats)
library(ggpubr)
library(patchwork)

EA_df3 <- read.csv("Data_Clean/E2A_CornPropPENSTRAS.csv") 

EA_df3$Food <- recode(EA_df3$Food,
                      "PENSTRAS" = "Pen. corn",
                      "CORN" = "Cam. corn",
                      "PROPIONIC" = "Prop recipe",
                      # "UPLEM" = "EPFL corn"
                      )

# Identify species with all three food types present
species_all_foods <- EA_df3 %>%
  group_by(SpeciesName) %>%
  summarise(food_count = n_distinct(Food)) %>%
  filter(food_count == 3) %>%
  pull(SpeciesName)

# Filter original dataframe to keep only those species
EA_df <- EA_df3 %>%
  filter(SpeciesName %in% species_all_foods)

####
# BOXPLOTS FOR A VISUAL OF THE VARIANCE BY SPECIES
####
species_list <- split(EA_df, EA_df$SpeciesName)

# Open PNG device; adjust width and height in pixels as needed
png(filename = "Figures/BuildingBlocks/Egg2Adult_by_Food_and_Species.png", width = 3500, height = 3000, res = 150)

par(mfrow = c(5, 4)) # 5 rows, 4 cols

food_colors <- c("grey90", "grey65", "grey40")


unique_blocks <- sort(unique(EA_df$Block))
n_blocks <- length(unique_blocks)

# Always generate palette with brewer.pal, capped at 12 colors (Set3 max)
n_colors <- max(3, min(n_blocks, 12))
block_colors <- brewer.pal(n = n_colors, name = "Set3")

# If you have more blocks than 12, repeat colors (or you can extend with other palettes)
if (n_blocks > n_colors) {
  block_colors <- rep(block_colors, length.out = n_blocks)
}

block_color_map <- setNames(block_colors, unique_blocks)


for (i in seq_along(species_list)) {
  species_data <- species_list[[i]]
  
  boxplot(Egg2Adult ~ Food, data = species_data, main = names(species_list)[i],
          col = food_colors, cex.main = 3, cex.axis = 2,
          ylim = c(10,25), yaxp = c(10,25,5), xlab = "")
  
  x_jitter <- jitter(as.numeric(factor(species_data$Food, levels = c("Pen. corn", "Cam. corn", "Prop recipe"))))
  
  point_cols <- block_color_map[species_data$Block]
  
  points(x_jitter, species_data$Egg2Adult, col = point_cols, cex = 1.5, pch = 16)
}

#legend("topright", legend = unique_blocks, col = block_color_map, pch = 16, cex = 1.5, bty = "n")

dev.off()  # close the PNG device


str(EA_df)

####
# LINE PLOT
####

# Step 1: Calculate mean and SD Egg2Adult per Species and Food
means_df <- EA_df %>%
  group_by(SpeciesName, Food) %>%
  summarise(mean_Egg2Adult = mean(Egg2Adult),
            sd_Egg2Adult = sd(Egg2Adult),
            .groups = "drop")

# Step 2: Rank species by overall mean
species_order <- means_df %>%
  group_by(SpeciesName) %>%
  summarise(overall_mean = mean(mean_Egg2Adult), .groups = "drop") %>%
  arrange(overall_mean) %>%
  mutate(SpeciesName = factor(SpeciesName, levels = SpeciesName))  # sets order

# Step 3: Join ranking back into data
means_ranked <- means_df %>%
  left_join(species_order %>% select(SpeciesName), by = "SpeciesName") %>%
  mutate(SpeciesName = factor(SpeciesName, levels = levels(species_order$SpeciesName)))

# Step 4: Custom food colors
food_colors <- c("Pen. corn" = "#0087d0", "Cam. corn" = "#ffba77", "Prop recipe" = "#ac006d")

# Step 5: Plot
p <- ggplot() +
  # geom_line(data = means_ranked %>% filter(!is.na(mean_Egg2Adult)),
  #           aes(x = SpeciesName, y = mean_Egg2Adult, group = Food, color = Food),
  #           linewidth = 1.2) +
  geom_point(data = means_ranked %>% filter(!is.na(mean_Egg2Adult)),
             aes(x = SpeciesName, y = mean_Egg2Adult, color = Food),
             size = 3, alpha = 0.6, position = position_dodge(width = .25)) +
  geom_errorbar(data = means_ranked %>% filter(!is.na(mean_Egg2Adult) & !is.na(sd_Egg2Adult)),
                aes(x = SpeciesName,
                    ymin = mean_Egg2Adult - sd_Egg2Adult,
                    ymax = mean_Egg2Adult + sd_Egg2Adult,
                    color = Food),
                width = 0.3, linewidth = 0.8, alpha = 0.3, position = position_dodge(width = .25)) +
  scale_color_manual(values = food_colors) +
  scale_y_continuous(limits = c(8, 23), breaks = seq(8, 23, by = 3)) +
  labs(title = "", # Mean Egg-to-Adult by species (error bars = StDev)
       x = "",
       y = "Egg-to-Adult mean (days)") +
  theme_classic(base_size = 14) +
  theme(axis.text.x = element_text(angle = 45, hjust = 1, size = 12),
        axis.title.y = element_text(size = 14),
        legend.title = element_blank(),
        legend.text = element_text(size = 18),
        plot.title = element_text(hjust = 0.5))
p

# Save to file
ggsave("Figures/BuildingBlocks/Egg2Adult_means_by_SpeciesRank_and_Food_with_SD_ggplot.png",
       plot = p, width = 12, height = 8, dpi = 300)


###
# lm()
###

# Fit linear model
lm_fit <- lm(mean_Egg2Adult ~ Food, data = means_df)
print(summary(lm_fit)) # yup, significant diff between PROP and CORN (P = .012)

str(means_df)
# print(means_df, n = 100)

# Calculate summary stats (mean and sd) for each Food
summary_stats <- means_df %>%
  group_by(Food) %>%
  summarise(
    mean_val = mean(mean_Egg2Adult),
    sd_val = sd(mean_Egg2Adult)
  )

# Run one-way ANOVA
anova_res <- aov(mean_Egg2Adult ~ Food, data = means_df)
summary(anova_res)
# Run Tukey's HSD post-hoc test
tukey_res <- TukeyHSD(anova_res)
print(tukey_res) # p = .031 for PROP-CORN, .257 for PENSTRAS-PROP

# define comparisons to plot based on the ANOVA tukeyHSD results
my_comparisons <- list(c("Cam. corn", "Prop recipe"))

# Plot
p3<-ggplot(means_df, aes(x = Food, y = mean_Egg2Adult, color = SpeciesName)) +
  geom_jitter(width = 0.2, height = 0, size = 2, alpha = 0.7) +
  geom_point(data = summary_stats, aes(x = Food, y = mean_val), color = "black", size = 4, inherit.aes = FALSE) +
  geom_errorbar(data = summary_stats, aes(x = Food, ymin = mean_val - sd_val, ymax = mean_val + sd_val),
                width = 0.1, color = "black", size = 1, inherit.aes = FALSE) +
  theme_classic() +
  theme(legend.position = "none") +
  labs(y = "Egg-to-Adult species means (days)", x = "",
       title = "",
       color = "Species") +
  theme(axis.text.x = element_text(angle = 45, hjust = 1, size = 14),
        axis.title.y = element_text(size = 14))
p3 <- p3 + coord_cartesian(ylim = c(12, 22))
p3 <- p3 + scale_y_continuous(breaks = seq(12, 22, by = 1))
p3 <- p3 + 
  stat_compare_means(comparisons = my_comparisons, method = "t.test", 
                     label = "p.signif", tip.length = 0.01, size = 6, fontface = "bold", lwd = 4)
p3

ggsave("Figures/BuildingBlocks/Egg2Adult_mean_SD_vertical_lines.png", plot = p3, width = 10, height = 6, dpi = 300)


###
# COMBO PLOT
###
# Arrange plots with labels A and B
combined_plot <- p + p3 + 
  plot_annotation(tag_levels = 'A')

# Save or print
print(combined_plot)
ggsave("Figures/Fig3supp1_CORN-PENSTRAS-PROP.png", plot = combined_plot, width = 16, height = 6, dpi = 300)


# # plot df
# summary_df <- EA_df %>%
#   filter(Food %in% c("Pen. corn", "Cam. corn")) %>%
#   group_by(Species, Food) %>%
#   summarize(Mean_Egg2Adult = mean(Egg2Adult, na.rm = TRUE),
#             SD_Egg2Adult = sd(Egg2Adult, na.rm = TRUE))
# 
# # Ungroup the summary_df
# summary_df <- ungroup(summary_df)
# 
# # Calculate the difference between mean Egg2Adult for CORN and PENSTRAS within each species
# difference_df <- summary_df %>%
#   filter(Food %in% c("Pen. corn", "Cam. corn")) %>%
#   group_by(Species) %>%
#   summarize(Difference = mean(Mean_Egg2Adult[Food == "Cam. corn"], na.rm = TRUE) - mean(Mean_Egg2Adult[Food == "Pen. corn"], na.rm = TRUE))
# 
# quartz(h=8,w=7)
# # Plot the differences
# ggplot(difference_df, aes(x = Species, y = Difference)) +
#   geom_bar(stat = "identity", fill = "skyblue", color = "black") +
#   geom_text(aes(label = round(Difference, 2)), vjust = -0.5, size = 2) +
#   labs(title = "",
#        x = "",
#        y = "avgCORN - avgPENSTRAS") +
#   theme(axis.text.x = element_text(angle = 90, vjust = 0.5, hjust=1)) +
#   ylim(-2,2)  # Rotate x-axis labels vertically
# 
```
