## Supplementary material for "Life history and infection susceptibility parameters of *Drosophila* species reared on a common diet": File S2: FigD_Magick_together.R

```
##############
# magick together
##############
library(magick)

# Read the images
corr_img <- image_read("Figures/BuildingBlocks/FigD_Correlations.png")
gfp_img  <- image_read("Figures/BuildingBlocks/FigD_GFPsignal.png")
facet_img <- image_read("Figures/BuildingBlocks/FigD_cornmeal_facet_surv.png")
top1 <- image_read("Figures/BuildingBlocks/FigD_CFUs_Corn_vs_Prop_MRS.png")
top2 <- image_read("Figures/BuildingBlocks/FigD_CFUs_Corn_vs_Prop_YPG.png")
top3 <- image_read("Figures/BuildingBlocks/FigD_CFUs_Corn_vs_Prop_LB.png")
top4 <- image_read("Figures/BuildingBlocks/FigD_CFUs_Corn_vs_Prop_BHI.png")

top_row <- c(top1,top2,top3,top4) %>%
  image_append(stack = FALSE)  # FALSE = horizontal

# Create a transparent spacer (adjust height as needed)
spacer_height <- 80  # pixels
spacer <- image_blank(width = image_info(top_row)$width, 
                      height = spacer_height, 
                      color = "white")  # "none" = transparent

# Combine top row: corr (left) + GFP (right)
mid_row <- image_append(c(corr_img, gfp_img))

# Combine top row with bottom row (facet)
final_plot <- image_append(c(top_row, spacer, mid_row, facet_img), stack = TRUE)

# Save final combined image
image_write(final_plot, path = "Figures/BuildingBlocks/Combined_FigureD.png", format = "png")


### add letters
# Load the image
library(png)
img <- readPNG("Figures/BuildingBlocks/Combined_FigureD.png")
img_dim <- dim(img)

# Open PNG device (same size as image)
png("Figures/Fig4_MillieData.png", width = img_dim[2], height = img_dim[1], bg = "transparent") # bg = "transparent" prevents the hairline border being drawn alongside the par expression below
par(mar = c(0, 0, 0, 0), xaxs = "i", yaxs = "i") # Remove margins and set exact axis limits

# Plot blank canvas
plot(1, type = "n", xlim = c(0, img_dim[2]), ylim = c(0, img_dim[1]),
     xaxt = 'n', yaxt = 'n', xlab = '', ylab = '', asp = 1)

# Draw the image
rasterImage(img, 0, 0, img_dim[2], img_dim[1])

# Add labels
text(x = 650, y = 3300, labels = "A", cex = 8, font = 2)
text(x = 1480, y = 3300, labels = "B", cex = 8, font = 2)
text(x = 2330, y = 3300, labels = "C", cex = 8, font = 2)
text(x = 3160, y = 3300, labels = "D", cex = 8, font = 2)
text(x = 50, y = 2500, labels = "E", cex = 8, font = 2)
text(x = 2700, y = 2500, labels = "F", cex = 8, font = 2)
text(x = 50, y = 1100, labels = "G", cex = 8, font = 2)

# Close device to save
dev.off()
```
