## Supplementary material for "Life history and infection susceptibility parameters of *Drosophila* species reared on a common diet": File S2: FigD_Mark_CFU_data.R

```
# load libraries
library(readxl)
library(readr)
library(ape)
library(ggtree)
library(ggplot2)
library(dplyr)
library(patchwork)
library(survminer)
library(survival)
library(phytools)
library(stargazer)
library(tidyverse)
library(lme4)
library(lmerTest)
library(ggpubr)


########
# CFUs
########

df1<-read_excel("Data_Raw/VialMicrobiome_CFUs.xlsx", sheet = 1)
df1_sub<-df1 %>%
  select(ID.Vial,Species,Diet,Collected.Date, NormalizedCFUs, Log10normCFUs,PlateType)
df2<-read_excel("Data_Raw/VialMicrobiome_CFUs.xlsx", sheet = 2)
df2_sub<-df2 %>%
  select(ID.Vial,Species,Diet,Collected.Date, NormalizedCFUs, Log10normCFUs,PlateType)

df<-rbind(df1_sub,df2_sub)
df$PlateType <- factor(df$PlateType, levels = c("MRS","BHI","LB","YPG"))

str(df)
unique(df$Diet)
df$Species <- factor(df$Species, levels = c("D. melanogaster", "D. virilis", "D. immigrans")) # reorder

df <- df %>%
  filter(Diet %in% c("Corn", "Prop"))

df$Collected.Date<-as.factor(df$Collected.Date)
df

PilotData<-lmer(df$Log10normCFUs~df$Diet + (1|df$Collected.Date) + df$Species + df$PlateType) #lme4 package wouldn't install
summary(PilotData)


# Make sure Diet and Species are factors
df$Diet <- factor(df$Diet, levels = c("Corn", "Prop"))
df$Species <- factor(df$Species)

# Run one-way ANOVA
anova_res <- aov(Log10normCFUs ~ Diet+Species+Collected.Date + df$PlateType, data = df)
summary(anova_res)
# Run Tukey's HSD post-hoc test
tukey_res <- TukeyHSD(anova_res)
print(tukey_res) # p < .001 for PROP vs CORN

df

# # Make sure Diet is a factor for proper grouping, level order
# df$Diet <- factor(df$Diet, levels = c("Corn", "Prop"))

# just subsetting the df cleanly later for the plots
df_MRS <- df %>% 
  filter(PlateType == "MRS")
df_LB <- df %>% 
  filter(PlateType == "LB")
df_BHI <- df %>% 
  filter(PlateType == "BHI")
df_YPG <- df %>% 
  filter(PlateType == "YPG")


##########
# MRS
##########

# Calculate summary stats for error bars (mean ± SD)
summary_df <- df_MRS %>%
  group_by(Diet) %>%
  summarise(
    mean = mean(Log10normCFUs, na.rm = TRUE),
    sd = sd(Log10normCFUs, na.rm = TRUE),
    .groups = 'drop'
  )

# Plot
p<-ggplot(df_MRS, aes(x = Diet, y = Log10normCFUs, fill = Species)) +
  geom_jitter(aes(shape = Species, fill = Species),
              position = position_jitterdodge(jitter.width = 0.3, dodge.width = 0.8),
              size = 2,
              alpha = 0.7,
              color = "black",   # black border to avoid missing warning
              stroke = 0) +      # but no visible border thickness
  scale_fill_manual(values = c(
    "D. melanogaster" = "#1b9e77",
    "D. virilis" = "#d95f02",
    "D. immigrans" = "#7570b3")) +
  scale_shape_manual(values = c(21, 22, 24)) +  # filled shapes that accept fill color
  geom_errorbar(data = summary_df,
                aes(x = Diet, ymin = mean - sd, ymax = mean + sd),
                width = 0.2,
                size = 0.5,
                color = "black",
                inherit.aes = FALSE) +
  geom_segment(data = summary_df,
               aes(x = as.numeric(Diet) - 0.3, xend = as.numeric(Diet) + 0.3,
                   y = mean, yend = mean),
               color = "black",
               size = 1,
               inherit.aes = FALSE) +
  theme_classic() +
  labs(x = NULL, y = "MRS CFUs (Log10)", fill = "Species") +
  theme(axis.title.y = element_text(size = 12),
        legend.position = "left",
        legend.text = element_text(face = "italic", size = 16),
        legend.title = element_text(size = 16),
        axis.text.x = element_text(size = 12)) +
  scale_y_continuous(limits = c(0,11), breaks = seq(0, 10, by = 2))
# the actual statistical test is above in the Tukey's HSD and lmer(), but t.test here allows the plot to draw an error bar, and the asterisk count is the same anyways.
# Very lazy coding here, but the result works.
p2 <- p + 
  stat_compare_means(
    comparisons = list(c("Corn", "Prop")),
    method = "t.test",
    label = "p.signif",
    label.y = max(df_MRS$Log10normCFUs, na.rm = TRUE) + 1, size = 7
  )
p2
ggsave("Figures/BuildingBlocks/FigD_CFUs_Corn_vs_Prop_MRS.png", plot = p2, width = 5, height = 2.4, dpi = 300)

##########
# YPG
##########

# Calculate summary stats for error bars (mean ± SD)
summary_df <- df_YPG %>%
  group_by(Diet) %>%
  summarise(
    mean = mean(Log10normCFUs, na.rm = TRUE),
    sd = sd(Log10normCFUs, na.rm = TRUE),
    .groups = 'drop'
  )

# Plot
p<-ggplot(df_YPG, aes(x = Diet, y = Log10normCFUs, fill = Species)) +
  geom_jitter(aes(shape = Species, fill = Species),
              position = position_jitterdodge(jitter.width = 0.3, dodge.width = 0.8),
              size = 2,
              alpha = 0.7,
              color = "black",   # black border to avoid missing warning
              stroke = 0) +      # but no visible border thickness
  scale_fill_manual(values = c(
    "D. melanogaster" = "#1b9e77",
    "D. virilis" = "#d95f02",
    "D. immigrans" = "#7570b3")) +
  scale_shape_manual(values = c(21, 22, 24)) +  # filled shapes that accept fill color
  geom_errorbar(data = summary_df,
                aes(x = Diet, ymin = mean - sd, ymax = mean + sd),
                width = 0.2,
                size = 0.5,
                color = "black",
                inherit.aes = FALSE) +
  geom_segment(data = summary_df,
               aes(x = as.numeric(Diet) - 0.3, xend = as.numeric(Diet) + 0.3,
                   y = mean, yend = mean),
               color = "black",
               size = 1,
               inherit.aes = FALSE) +
  theme_classic() +
  labs(x = NULL, y = "YPG CFUs (Log10)", fill = "Species") +
  theme(axis.title.y = element_text(size = 12),
        legend.position = "left",
        legend.text = element_text(face = "italic"),
        axis.text.x = element_text(size = 12)) +
  scale_y_continuous(limits = c(0,11), breaks = seq(0, 10, by = 2))
# the actual statistical test is above in the Tukey's HSD and lmer(), but t.test here allows the plot to draw an error bar, and the asterisk count is the same anyways.
# Very lazy coding here, but the result works.
p2 <- p + 
  stat_compare_means(
    comparisons = list(c("Corn", "Prop")),
    method = "t.test",
    label = "p.signif",
    label.y = max(df_YPG$Log10normCFUs, na.rm = TRUE) + 1, size = 7
  )
p2
p3 <- p2 + 
  theme(legend.position = "none")
ggsave("Figures/BuildingBlocks/FigD_CFUs_Corn_vs_Prop_YPG.png", plot = p3, width = 2.8, height = 2.4, dpi = 300)

##########
# LB
##########

# Calculate summary stats for error bars (mean ± SD)
summary_df <- df_LB %>%
  group_by(Diet) %>%
  summarise(
    mean = mean(Log10normCFUs, na.rm = TRUE),
    sd = sd(Log10normCFUs, na.rm = TRUE),
    .groups = 'drop'
  )

# Plot
p<-ggplot(df_LB, aes(x = Diet, y = Log10normCFUs, fill = Species)) +
  geom_jitter(aes(shape = Species, fill = Species),
              position = position_jitterdodge(jitter.width = 0.3, dodge.width = 0.8),
              size = 2,
              alpha = 0.7,
              color = "black",   # black border to avoid missing warning
              stroke = 0) +      # but no visible border thickness
  scale_fill_manual(values = c(
    "D. melanogaster" = "#1b9e77",
    "D. virilis" = "#d95f02",
    "D. immigrans" = "#7570b3")) +
  scale_shape_manual(values = c(21, 22, 24)) +  # filled shapes that accept fill color
  geom_errorbar(data = summary_df,
                aes(x = Diet, ymin = mean - sd, ymax = mean + sd),
                width = 0.2,
                size = 0.5,
                color = "black",
                inherit.aes = FALSE) +
  geom_segment(data = summary_df,
               aes(x = as.numeric(Diet) - 0.3, xend = as.numeric(Diet) + 0.3,
                   y = mean, yend = mean),
               color = "black",
               size = 1,
               inherit.aes = FALSE) +
  theme_classic() +
  labs(x = NULL, y = "LB CFUs (Log10)", fill = "Species") +
  theme(axis.title.y = element_text(size = 12),
        legend.position = "left",
        legend.text = element_text(face = "italic"),
        axis.text.x = element_text(size = 12)) +
  scale_y_continuous(limits = c(0,11), breaks = seq(0, 10, by = 2))
# the actual statistical test is above in the Tukey's HSD and lmer(), but t.test here allows the plot to draw an error bar, and the asterisk count is the same anyways.
# Very lazy coding here, but the result works.
p2 <- p + 
  stat_compare_means(
    comparisons = list(c("Corn", "Prop")),
    method = "t.test",
    label = "p.signif",
    label.y = max(df_LB$Log10normCFUs, na.rm = TRUE) + 1, size = 7
  )
p2
p3 <- p2 + 
  theme(legend.position = "none")
ggsave("Figures/BuildingBlocks/FigD_CFUs_Corn_vs_Prop_LB.png", plot = p3, width = 2.8, height = 2.4, dpi = 300)

##########
# BHI
##########

# Calculate summary stats for error bars (mean ± SD)
summary_df <- df_BHI %>%
  group_by(Diet) %>%
  summarise(
    mean = mean(Log10normCFUs, na.rm = TRUE),
    sd = sd(Log10normCFUs, na.rm = TRUE),
    .groups = 'drop'
  )

# Plot
p<-ggplot(df_BHI, aes(x = Diet, y = Log10normCFUs, fill = Species)) +
  geom_jitter(aes(shape = Species, fill = Species),
              position = position_jitterdodge(jitter.width = 0.3, dodge.width = 0.8),
              size = 2,
              alpha = 0.7,
              color = "black",   # black border to avoid missing warning
              stroke = 0) +      # but no visible border thickness
  scale_fill_manual(values = c(
    "D. melanogaster" = "#1b9e77",
    "D. virilis" = "#d95f02",
    "D. immigrans" = "#7570b3")) +
  scale_shape_manual(values = c(21, 22, 24)) +  # filled shapes that accept fill color
  geom_errorbar(data = summary_df,
                aes(x = Diet, ymin = mean - sd, ymax = mean + sd),
                width = 0.2,
                size = 0.5,
                color = "black",
                inherit.aes = FALSE) +
  geom_segment(data = summary_df,
               aes(x = as.numeric(Diet) - 0.3, xend = as.numeric(Diet) + 0.3,
                   y = mean, yend = mean),
               color = "black",
               size = 1,
               inherit.aes = FALSE) +
  theme_classic() +
  labs(x = NULL, y = "BHI CFUs (Log10)", fill = "Species") +
  theme(axis.title.y = element_text(size = 12),
        legend.position = "left",
        legend.text = element_text(face = "italic"),
        axis.text.x = element_text(size = 12)) +
  scale_y_continuous(limits = c(0,11), breaks = seq(0, 10, by = 2))
# the actual statistical test is above in the Tukey's HSD and lmer(), but t.test here allows the plot to draw an error bar, and the asterisk count is the same anyways.
# Very lazy coding here, but the result works.
p2 <- p + 
  stat_compare_means(
    comparisons = list(c("Corn", "Prop")),
    method = "t.test",
    label = "p.signif",
    label.y = max(df_BHI$Log10normCFUs, na.rm = TRUE) + 1, size = 7
  )
p2
p3 <- p2 + 
  theme(legend.position = "none")
ggsave("Figures/BuildingBlocks/FigD_CFUs_Corn_vs_Prop_BHI.png", plot = p3, width = 2.8, height = 2.4, dpi = 300)
```
