## Supplementary material for "Life history and infection susceptibility parameters of *Drosophila* species reared on a common diet": File S2: FigD_Pearson_Corr_MillieSurvs.R

```
library(dplyr)
library(lme4)
library(ggplot2)
library(broom.mixed)

df_Agfp<-read_excel("Data_Raw/Millie_surv_a_gfp.xlsx")
df_Asic<-read_excel("Data_Raw/Millie_surv_a_sic.xlsx")
df_CI<-read_excel("Data_Raw/Millie_surv_c_i.xlsx")

str(df_Agfp)
str(df_Asic)
str(df_CI)

unique(df_Agfp$Experiment)
unique(df_Asic$Experiment)
unique(df_CI$Experiment)

# fix excel date errors
df_Asic <- df_Asic %>%
  mutate(Experiment = ifelse(Experiment == "45843", "2025-22-06", Experiment))

df_CI <- df_CI %>%
  mutate(Experiment = case_when(
    Experiment == "45843" ~ "2025-22-06",
    Experiment == "45784" ~ "2025-13-05",
    TRUE ~ Experiment
  ))

# --- 1. Clean and align data ---------------------------------------

df_Agfp_clean <- df_Agfp %>%
  select(animal, Experiment, rmean) %>%
  filter(!is.na(animal) & !is.na(rmean)) %>%
  mutate(Treatment = "A_GFP")

sum_df_Agfp_clean <- df_Agfp_clean %>%
  group_by(animal) %>%
  summarise(rmean = mean(rmean, na.rm = TRUE))

df_Asic_clean <- df_Asic %>%
  select(animal, Experiment, rmean) %>%
  filter(!is.na(animal) & !is.na(rmean)) %>%
  mutate(Treatment = "A_sicerae")

sum_df_Asic_clean <- df_Asic_clean %>%
  group_by(animal) %>%
  summarise(rmean = mean(rmean, na.rm = TRUE))

df_CI_clean <- df_CI %>%
  select(animal, Experiment, rmean) %>%
  filter(!is.na(animal) & !is.na(rmean)) %>%
  mutate(Treatment = "Clean_Injury")

# --- 2. Pairwise joins ---------------------------------------------

df_gfp_sic <- inner_join(sum_df_Agfp_clean, sum_df_Asic_clean, by = c("animal"), suffix = c("_Agfp", "_Asic"))
df_gfp_ci  <- inner_join(df_Agfp_clean, df_CI_clean,  by = c("animal","Experiment"), suffix = c("_Agfp", "_CI"))
df_sic_ci  <- inner_join(df_Asic_clean, df_CI_clean,  by = c("animal","Experiment"), suffix = c("_Asic", "_CI"))

str(df_gfp_sic)
str(df_gfp_ci)
str(df_sic_ci)

# --- 3. Fit linear models ------------------------------------

# A_GFP vs A_sicerae: different experiment sets → include both as random effects
m_gfp_sic <- lm(
  rmean_Asic ~ rmean_Agfp,
  data = df_gfp_sic)

summary(m_gfp_sic)

# --- Mixed Models --------------------------------------------------------
library(lmerTest)

# A_GFP vs Clean Injury: overlapping experiments → single random effect
m_gfp_ci <- lmer(
  rmean_CI ~ rmean_Agfp + (1 | Experiment),
  data = df_gfp_ci)
VarCorr(m_gfp_ci)
summary(m_gfp_ci)
m_gfp_ci <- lm(rmean_CI ~ rmean_Agfp, data = df_gfp_ci)
summary(m_gfp_ci)

# A_sicerae vs Clean Injury: overlapping experiments → single random effect
m_sic_ci <- lmer(
  rmean_CI ~ rmean_Asic + (1 | Experiment),
  data = df_sic_ci)
VarCorr(m_sic_ci)
summary(m_sic_ci)
m_sic_ci <- lm(rmean_CI ~ rmean_Asic, data = df_sic_ci)
summary(m_sic_ci)


# --- Summarize fixed effects --------------------------------------------

results <- bind_rows(
  tidy(m_gfp_sic, effects = "fixed") %>%
    mutate(Comparison = "A_GFP vs A_sicerae"),
  
  tidy(m_gfp_ci, effects = "fixed") %>%
    mutate(Comparison = "A_GFP vs Clean_Injury"),
  
  tidy(m_sic_ci, effects = "fixed") %>%
    mutate(Comparison = "A_sicerae vs Clean_Injury")
  )

results %>%
  filter(term == "(Intercept)" | grepl("rmean_", term)) %>%
  select(Comparison, term, estimate, std.error, statistic, p.value)

####
##
# reshape the data for plotting purposes
##
####

library(ggplot2)
library(dplyr)
library(broom)

# --- 1. Compute slope and R2 for each comparison ------------------------
# Helper function to extract slope, p-value, and R² using lm
get_stats <- function(df, x, y, comparison) {
  mod <- lm(as.formula(paste(y, "~", x)), data = df)
  tidy_mod <- broom::tidy(mod) %>% filter(term != "(Intercept)")
  glance_mod <- broom::glance(mod)
  
  tibble(
    Comparison = comparison,
    slope = tidy_mod$estimate,
    p_value = tidy_mod$p.value,
    R2 = glance_mod$r.squared
  )
}

stats_list <- bind_rows(
  get_stats(df_gfp_sic, "rmean_Agfp", "rmean_Asic", "A_GFP vs A_sicerae"),
  get_stats(df_gfp_ci, "rmean_Agfp", "rmean_CI", "A_GFP vs Clean_Injury"),
  get_stats(df_sic_ci, "rmean_Asic", "rmean_CI", "A_sicerae vs Clean_Injury")
) %>%
  mutate(label = paste0("_",Comparison, "_<br>","\nslope = ", round(slope, 2),
                        ", R² = ", round(R2, 2)
                        ))

# fix names for plot
stats_list <- stats_list %>%
  mutate(
    label = label %>%
      gsub("A_GFP", "**A. indonesiensis-GFP**", .) %>%
      gsub("A_sicerae", "**A. sicerae**", .) %>%
      gsub("Clean_Injury", "**Clean injury**", .)
  )

stats_list$label <- gsub(" vs", "\n**vs.**", stats_list$label) # gsub() to replace the " vs" in the labels with a next-line hard return 
stats_list$label <- gsub("\n", "<br>", stats_list$label) # gsub() to replace the "\n" in the labels with "<br> per markdown in order to allow bold text
# stats_list$label <- gsub("slope", "_slope", stats_list$label) # gsub() to replace the "\n" in the labels with "<br> per markdown in order to allow bold text
stats_list

# --- 2. Prepare data for plotting --------------------------------------
df_plot <- bind_rows(
  df_gfp_sic %>%
    transmute(
      Comparison = "A_GFP vs A_sicerae",
      animal = animal,
      X = rmean_Agfp,
      Y = rmean_Asic
    ),
  
  df_gfp_ci %>%
    transmute(
      Comparison = "A_GFP vs Clean_Injury",
      animal = animal,
      X = rmean_Agfp,
      Y = rmean_CI,
      Experiment = Experiment
    ),
  
  df_sic_ci %>%
    transmute(
      Comparison = "A_sicerae vs Clean_Injury",
      animal = animal,
      X = rmean_Asic,
      Y = rmean_CI,
      Experiment = Experiment
    )
) %>%
  left_join(stats_list %>% select(Comparison, label), by = "Comparison")

str(df_plot)


# --- 3. Faceted scatter plot with regression lines and R² --------------
library(ggtext)
facet_corr<-ggplot(df_plot, aes(x = X, y = Y, color = animal)) +
  geom_point(size = 2, alpha = 0.2) +
  geom_smooth(method = "lm", se = FALSE, color = "black", linewidth = 0.8) +
  facet_wrap(~label, scales = "fixed") + # label is where the plot titles are stored
  coord_cartesian(xlim = c(1, 15), ylim = c(1, 15)) +
  scale_x_continuous(breaks = seq(1, 15, 2)) +
  scale_y_continuous(breaks = seq(1, 15, 2)) +
  theme_minimal(base_size = 12) +
  labs(
    x = "Mean mortality treatment 2 (days)",
    y = "Mean mortality treatment 1 (days)",
    title = "",
    color = "Animal"
  ) +
  theme(legend.position = "none",
        axis.title = element_text(size=14),
        axis.text = element_text(size = 12),
        strip.text = ggtext::element_markdown(size = 14) # facet label text size using markdown to custom bold just part of the strip text
        # strip.text = element_text(size = 14)   # facet label text size
  )
facet_corr

ggsave("Figures/BuildingBlocks/FigD_Correlations.png", facet_corr, width = 9, height = 4, dpi = 300)
```
