## Supplementary material for "Life history and infection susceptibility parameters of *Drosophila* species reared on a common diet": File S2: FigD_PropCorn.R

```
# load libraries
library(readxl)
library(readr)
library(ggtree)
library(ggplot2)
library(dplyr)
library(patchwork)
library(survminer)
library(survival)


#############
# facet plots
#############
FlyData <- read_excel("Data_Raw/Millie_Corn_vs_Prop_4plots.xlsx")
FlyData$Diet <- FlyData$Media

FlyData$Diet <- recode(FlyData$Diet,
                          "Cornmeal" = "Cam. corn",
                          "Propionic" = "Prop recipe"
                          )

FlyData <- FlyData %>%
  mutate(Strata = interaction(Treatment, Diet))


# rename clean injury and A sicerae to look good
FlyData$Treatment <- recode(FlyData$Treatment,
                            "Clean_Injury" = "Clean injury",
                            "A_sicerae"   = "A. sicerae")

# recode to make the names italics and add superscripts
FlyData$Genotype <- recode(FlyData$Genotype,
                           "D.melanogaster_Isow118(+)_" = "italic('D. melanogaster w')^{1118}",
                           "D.melanogaster_DptBKO_"    = "italic('D. melanogaster DptB')^{KO}",
                           "D.immigrans"                = "italic('D. immigrans')",
                           "D.putrida"                  = "italic('D. putrida')"
                           )
FlyData$Genotype <- factor(FlyData$Genotype, levels = unique(FlyData$Genotype))

# FlySurv <- survfit(Surv(Time2, Censored) ~ Strata + Genotype, data = FlyData)
str(FlyData)
unique(FlyData$Treatment)
unique(FlyData$Genotype)

# subset to remove extra species
FlyData <- FlyData %>%
  filter(Genotype %in% c(
    "italic('D. melanogaster w')^{1118}",
    "italic('D. melanogaster DptB')^{KO}",
    "italic('D. immigrans')",
    "italic('D. putrida')")
    )

# Make sure Treatment and Diet are factors
FlyData$Treatment <- as.factor(FlyData$Treatment)
FlyData$Diet <- as.factor(FlyData$Diet)

# Fit survival
fit <- survfit(Surv(Time2, Censored) ~ Genotype + Treatment + Diet, data = FlyData)

# Extract plot data
plot_data <- ggsurvplot(fit, data = FlyData, ggtheme = theme_bw(), conf.int = FALSE)$plot$data

# Now plot with ggplot2
# p2 <- ggplot(plot_data, aes(x = time, y = surv, color = Diet, linetype = Treatment)) +
#   geom_step() +
#   facet_wrap(~Genotype, scales = "free_y", nrow = 2, ncol = 4,
#              labeller = label_parsed) +  # <-- this tells ggplot to *interpret* superscripts
#   theme_bw() +
#   labs(color = "Diet", linetype = "Treatment", y = "Survival") +
#   scale_x_continuous(breaks = seq(0, 14, by = 2)) +
#   scale_y_continuous(limits = c(0, 1), breaks = seq(0, 1, by = 0.25))
# p2

# Make sure plot_data$Treatment is a factor so we can label it cleanly
plot_data$Treatment <- factor(plot_data$Treatment)

# Create custom labels for the Treatment legend to be italics
treatment_labels <- sapply(levels(plot_data$Treatment), function(x) {
  if (x == "A. sicerae") {
    "italic('A. sicerae')"  # italicize this one
  } else {
    paste0("'", x, "'")     # keep others plain
  }
})

# Define the desired order of Genotypes (in plotmath format)
plot_order <- c(
  "italic('D. melanogaster w')^{1118}",
  "italic('D. melanogaster DptB')^{KO}",
  "italic('D. immigrans')",
  "italic('D. putrida')"
  )

# Apply the order to both FlyData and plot_data
FlyData$Genotype <- factor(FlyData$Genotype, levels = plot_order)
plot_data$Genotype <- factor(plot_data$Genotype, levels = plot_order)

# plot
p2 <- ggplot(plot_data) +
  geom_step(aes(x = time, y = surv, color = Diet, linetype = Treatment), size = 1.2) +
  facet_wrap(~Genotype, scales = "free_y", nrow = 2, ncol = 4,
             labeller = label_parsed) +
  theme_bw() +
  labs(color = "Diet", linetype = "Treatment", y = "Survival", x = "Time (days)") +
  scale_x_continuous(breaks = seq(0, 14, by = 2)) +
  scale_y_continuous(limits = c(0, 1), breaks = seq(0, 1, by = 0.25)) +
  scale_linetype_discrete(labels = parse(text = treatment_labels))
p2 <- p2 +
  theme(
    axis.text       = element_text(size = 14),
    axis.title      = element_text(size = 16),
    strip.text      = element_text(size = 15), # facet label text
    legend.text = element_text(size = 16),
    legend.title = element_text(size = 16)
  )
p2

ggsave("Figures/BuildingBlocks/FigD_cornmeal_facet_surv.png", p2, dpi = 300, width = 14, height = 3.5)

############
# GFP figure
############
GFPresults <- read_excel("Data_Raw/Millie_GFP_Results.xlsx")
GFPresults <- GFPresults %>%
  filter(Day == 5,
         Treatment == "A_GFP")
str(GFPresults)

unique(GFPresults$Genotype)
# recode to make the names italics and add superscripts
GFPresults$Genotype <- recode(
  GFPresults$Genotype,
  "D.putrida"             = "italic('D. putrida')",
  "DptB[KO]"              = "italic('D. melanogaster DptB')^{KO}",
  "iso DrosDel wild-type" = "italic('D. melanogaster w')^{1118}",
  "D.immigrans"           = "italic('D. immigrans')"
)

# Summarize to get means and SEMs for each group
summary_day5 <- GFPresults %>%
  group_by(Genotype, Treatment) %>%
  summarise(
    mean_GFP = mean(GFP_Signal, na.rm = TRUE),
    sem_GFP = sd(GFP_Signal, na.rm = TRUE) / sqrt(n()),
    .groups = "drop"
  )
summary_day5

####################
# Barplot of GFP signal
####################

breaks <- c(1, 2.5, 4)  # numeric values corresponding to colors
colors <- c("firebrick", "grey70", "lightblue")


plot_order <- c(
  "italic('D. melanogaster w')^{1118}",
  "italic('D. melanogaster DptB')^{KO}",
  "italic('D. immigrans')",
  "italic('D. putrida')"
  )
# Summary data
summary_day5$Genotype <- factor(summary_day5$Genotype, levels = plot_order)
# Raw data (for jitter points)
GFPresults$Genotype <- factor(GFPresults$Genotype, levels = plot_order)

###
### Have to subset both summary_day5 and jittered data points because of how these have been added.
###
summary_day5 <- summary_day5 %>%
  filter(Genotype %in% c(
    "italic('D. melanogaster w')^{1118}",
    "italic('D. melanogaster DptB')^{KO}",
    "italic('D. immigrans')",
    "italic('D. putrida')")
  )

GFPresults <- GFPresults %>%
  filter(Genotype %in% c(
    "italic('D. melanogaster w')^{1118}",
    "italic('D. melanogaster DptB')^{KO}",
    "italic('D. immigrans')",
    "italic('D. putrida')")
  )

# # make bold + italics
# GFPresults$Genotype <- gsub(
#   "^italic\\((.+)\\)(\\^\\{.*\\})?$",   # match optional superscript
#   "bold(italic(\\1))\\2",               # wrap the main part in bold+italic, keep superscript
#   GFPresults$Genotype
# )
# GFPresults$Genotype <- factor(GFPresults$Genotype)

# barplot itself with plotmath names
p_bar <- ggplot(summary_day5, aes(x = Genotype, y = mean_GFP, fill = mean_GFP)) +
  geom_col(position = position_dodge(width = 0.9), color = "black") +
  geom_errorbar(
    aes(ymin = mean_GFP - sem_GFP, ymax = mean_GFP + sem_GFP),
    width = 0.2, position = position_dodge(width = 0.9)
  ) +
  geom_jitter(
    data = GFPresults,
    aes(x = Genotype, y = GFP_Signal),
    width = 0.25, alpha = 0.3, height = 0, size = 1.8, show.legend = FALSE,
    inherit.aes = FALSE) +
  labs(x = "", y = "GFP signal") +
  theme_bw(base_size = 12) +
  scale_fill_gradientn(colors = colors, values = scales::rescale(breaks), limits = c(min(breaks), max(breaks))
  ) +
  scale_x_discrete(labels = function(x) parse(text = x)) +
  theme(axis.text.x = element_text(angle = 60, hjust = 1, size = 13),
        panel.grid.major.x = element_blank(), legend.position = "none",
        axis.text.y = element_text(size = 14),
        axis.title.y = element_text(size=14))
p_bar

ggsave("Figures/BuildingBlocks/FigD_GFPsignal.png", p_bar, dpi = 300, width = 5, height = 5)
```
