## Supplementary material for "Life history and infection susceptibility parameters of *Drosophila* species reared on a common diet": File S2: FigD_treeplotversion_unused.R

```
# Read data and tree
FlyData <- read_excel("Data_Raw/Millie_corn_vs_prop.xlsx")
edtree <- read.nexus("Data_Raw/Drosophilidae_tree_25_03_2025_EGRagnoni.tree.nex")

print(FlyData,n=80)

# Add variants to the tree
# Identify the node to split (e.g., D. melanogaster)
target_tip <- "D.melanogaster"
variant1 <- "D.melanogaster_iso_w1118"
variant2 <- "D.melanogaster_DptBKO"

# Add the variants as tips using bind.tip()
edtree <- bind.tip(edtree, tip.label = variant1, edge.length = 0.00, where = which(edtree$tip.label == target_tip))
edtree <- bind.tip(edtree, tip.label = variant2, edge.length = 0.00, where = which(edtree$tip.label == target_tip))

# Prune unused tips
edtree <- drop.tip(edtree, sort(edtree$tip.label[
  as.character(unique(edtree$tip.label)) %in% FlyData$Genotype == FALSE]))


# Plot the updated tree
plot(edtree)

# Create tree plot with ggtree
p_tree <- ggtree(edtree) +
  geom_tiplab(aes(label = paste0("italic('", label, "')")),
              size = 3,
              align = TRUE,
              parse = TRUE) +   # parse expressions like italic()
  theme_tree2() +
  theme(
    axis.text.x  = element_blank(),
    axis.ticks.x = element_blank(),
    axis.title.x = element_blank(),
    axis.line.x  = element_blank(),
    plot.margin  = margin(r = 10)   # optional: extra space on right
  ) +
  coord_cartesian(clip = "off")

tip_labels <- p_tree$data %>%
  filter(isTip) %>%
  arrange(y) %>%   # sort by the y-coordinate to match vertical order in the plot
  pull(label)
tip_labels


### Egg-to-adult Bar plot

# Define order of species (top to bottom in plot)
species_order <- c("D.bifasciata","D.melanogaster_DptBKO","D.melanogaster_iso_w1118","D. immigrans","D.putrida","D.bizonata", "D.multispina", "D.deflecta") # because of coord_flip, this has to be in reverse of the tree itself top to bottom
species_order <- tip_labels

head(FlyData)

# Reorder factor levels accordingly
summary_df <- FlyData %>%
  mutate(Genotype = factor(Genotype, levels = species_order)) %>%
  mutate(mortality = 1-(rmean/14))


# First, summarize mortality per Diet/Bacteria/Genotype across Dates
summary_plot <- summary_df %>%
  group_by(Diet, Bacteria, Genotype) %>%
  summarise(
    mean_mort = mean(mortality, na.rm = TRUE),
    sd_mort   = sd(mortality, na.rm = TRUE),
    n         = n(),
    se_mort   = sd_mort / sqrt(n),
    .groups = "drop"
  )

# create a colour scale
scale_fill_gradient2(
  low = "firebrick",
  mid = "grey70",
  high = "lightblue",
  midpoint = 0.5,
  limits = c(0, 1),
  name = "Mortality"
)

#plot
p_bar <- ggplot(summary_plot, aes(x = Genotype,
                                  y = mean_mort,
                                  fill = mean_mort)) +
  geom_col(width = 0.6) +
  geom_errorbar(aes(ymin = mean_mort - sd_mort,
                    ymax = mean_mort + sd_mort),
                width = 0.2,
                na.rm = TRUE) +
  coord_flip() +
  facet_wrap(~ Bacteria+Diet, ncol = 4, nrow = 1, scales = "free_y")+
  scale_y_continuous(limits = c(-.05, 1), # limit of -.05 to allow error bars to plot
                     breaks = seq(0, 1, by = 0.25)) + 
  scale_fill_gradient2(
    low = "firebrick",
    mid = "grey70",
    high = "lightblue",
    midpoint = 0.5,
    limits = c(0, 1),
    name = "Mortality"
  ) +
  labs(x = "", y = "Mortality (Mean ± StDev)",
       title = "") +
  theme_bw() +
  theme(strip.background = element_rect(fill = "grey90"),
        strip.text = element_text(face = "bold"),
        legend.position = "right",
        axis.text.y = element_text(face = "italic", size = 0))
p_bar


p_combined <- p_tree + p_bar + plot_layout(widths = c(4, 25))& 
  theme(plot.margin = margin(r = 100))  # add right margin to tree
p_combined

# Save as PNG
ggsave(filename = "Figures/z_not_used/Figure_CornmealPropionic_barplot.png", plot = p_combined, width = 14, height = 6, dpi = 300)
```
