## Supplementary material for "Life history and infection susceptibility parameters of *Drosophila* species reared on a common diet": File S2: FigDsupp_PropCorn_fullspecies.R

```
# load libraries
library(readxl)
library(readr)
library(ggtree)
library(ggplot2)
library(dplyr)
library(patchwork)
library(survminer)
library(survival)

#############
# facet plots
#############
FlyData <- read_excel("Data_Raw/Millie_Corn_vs_Prop_4plots.xlsx")
FlyData$Diet <- FlyData$Media

FlyData$Diet <- recode(FlyData$Diet,
                       "Cornmeal" = "Cam. corn",
                       "Propionic" = "Prop recipe"
                       )

FlyData <- FlyData %>%
  mutate(Strata = interaction(Treatment, Diet))

# rename clean injury and A sicerae to look good
FlyData$Treatment <- recode(FlyData$Treatment,
                            "Clean_Injury" = "Clean injury",
                            "A_sicerae"   = "A. sicerae")

# recode to make the names italics and add superscripts
FlyData$Genotype <- recode(FlyData$Genotype,
                           "D.melanogaster_Isow118(+)_" = "italic('D. melanogaster w')^{1118}",
                           "D.melanogaster_DptBKO_"    = "italic('D. melanogaster DptB')^{KO}",
                           "D.bizonata"                 = "italic('D. bizonata')",
                           "D.deflecta"                 = "italic('D. deflecta')",
                           "D.multispina"               = "italic('D. multispina')",
                           "D.immigrans"                = "italic('D. immigrans')",
                           "D.putrida"                  = "italic('D. putrida')",
                           "D.bifasciata"               = "italic('D. bifasciata')"
                           )
FlyData$Genotype <- factor(FlyData$Genotype, levels = unique(FlyData$Genotype))

# FlySurv <- survfit(Surv(Time2, Censored) ~ Strata + Genotype, data = FlyData)
str(FlyData)
unique(FlyData$Treatment)
unique(FlyData$Genotype)

# Make sure Treatment and Diet are factors
FlyData$Treatment <- as.factor(FlyData$Treatment)
FlyData$Diet <- as.factor(FlyData$Diet)

# Fit survival
fit <- survfit(Surv(Time2, Censored) ~ Genotype + Treatment + Diet, data = FlyData)

# Extract plot data
plot_data <- ggsurvplot(fit, data = FlyData, ggtheme = theme_bw(), conf.int = FALSE)$plot$data

# Make sure plot_data$Treatment is a factor so we can label it cleanly
plot_data$Treatment <- factor(plot_data$Treatment)

# Create custom labels for the Treatment legend to be italics
treatment_labels <- sapply(levels(plot_data$Treatment), function(x) {
  if (x == "A. sicerae") {
    "italic('A. sicerae')"  # italicize this one
  } else {
    paste0("'", x, "'")     # keep others plain
  }
})

# Define the desired order of Genotypes (in plotmath format)
plot_order <- c(
  "italic('D. melanogaster w')^{1118}",
  "italic('D. melanogaster DptB')^{KO}",
  "italic('D. bifasciata')",
  "italic('D. immigrans')",
  "italic('D. putrida')",
  "italic('D. bizonata')",
  "italic('D. multispina')",
  "italic('D. deflecta')"
  )

# Apply the order to both FlyData and plot_data
FlyData$Genotype <- factor(FlyData$Genotype, levels = plot_order)
plot_data$Genotype <- factor(plot_data$Genotype, levels = plot_order)

# plot
p2 <- ggplot(plot_data) +
  geom_step(aes(x = time, y = surv, color = Diet, linetype = Treatment), size = 1.2) +
  facet_wrap(~Genotype, scales = "free_y", nrow = 2, ncol = 4,
             labeller = label_parsed) +
  theme_bw() +
  labs(color = "Diet", linetype = "Treatment", y = "Survival", x = "Time (days)") +
  scale_x_continuous(breaks = seq(0, 14, by = 2)) +
  scale_y_continuous(limits = c(0, 1), breaks = seq(0, 1, by = 0.25)) +
  scale_linetype_discrete(labels = parse(text = treatment_labels))
p2 <- p2 +
  theme(
    axis.text       = element_text(size = 10),
    axis.title      = element_text(size = 13),
    strip.text      = element_text(size = 9), # facet label text
    legend.text = element_text(size = 16),
    legend.title = element_text(size = 16)
  )
p2

ggsave("Figures/BuildingBlocks/FigDsupp_cornmeal_facet_surv.png", p2, dpi = 300, width = 10, height = 4)

############
# GFP figure
############
GFPresults <- read_excel("Data_Raw/Millie_GFP_Results.xlsx")
GFPresults <- GFPresults %>%
  filter(Day == 5,
         Treatment == "A_GFP")
str(GFPresults)

unique(GFPresults$Genotype)
# recode to make the names italics and add superscripts
GFPresults$Genotype <- recode(
  GFPresults$Genotype,
  "D.suzukii"             = "italic('D. suzukii')",
  "D.micromelanica"       = "italic('D. micromelanica')",
  "D.phalerata"           = "italic('D. phalerata')",
  "D.putrida"             = "italic('D. putrida')",
  "D.bifasciata"          = "italic('D. bifasciata')",
  "DptB[KO]"              = "italic('D. melanogaster DptB')^{KO}",
  "iso DrosDel wild-type" = "italic('D. melanogaster w')^{1118}",
  "D.ananassae"           = "italic('D. ananassae')",
  "D.immigrans"           = "italic('D. immigrans')"
)

# Summarize to get means and SEMs for each group
summary_day5 <- GFPresults %>%
  group_by(Genotype, Treatment) %>%
  summarise(
    mean_GFP = mean(GFP_Signal, na.rm = TRUE),
    sem_GFP = sd(GFP_Signal, na.rm = TRUE) / sqrt(n()),
    .groups = "drop"
  )
summary_day5

####################
# Barplot
####################

breaks <- c(1, 2.5, 4)  # numeric values corresponding to colors
colors <- c("firebrick", "grey70", "lightblue")


plot_order <- c(
  "italic('D. melanogaster w')^{1118}",
  "italic('D. melanogaster DptB')^{KO}",
  "italic('D. suzukii')",
  "italic('D. ananassae')",
  "italic('D. bifasciata')",
  "italic('D. micromelanica')",
  "italic('D. immigrans')",
  "italic('D. phalerata')",
  "italic('D. putrida')"
  )
# Summary data
summary_day5$Genotype <- factor(summary_day5$Genotype, levels = plot_order)
# Raw data (for jitter points)
GFPresults$Genotype <- factor(GFPresults$Genotype, levels = plot_order)


# barplot itself with plotmath names
p_bar <- ggplot(summary_day5, aes(x = Genotype, y = mean_GFP, fill = mean_GFP)) +
  geom_col(position = position_dodge(width = 0.9), color = "black") +
  geom_errorbar(
    aes(ymin = mean_GFP - sem_GFP, ymax = mean_GFP + sem_GFP),
    width = 0.2, position = position_dodge(width = 0.9)
  ) +
  geom_jitter(
    data = GFPresults,
    aes(x = Genotype, y = GFP_Signal),
    width = 0.25, alpha = 0.3, height = 0, size = 1.8, show.legend = FALSE,
    inherit.aes = FALSE) +
  labs(x = "", y = "GFP signal") +
  theme_bw(base_size = 12) +
  scale_fill_gradientn(colors = colors, values = scales::rescale(breaks), limits = c(min(breaks), max(breaks))
  ) +
  scale_x_discrete(labels = function(x) parse(text = x)) +
  theme(axis.text.x = element_text(angle = 55, hjust = 1, size = 9),
        panel.grid.major.x = element_blank(), legend.position = "none",
        axis.text.y = element_text(size = 14),
        axis.title.y = element_text(size=14))
p_bar

ggsave("Figures/BuildingBlocks/FigDsupp_GFPsignal.png", p_bar, dpi = 300, width = 5, height = 4)


#############
# now a tree
#############

AcetoSpecies <- read.nexus("Data_Raw/Drosophilidae_tree_25_03_2025_EGRagnoni.tree.nex")
AcetoSpecies$tip.label

AcetoSpeciesList <- c(
  "D.ananassae",
  "D.bifasciata",
  "D.bizonata",
  "D.deflecta",
  "D.immigrans",
  "D.melanogaster",
  "D.multispina",
  "D.phalerata",
  "D.putrida",
  "D.suzukii",
  "D.micromelanica"
  ) # define

# Add variants to the tree
# Identify the node to split (e.g., D. melanogaster)
target_tip <- "D.melanogaster"
variant1 <- "D.melanogaster_iso_w1118"
variant2 <- "D.melanogaster_DptBKO"

# Add the variants as tips using bind.tip()
AcetoSpecies <- bind.tip(AcetoSpecies, tip.label = variant1, edge.length = 0.00, where = which(AcetoSpecies$tip.label == target_tip))
AcetoSpecies <- bind.tip(AcetoSpecies, tip.label = variant2, edge.length = 0.00, where = which(AcetoSpecies$tip.label == target_tip))

# Prune unused tips
AcetoSpecies <- drop.tip(AcetoSpecies, sort(AcetoSpecies$tip.label[
  as.character(unique(AcetoSpecies$tip.label)) %in% AcetoSpeciesList == FALSE]))

str(AcetoSpecies)

# Base tree plot
p <- ggtree(AcetoSpecies, layout = "rectangular") + 
  theme_tree2() +
  theme(
    axis.text.x = element_blank(),
    axis.ticks.x = element_blank(),
    axis.line.x = element_blank()
  )

# Set scale bar length
tree_width <- max(p$data$x) - min(p$data$x)
bar_length <- 0.2 * tree_width

# Add truncated scale bar at y = 1 (or any y-coordinate within tree)
AcetoSpecies_tree <- p + 
  geom_segment(aes(x = 0, xend = bar_length, y = 9.7, yend = 9.7),
               color = "black", linewidth = 0.5) +
  annotate("text", x = bar_length/2, y = 10, label = "0.2", size = 3) +
  geom_tiplab(aes(label = label), data = p$data[p$data$isTip, ], size = 3, fontface = "italic") +
  scale_x_continuous(expand = expansion(mult = c(0.05, 0.45))) +
  scale_y_continuous(expand = expansion(mult = c(0.05, 0.15))) +
  scale_y_reverse(expand = expansion(mult = c(0.05, 0.15)))
AcetoSpecies_tree

ggsave("Figures/BuildingBlocks/FigDsupp_AcetoSpecies_tree.png", AcetoSpecies_tree, width = 4, height = 3.5, dpi = 300)


##############
# magick together
##############
library(magick)

# Read the images
tree_img <- image_read("Figures/BuildingBlocks/FigDsupp_AcetoSpecies_tree.png")
gfp_img  <- image_read("Figures/BuildingBlocks/FigDsupp_GFPsignal.png")
facet_img <- image_read("Figures/BuildingBlocks/FigDsupp_cornmeal_facet_surv.png")

# Combine top row: tree (left) + GFP (right)
top_row <- image_append(c(tree_img, gfp_img))

# Combine top row with bottom row (facet)
final_plot <- image_append(c(top_row, facet_img), stack = TRUE)

# Save final combined image
image_write(final_plot, path = "Figures/BuildingBlocks/Combined_FigDsupp.png", format = "png")


### add letters
# Load the image
library(png)
img <- readPNG("Figures/BuildingBlocks/Combined_FigDsupp.png")
img_dim <- dim(img)

# Open PNG device (same size as image)
png("Figures/Fig4supp_MillieData.png", width = img_dim[2], height = img_dim[1], bg = "transparent") # bg = "transparent" prevents the hairline border being drawn alongside the par expression below
par(mar = c(0, 0, 0, 0), xaxs = "i", yaxs = "i") # Remove margins and set exact axis limits

# Plot blank canvas
plot(1, type = "n", xlim = c(0, img_dim[2]), ylim = c(0, img_dim[1]),
     xaxt = 'n', yaxt = 'n', xlab = '', ylab = '', asp = 1)

# Draw the image
rasterImage(img, 0, 0, img_dim[2], img_dim[1])

# Add labels
text(x = 50, y = 2350, labels = "A", cex = 6, font = 2)
text(x = 1200, y = 2350, labels = "B", cex = 6, font = 2)
text(x = 50, y = 1175, labels = "C", cex = 6, font = 2)

# Close device to save
dev.off()
```
