## Supplementary material for "Life history and infection susceptibility parameters of *Drosophila* species reared on a common diet": File S2: FigE_JH10_surv_data.R

```
# load libraries
library(readxl)
library(readr)
library(ape)
library(ggtree)
library(ggplot2)
library(dplyr)
library(patchwork)
library(survminer)
library(survival)
library(phytools)
library(stargazer)
library(tidyverse)
library(lme4)
library(lmerTest)
library(ggpubr)


#############
# Load the data
#############
FlyDataAJ <- read_excel("Data_Raw/Alex_InfectionResults.xlsx") # OD600 = 0.2, 22'C
FlyDataEGR <- read_excel("Data_Raw/Eduardo_Pbur_JH10.xlsx") # let's confirm what Eduardo's 0.1, 0.5 look like, 20'C


# First, let's subset and standardise the names
FlyDataAJ$Experiment <- as.factor(FlyDataAJ$Experiment)
FlyDataAJ$Time2 <- round(as.numeric(FlyDataAJ$Time2),1)
FlyDataAJ$od <- "0.2"
FlyDataAJ$od <- as.numeric(FlyDataAJ$od)
FlyDataAJ <- FlyDataAJ %>%
  mutate(Genotype = recode(Genotype,
                           "isoDrosDelw1118" = "w[1118]",
                           "isoDptAS69R"     = "DptA[S69R]",
                           "isoDptAΔ822"     = "DptA[Δ822]",
                           "isoDptBKO"       = "DptB[KO]",
                           "isoDptSK1"       = "Dpt[SK1]",
                           "RelEzoi"         = "Rel[E20]")) %>%
  filter(Genotype %in% c("w[1118]", "DptA[S69R]", "DptA[Δ822]", 
                         "DptB[KO]", "Dpt[SK1]", "Rel[E20]")) %>%
  mutate(Treatment = recode(Treatment,
                            "Providencia_burhodogranareia" = "P. burhodogranariea str. B",
                            "Providencia_rettgeri" = "P. rettgeri str. Dmel",
                            "Providencia_JH10" = "P. rettgeri str. JH10"))
# rename Alex's genotypes etc...
str(FlyDataAJ)
unique(FlyDataAJ$Time2)


# First, let's subset and standardise the names
FlyDataEGR$Time2 <- round(FlyDataEGR$dia_morte/24,1)
FlyDataEGR$Experiment <- as.factor(FlyDataEGR$date)
FlyDataEGR$Censored <- FlyDataEGR$morte_confirmada
FlyDataEGR$Person <- "Eduardo"
FlyDataEGR$Temp <- FlyDataEGR$temperature
FlyDataEGR$od[FlyDataEGR$od == 0.51] <- 0.50 # just recode the 0.51 entries as 0.50
FlyDataEGR <- FlyDataEGR %>%
  filter(od != 0.1) # get rid of the pilot experiments with 0.1

FlyDataEGR <- FlyDataEGR %>%
  filter(code %in% c("SK1", "AS69R", "A822", 
                         "BKO", "Relish","Relish Iso","WT")) %>%
  mutate(Genotype = recode(code,
                           "WT" = "w[1118]",
                           "SK1" = "Dpt[SK1]",
                           "AS69R" = "DptA[S69R]",
                           "A822" = "DptA[Δ822]",
                           "BKO" = "DptB[KO]",
                           "Relish Iso" = "Rel[E20]",
                           "Relish" = "Rel[E20]")) %>%
  mutate(Treatment = recode(bacteria,
                            "Providencia burhodogranariea" = "P. burhodogranariea str. B",
                            "Providencia JH10" = "P. rettgeri str. Dmel",
                            "Providencia rettgeri" = "P. rettgeri str. JH10",
                            "Control" = "Clean injury"))

# str(FlyDataEGR)
# str(FlyDataAJ)

# Filter data for each treatment and run Cox models for each
# ---
FlyDataCI <- FlyDataEGR %>% 
  filter(Treatment == "Clean injury")
FlyDataCI <- FlyDataCI %>% mutate(Genotype = relevel(factor(Genotype), ref = "w[1118]"))
cox_CI <- coxph(Surv(Time2, Censored) ~ Genotype, data = FlyDataCI)
# ---
FlyDataPbur_B <- FlyDataAJ %>% 
  filter(Treatment == "P. burhodogranariea str. B")
FlyDataPbur_B <- FlyDataPbur_B %>% mutate(Genotype = relevel(factor(Genotype), ref = "w[1118]"))
cox_Pbur_B <- coxph(Surv(Time2, Censored) ~ Genotype, data = FlyDataPbur_B)
# ---
FlyDataPretDmel <- FlyDataAJ %>% 
  filter(Treatment == "P. rettgeri str. Dmel")
FlyDataPretDmel <- FlyDataPretDmel %>% mutate(Genotype = relevel(factor(Genotype), ref = "w[1118]"))
cox_PretDmel <- coxph(Surv(Time2, Censored) ~ Genotype, data = FlyDataPretDmel)
# ---
FlyDataPretJH10 <- FlyDataAJ %>% 
  filter(Treatment == "P. rettgeri str. JH10")
FlyDataPretJH10 <- FlyDataPretJH10 %>% mutate(Genotype = relevel(factor(Genotype), ref = "w[1118]"))
cox_PretJH10 <- coxph(Surv(Time2, Censored) ~ Genotype, data = FlyDataPretJH10)

summary(cox_CI) # EGR data
summary(cox_Pbur_B) # AJ data
summary(cox_PretDmel) # AJ data
summary(cox_PretJH10) # AJ data

sink("Figures/FigE_PretJH10_coxph.txt")
cat("\n====================\n1. P. burhodogranriea\n====================\n\n")
print(summary(cox_Pbur_B))
cat("\n====================\n1. P. rettgeri str Dmel\n====================\n\n")
print(summary(cox_PretDmel))
cat("\n====================\n1. P. rettgeri str JH10\n====================\n\n")
print(summary(cox_PretJH10))
sink()

# Fit the survival model using the interaction
fit <- survfit(Surv(Time2, Censored) ~ Genotype + Treatment, data = FlyDataCI)
# Plot with ggsurvplot_facet
pCI <- ggsurvplot(
  fit,
  data = FlyDataCI,
  legend.title = "Genotype",
  xlim = c(0,3),
  break.time.by = 1,
  xlab = "Time (days)",
  ylab = "Survival probability",
  palette = "Dark2"
)
pCI$plot <- pCI$plot +
  scale_color_discrete(name = "Genotype", 
                       labels = levels(FlyDataCI$Genotype)) +
  theme(legend.position = "left")

# create BuildingBlocks folder
if (!dir.exists("Figures/z_not_used")) {
  dir.create("Figures/z_not_used", recursive = TRUE)
}

ggsave("Figures/z_not_used/FigEsupp_CI_EGR_Data.png", pCI$plot, dpi = 300, width = 5, height = 3)

###
# now for Alex's data
###

FlyDataAJ$Genotype<-as.factor(FlyDataAJ$Genotype)
unique(FlyDataAJ$Treatment)

FlyDataPbur_B <- FlyDataAJ %>% 
  filter(Treatment == "P. burhodogranariea str. B")
FlyDataPbur_B <- FlyDataPbur_B %>% mutate(Genotype = relevel(factor(Genotype), ref = "w[1118]"))
FlyDataPretDmel <- FlyDataAJ %>% 
  filter(Treatment == "P. rettgeri str. Dmel")
FlyDataPretDmel <- FlyDataPretDmel %>% mutate(Genotype = relevel(factor(Genotype), ref = "w[1118]"))
FlyDataPretJH10 <- FlyDataAJ %>% 
  filter(Treatment == "P. rettgeri str. JH10")
FlyDataPretJH10 <- FlyDataPretJH10 %>% mutate(Genotype = relevel(factor(Genotype), ref = "w[1118]"))

# specify labels manually so I can make them nice
legend_labels <- sapply(levels(FlyDataPbur_B$Genotype), function(lab) {
  # Replace [...] with superscript
  lab2 <- gsub("\\[(.*)\\]", "^{\\1}", lab)
  # Replace spaces with * so plotmath concatenates them
  gsub(" ", "*", lab2)
})

# plot Pbur B separately so the legend is on the left
fit <- survfit(Surv(Time2, Censored) ~ Genotype + Treatment, data = FlyDataPbur_B)

# plot it
p <- ggsurvplot(
  fit,
  data = FlyDataPbur_B,
  legend.title = "Genotype",
  xlim = c(0, 3),
  break.time.by = 1,
  xlab = "Time (days)",
  ylab = "Survival probability",
  palette = "Dark2"
)

legend_labels <- paste0("italic(", legend_labels, ")")

# Apply custom legend labels with parse
p$plot <- p$plot +
  scale_color_discrete(
    name = "Genotype",
    labels = parse(text = legend_labels)
  ) +
  theme(legend.position = "left")

# Save plot
outfile <- paste0("Figures/BuildingBlocks/FlyDataPbur_B_FigE_Survival.png")
ggsave(outfile, p$plot, dpi = 300, width = 4.4, height = 3)

# plot survivals
FlyData_names <- c("FlyDataPretDmel","FlyDataPretJH10")
for (data_name in FlyData_names) {
  
  # Get the data object by name
  df <- get(data_name)
  
  # Make and plot survival model
  fit <- survfit(Surv(Time2, Censored) ~ Genotype + Treatment, data = df)
  p <- ggsurvplot(
    fit,
    data = df,
    legend.title = "Genotype",
    xlim = c(0, 3),
    break.time.by = 1,
    xlab = "Time (days)",
    ylab = "Survival probability",
    palette = "Dark2"
  )
  p$plot <- p$plot +
    scale_color_discrete(name = "Genotype",
                         labels = levels(df$Genotype)) +
    theme(legend.position = "none")
  
  # Save plot
  outfile <- paste0("Figures/BuildingBlocks/", data_name, "_FigE_Survival.png")
  ggsave(outfile, p$plot, dpi = 300, width = 3, height = 3)
}


################
# JH10 tree
################
# Read the tree (Nexus format)
JH10_tree <- read.tree("Data_Raw/JH10_ML_100bs.newick")
JH10_tree <- JH10_tree[[1]]

# Base tree plot
p <- ggtree(JH10_tree, layout = "rectangular") +
  scale_y_reverse() +
  geom_text2(aes(subset = (!isTip & !is.na(as.numeric(label))),
                 label = round(as.numeric(label), 0)),
             hjust = 1.2, vjust = -0.3, size = 2.5) +
  theme_tree2() +
  theme(axis.text.x = element_blank(),
        axis.ticks.x = element_blank(),
        axis.line.x = element_blank()) +
  ggtitle("")

# Set scale bar length
tree_width <- max(p$data$x) - min(p$data$x)
bar_length <- 0.2 * tree_width

# Tip to highlight
tip_to_highlight <- 14   # literal tip number

# Plot tree with custom scale bar and tip labels
JH10treeplot <- p +
  geom_segment(aes(x = 0, xend = bar_length, y = 12.5, yend = 12.5),
               color = "black", linewidth = 0.5) +
  annotate("text", x = bar_length/2, y = 12, label = "0.2", size = 4) +
  scale_x_continuous(expand = expansion(mult = c(0., 0.6))) + # give extra room for tip labels
  # All tips except tip #14
  # geom_tiplab(data = p$data[!p$data$isTip | (1:Ntip(JH10_tree) != 14), ],
  #             aes(label = label),
  #             size = 3, parse = TRUE, fontface = "italic") +
  geom_tiplab(
    data = p$data[!p$data$isTip | (1:Ntip(JH10_tree) != 14), ],
    aes(label = gsub("'", "", label)),  # remove all single quotes
    size = 4,
    fontface = "italic"
  )+
  geom_tiplab(
    data = p$data[p$data$isTip & (1:Ntip(JH10_tree) == 14), ],
    aes(x = x + .0022, y = y, label = "Providencia sp. JH10"),  # custom text here
    size = 4,
    color = "red",
    fontface = "bold.italic"
  )
JH10treeplot

# Save to file
ggsave("Figures/BuildingBlocks/JH10_100bs_ML_tree.png", JH10treeplot,
       width = 10, height = 4, dpi = 300)


##############
# stick em together
##############
library(magick)

# Read images
mid_tree <- image_read("Figures/BuildingBlocks/JH10_100bs_ML_tree.png")
bottom1 <- image_read("Figures/BuildingBlocks/FlyDataPbur_B_FigE_Survival.png")
bottom2 <- image_read("Figures/BuildingBlocks/FlyDataPretDmel_FigE_Survival.png")
bottom3 <- image_read("Figures/BuildingBlocks/FlyDataPretJH10_FigE_Survival.png")
# top1 <- image_read("Figures/BuildingBlocks/FigD_CFUs_Corn_vs_Prop.png")
# top2 <- image_read("Figures/BuildingBlocks/FigD_CFUs_Corn_vs_Prop_NoLegend.png")
# top3 <- image_read("Figures/BuildingBlocks/FigD_CFUs_Corn_vs_Prop_NoLegend.png")
# top4 <- image_read("Figures/BuildingBlocks/FigD_CFUs_Corn_vs_Prop_NoLegend.png")

# Combine top images horizontally
bottom_row <- c(bottom1, bottom2, bottom3) %>%
  image_append(stack = FALSE)  # FALSE -> horizontal
top_row <- c(top1,top2,top3,top4) %>%
  image_append(stack = FALSE)  # FALSE -> horizontal

# Create a transparent spacer (adjust height as needed)
spacer_height <- 130  # pixels
spacer <- image_blank(width = image_info(mid_tree)$width, 
                      height = spacer_height, 
                      color = "none")  # transparent

# # Combine top + spacer + top
# final_plot <- c(top_row, spacer, mid_tree, spacer, bottom_row) %>%
#   image_append(stack = TRUE)   # TRUE -> vertical

# Combine top + spacer + top
final_plot <- c(mid_tree, spacer, bottom_row) %>%
  image_append(stack = TRUE)   # TRUE -> vertical

# Save plot collection image
image_write(final_plot, path = "Figures/BuildingBlocks/FigE_Combined.png", format = "png")

library(png)

# Load the image
img <- readPNG("Figures/BuildingBlocks/FigE_Combined.png")
img_dim <- dim(img)

# Open PNG device (same size as image)
png("Figures/Fig5_AlexData_Labeled.png", width = img_dim[2], height = img_dim[1], bg = "transparent") # bg = "transparent" prevents the hairline border being drawn alongside the par expression below
par(mar = c(0, 0, 0, 0), xaxs = "i", yaxs = "i") # Remove margins and set exact axis limits

# Plot blank canvas
plot(1, type = "n", xlim = c(0, img_dim[2]), ylim = c(0, img_dim[1]),
     xaxt = 'n', yaxt = 'n', xlab = '', ylab = '', asp = 1)

# Draw the image
rasterImage(img, 0, 0, img_dim[2], img_dim[1])

# Add labels
# text(x = 450,  y = 2430, labels = "A", cex = 6, font = 2)
# text(x = 1140,  y = 2430, labels = "B", cex = 6, font = 2)
# text(x = 1800, y = 2430, labels = "C", cex = 6, font = 2)
# text(x = 2450, y = 2430, labels = "D", cex = 6, font = 2)
text(x = 70,  y = 2100, labels = "A", cex = 6, font = 2)
text(x = 410,  y = 950, labels = "B", cex = 6, font = 2)
text(x = 1340, y = 950, labels = "C", cex = 6, font = 2)
text(x = 2250, y = 950, labels = "D", cex = 6, font = 2)
text(x = 930, y = 930, labels = "P. burhodogranariea str B", cex = 4, font = 4)
text(x = 1850, y = 930, labels = "P. rettgeri str Dmel", cex = 4, font = 4)
text(x = 2760, y = 930, labels = "P. rettgeri str JH10", cex = 4, font = 4)

# Close device to save
dev.off()
```
