## Supplementary material for "Life history and infection susceptibility parameters of *Drosophila* species reared on a common diet": File S2: FigE_suppAcetoPEN.R

```
# Get bloating stats

FlyData2 <- read_excel("Data_Raw/Figesupp_AcetoPEN_ProvJH10_2024-11-26.xls", sheet = 2)
str(FlyData2)

FlyData2 <- FlyData2 %>%
  mutate(
    Genotype  = as.factor(Genotype),
    Treatment = as.factor(Treatment),
    Sex       = as.factor(Sex),
    Experiment= as.factor(Experiment),
    Person    = as.factor(Person),
    Temp      = as.numeric(Temp),
    Time      = as.numeric(Time),
    Bloat     = as.numeric(Bloat),
    TotalAlive= as.numeric(TotalAlive)
  )

FlyData2 <- FlyData2 %>%
  mutate(
    Genotype_clean = recode(Genotype,
                            "a. Spat53"             = "S. pattersoni",
                            "b. Z.tub 57"           = "Z. tuberculatus",
                            "c. Dmel 19"            = "D. melanogaster^19",
                            "d. Dput 62"            = "D. putrida",
                            "e. Dmel DrosDel L154"  = "D. melanogaster w^1118",
                            "f. DptAS69R"           = "DptA^S69R",
                            "g. DptA822"            = "DptA^Δ822",
                            "h. DptBKO"             = "DptB^KO",
                            "i. DptSK1"             = "Dpt^SK1",
                            "j. RelE20i"            = "Rel^E20"
    )
  )

FlyData2 <- FlyData2 %>%
  mutate(
    Genotype_expr = recode(Genotype,
                           "a. Spat53"             = "italic('S. pattersoni')",
                           "b. Z.tub 57"           = "italic('Z. tuberculatus')",
                           "c. Dmel 19"            = "italic('D. melanogaster') * phantom() ^19",
                           "d. Dput 62"            = "italic('D. putrida')",
                           "e. Dmel DrosDel L154"  = "italic('D. melanogaster w') * phantom() ^'1118'",
                           "f. DptAS69R"           = "italic('DptA') * phantom() ^'S69R'",
                           "g. DptA822"            = "italic('DptA') * phantom() ^'\u0394822'", # Δ
                           "h. DptBKO"             = "italic('DptB') * phantom() ^'KO'",
                           "i. DptSK1"             = "italic('Dpt') * phantom() ^'SK1'",
                           "j. RelE20i"            = "italic('Rel') * phantom() ^'E20'"
    )
  )

unique(FlyData2$Genotype_expr)
unique(FlyData2$Genotype_clean)

# calc percent bloated at a given time
FlyData2$pcBloat <- 100*(FlyData2$Bloat/FlyData2$TotalAlive)

FlyData2$Group <- as.factor(ifelse(FlyData2$Genotype %in% c("a. Spat53", "b. Z.tub 57", "c. Dmel 19", "d. Dput 62"), "Species",  "Diptericin panel"))
FlyData2$Group <- factor(FlyData2$Group, levels = c("Species", "Diptericin panel"))

str(FlyData2)

# First compute summary data so text matches the summarized bars
summary_data <- FlyData2 %>%
  group_by(Genotype_expr, Time, Group) %>%
  summarise(
    mean_pcBloat = mean(pcBloat, na.rm = TRUE),
    mean_TotalAlive = mean(TotalAlive, na.rm = TRUE),
    .groups = "drop"
  )

bloatplot<-ggplot(FlyData2, aes(x = Genotype_expr, y = pcBloat, fill = as.factor(Time))) +
  geom_bar(
    stat = "summary",
    fun = mean,
    position = position_dodge(width = 0.8)
  ) +
  geom_errorbar(
    stat = "summary",
    fun.data = mean_se,
    position = position_dodge(width = 0.8),
    width = 0.2
  ) +
  # ---- Add TotalAlive text here ----
geom_text(
  data = summary_data,
  aes(
    x = Genotype_expr,
    y = -3,  # slightly below the baseline (adjust as needed)
    label = round(mean_TotalAlive, 1),
    group = as.factor(Time)
  ),
  position = position_dodge(width = 0.8),
  vjust = 1,
  size = 3,
  color = "black"
) +
  scale_fill_brewer(palette = "Reds", name = "Time (Days)") +
  facet_grid(
    ~ Group,
    scales = "free_x",
    space  = "free_x"
  ) +
  labs(
    x = "Genotype",
    y = "% Bloat"
  ) +
  theme_bw(base_size = 12) +
  theme(
    axis.text.x = element_text(angle = 45, hjust = 1, vjust = 1),
    strip.text = element_text(face = "bold", size = 13),
    panel.grid.major.x = element_blank(),
    legend.position = "top"
  ) +
  scale_x_discrete(labels = function(x) parse(text = x))

ggsave(filename = "Figures/BuildingBlocks/Fig5supp2_PercentBloat_AcetoPEN.png",  bloatplot, width = 10, height = 5,  dpi = 300)


##############
# magick together
##############
library(magick)

# Read the images
top_img <- image_read("Figures/BuildingBlocks/Fig5supp1_AcetoPEN_pilot.png")
bot_img  <- image_read("Figures/BuildingBlocks/Fig5supp2_PercentBloat_AcetoPEN.png")

# Get the width of the top image
top_width <- image_info(top_img)$width

# Resize bottom image to match top image width, keeping aspect ratio
bot_img <- image_scale(bot_img, paste0(top_width))  # preserves aspect ratio

# Create a transparent spacer (adjust height as needed)
spacer_height <- 100  # pixels
spacer <- image_blank(width = image_info(top_img)$width,
                      height = spacer_height,
                      color = "white") # "none" = transparent

# Combine top row with bottom row (facet)
fig5supp_img <- image_append(c(top_img, spacer, bot_img), stack = TRUE)

# Save final combined image
image_write(fig5supp_img, path = "Figures/BuildingBlocks/Combined_Fig5supp.png", format = "png")


### add letters
# Load the image
library(png)
img <- readPNG("Figures/BuildingBlocks/Combined_Fig5supp.png")
img_dim <- dim(img)

# Open PNG device (same size as image)
png("Figures/Fig5supp_AcetoPENdata.png", width = img_dim[2], height = img_dim[1], bg = "transparent") # bg = "transparent" prevents the hairline border being drawn alongside the par expression below
par(mar = c(0, 0, 0, 0), xaxs = "i", yaxs = "i") # Remove margins and set exact axis limits

# Plot blank canvas
plot(1, type = "n", xlim = c(0, img_dim[2]), ylim = c(0, img_dim[1]),
     xaxt = 'n', yaxt = 'n', xlab = '', ylab = '', asp = 1)

# Draw the image
rasterImage(img, 0, 0, img_dim[2], img_dim[1])

# Add labels
text(x = 50, y = 2600, labels = "A", cex = 8, font = 2)
text(x = 50, y = 1050, labels = "B", cex = 8, font = 2)

# Close device to save
dev.off()
```
