## Supplementary material for "Life history and infection susceptibility parameters of *Drosophila* species reared on a common diet": File S2: setup_load_tree.R

```
# Read the Nexus tree file
tree <- read.nexus("Data_Raw/Drosophilidae_tree_25_03_2025_EGRagnoni.tree.nex")

# Get and display the tip labels (species names)
tip_labels <- tree$tip.label

# Clean the tip labels: add space after first period
tree$tip.label <- gsub("^(\\w+)\\.", "\\1. ", tree$tip.label)
# Fix the known typo: replace "D. teissieri" with correct spelling "D. teisseri"
tree$tip.label <- gsub("^D\\. teissieri$", "D. teisseri", tree$tip.label)

# Check the structure of the tree
str(tree)
```
