## Supplementary material for "Life history and infection susceptibility parameters of *Drosophila* species reared on a common diet": File S2: setup_PackageInstall_n_Load.R

```
# Ensure easypackages is available
if (!requireNamespace("easypackages", quietly = TRUE)) {
  install.packages("easypackages")
}

# Ensure BiocManager is available
if (!requireNamespace("BiocManager", quietly = TRUE)) {
  install.packages("BiocManager")
}

# List of required packages
cran_packages <- c(
  "coda", "MCMCglmm", "knitr", "kableExtra", "gt", "webshot2",
  "ggplot2", "ggpattern", "tidyr", "ape", "readxl", "dplyr", "readr", "patchwork", "viridisLite"
)

bioc_packages <- c(
  "ggtree", "phytools"
)

# Install any missing CRAN packages
missing_cran <- setdiff(cran_packages, rownames(installed.packages()))
if (length(missing_cran) > 0) {
  install.packages(missing_cran)
}

# Install any missing Bioconductor packages
missing_bioc <- setdiff(bioc_packages, rownames(installed.packages()))
if (length(missing_bioc) > 0) {
  BiocManager::install(missing_bioc)
}

# Load everything via easypackages
easypackages::libraries(c(cran_packages, bioc_packages))
```
