## Supplementary figures and images for "Life history and infection susceptibility parameters of *Drosophila* species reared on a common diet"

### Fig1_VialBottle.jpg

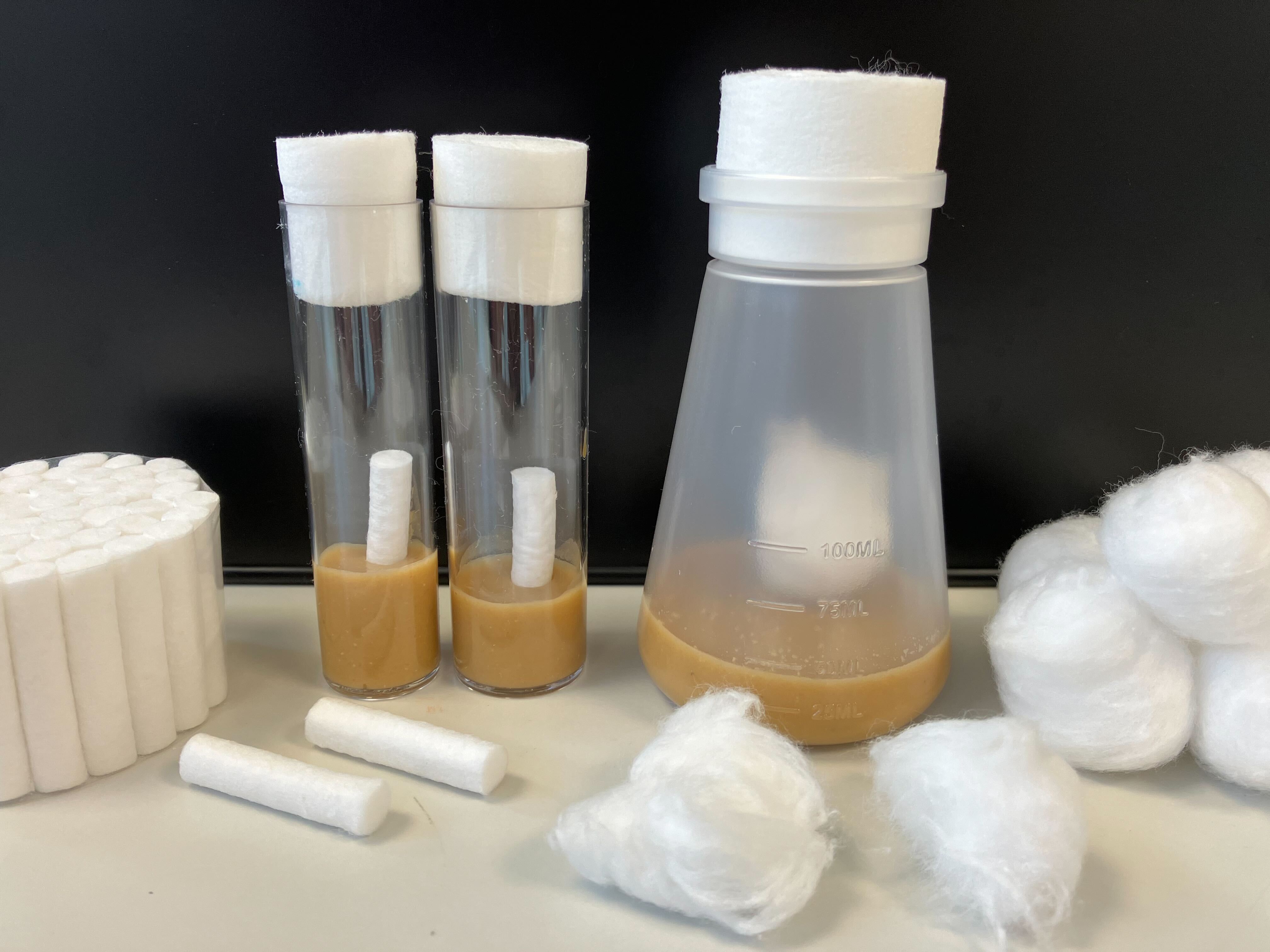

### Fig5supp1_AcetoPEN_pilot.png

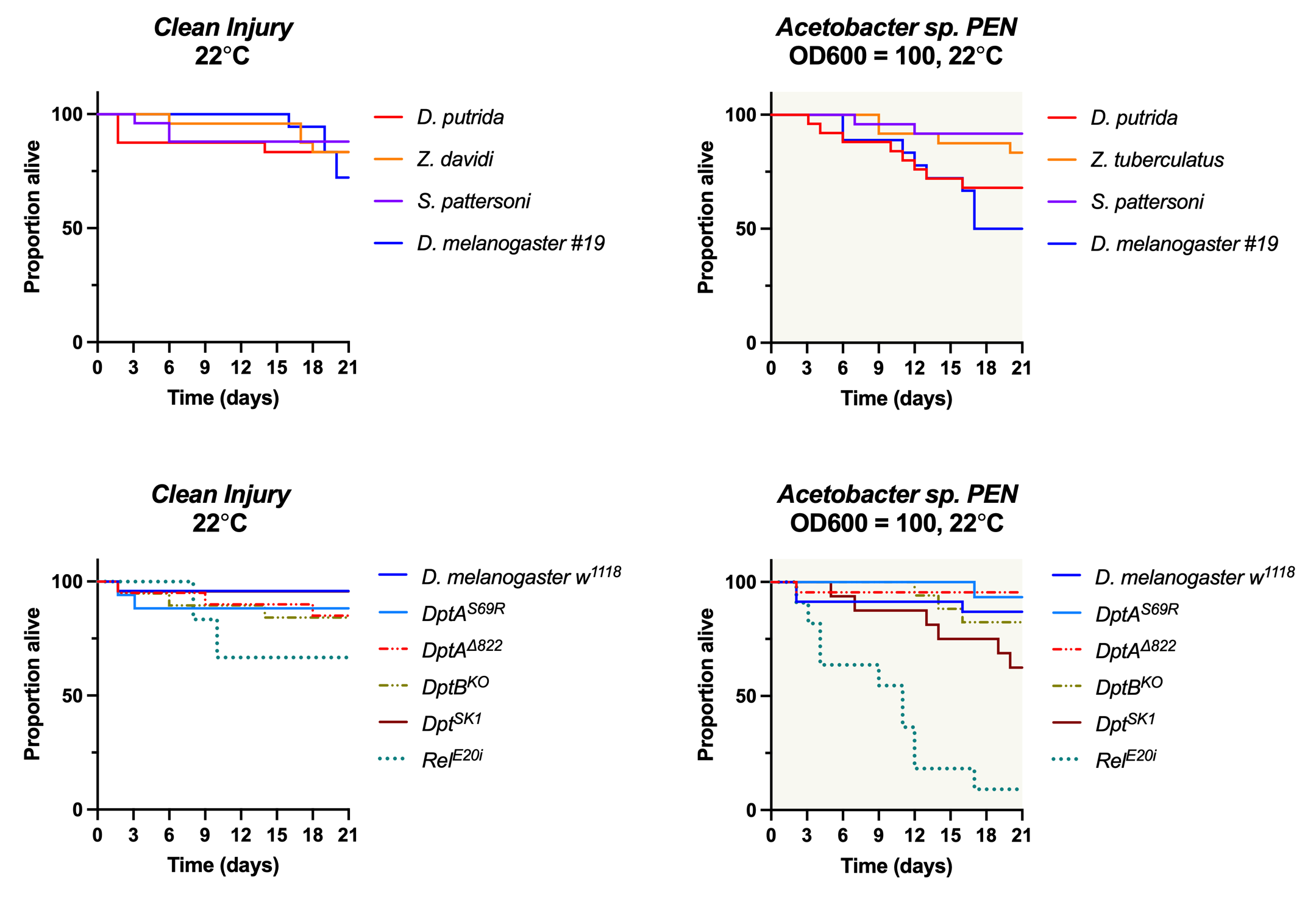

### FloatingLarvae.png

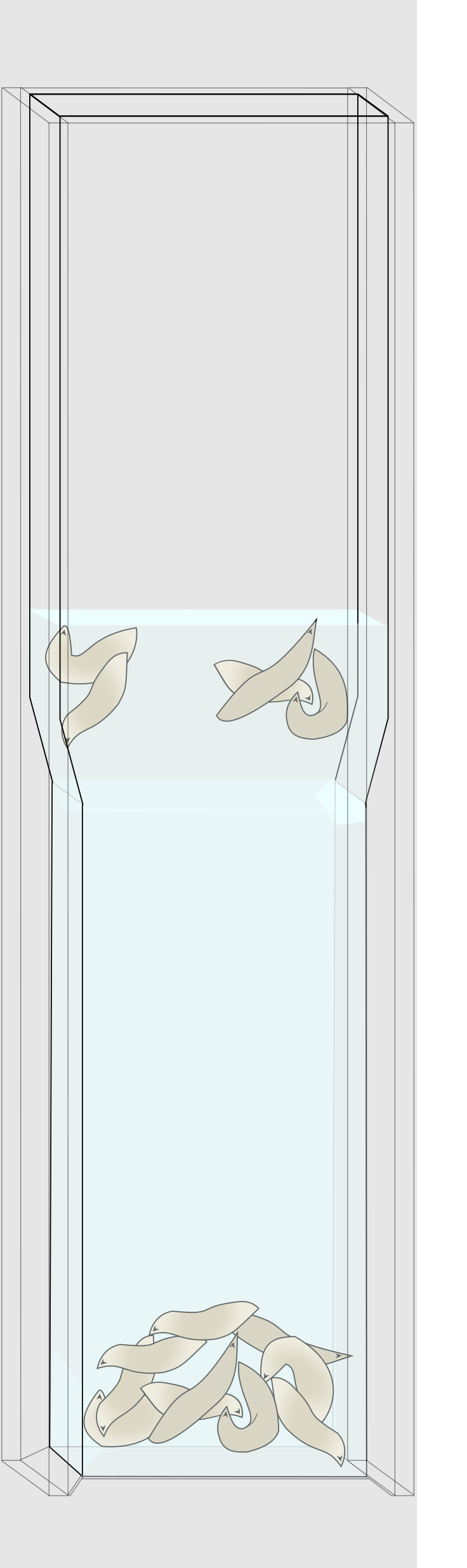
